# Inhibiting LXR*α* Phosphorylation in Hematopoietic Cells Reduces Inflammation and Attenuates Atherosclerosis and Obesity

**DOI:** 10.1101/2020.02.07.939090

**Authors:** Maud Voisin, Elina Shrestha, Claire Rollet, Tatjana Josefs, Tessa J. Barrett, Hye Rim Chang, Rachel Ruoff, Cyrus A. Nikain, Michela L. Garabedian, Emily J. Brown, Ira J. Goldberg, Inés Pineda-Torra, Edward A. Fisher, Michael J. Garabedian

**Author notes:** Corresponding Authors: Michael J. Garabedian, NYU School of Medicine, 450 E. 29th Street, New York, NY 10016, Edward A. Fisher, NYU School of Medicine, SB 705, 435 E. 30^th^ St., New York, NY 10016.

## Abstract

Atherosclerosis and obesity share pathological features including inflammation mediated by innate and adaptive immune cells. LXR*α*, a nuclear receptor, plays a central role in the transcription of inflammatory and lipid metabolic genes. LXR*α* is modulated by phosphorylation at serine 196 (LXR*α* pS196), however, the functional consequences of LXR*α* pS196 in hematopoietic cell precursors in atherosclerosis and obesity have not been investigated. To assess the importance of LXR*α* phosphorylation, bone marrow from LXR*α*WT and S196A mice was transplanted into *Ldlr^-/-^* mice, which were fed a high fat, high cholesterol diet prior to evaluation of atherosclerosis and obesity. Plaques from S196A mice showed reduced inflammatory monocyte recruitment, lipid accumulation, and macrophage proliferation. Expression profiling of CD68+ cells from S196A mouse plaques revealed downregulation of pro-inflammatory genes and upregulation of mitochondrial genes characteristic of anti-inflammatory macrophages. Furthermore, S196A mice had lower body weight and less visceral adipose tissue; this was associated with transcriptional reprograming of the adipose tissue macrophages and resolution of inflammation resulting in less fat accumulation within adipocytes. Thus, reducing LXR*α* pS196 in hematopoietic cells attenuates atherosclerosis and obesity by reprogramming the transcriptional activity of LXR*α* to an anti-inflammatory phenotype.

## INTRODUCTION

LXR*α* is a desmosterol- and oxysterol-activated transcription factor that controls the expression of genes involved in cholesterol homeostasis, apoptosis, inflammation, and cell movement (1, 2). Activation of LXR*α* inhibits atherosclerosis progression in mouse models, which depends in part on LXR*α* activity in macrophages (3, 4). During atherogenesis, immune cells— mostly cholesterol-loaded macrophages accumulate in the arterial wall and form plaques (5–9). Obesity, like atherosclerosis, is associated with lipid accumulation and chronic inflammation (10). Macrophages accumulate in the visceral adipose tissue (VAT) of animals after being fed a high fat diet (11, 12), and produce inflammatory cytokines that contribute to metabolic dysfunction (13).

We and others have shown that LXR*α*’s ability to activate transcription is modulated by phosphorylation at serine 196 (S196 in mouse LXR*α* and S198 in human LXR*α*), which significantly modifies its target gene repertoire (14–17). Whereas LXR*α* pS196 promotes an inflammatory gene signature, unphosphorylated LXR*α* stimulates an anti-inflammatory gene expression profile. Consistent with this idea, in mouse models we observed an increase in LXR*α* pS196 in plaque macrophages under inflammatory conditions associated with atherosclerosis progression, and a decrease in LXR*α* pS196 during the resolution of inflammation in regressing plaques (16, 18). Cholesterol loaded macrophages and hepatocytes also showed increased LXR*α* pS196 (15, 19). Thus, the lack of LXR*α* phosphorylation at S196 appears associated with an anti-inflammatory phenotype.

In this study, we examined the impact of LXR*α* pS196 in immune cells on atherosclerosis progression and obesity, using a bone marrow transplant approach from wild type (WT) and LXR*α* S196A knock-in mice, which are unable to be phosphorylated and should retain only the anti-inflammatory actions of LXR*α*. This approach has the potential to uncover the importance of LXR*α* phosphorylation activities within immune cells, while avoiding confounding effects of LXR*α* S196A expression in other organs. This strategy can also provide insight into potential communication between immune cell types when contrasted with effects on atherosclerosis and obesity from, for example, myeloid-specific LXR*α* S196A expression. Characterizing the role of LXR*α* pS196 in myeloid cells and their contribution to atherosclerosis and obesity has the potential to reveal new therapeutic strategies for both pathologies.

## RESULTS

### LXR*α* S196A promotes an atheroprotective lipoprotein phenotype in a bone marrow transplant model in *Ldlr^-/-^* mice

Given the role that LXR*α* plays in cholesterol homeostasis and control of inflammation, we hypothesized that reducing LXR*α* pS196 would decrease inflammation and reduce atherosclerosis progression. Thus, we used a LXR*α* phosphorylation-deficient knock-in mouse, LXR*α* S196A, in a bone marrow transplantation model (14). This approach allowed us to interrogate the requirement of LXR*α* phosphorylation on the entire cadre of immune cells expressing S196A compared to wild type (WT). LXR*α* S196A or wild type (WT) mice were used as donors for bone marrow transplant into *Ldlr^-/-^* recipients. Following reconstitution, mice were fed a western diet (42% kcal fat and 0.3% kcal cholesterol) for 16 weeks after which time atherogenesis was assessed (Fig. 1A). LXR*α* S196A did not alter the levels of total cholesterol (Fig. 1B). After lipoprotein separation by fast protein liquid chromatography (FPLC), S196A mice showed decreased atherogenic IDL/LDL cholesterol (LDL-C) (20) and increased atheroprotective HDL cholesterol (HDL-C) compared to controls (Fig. 1C). We confirmed this increase by directly measuring HDL-C concentration in the plasma (Fig. S1A). Non-esterified fatty acid (NEFA) and triglyceride (TG) levels in plasma were not changed by LXR*α* S196A (Figs. 1D and E). Thus, LXR*α* S196A mice exhibit an atheroprotective lipoprotein profile with high HDL-C and low LDL-C.

**Figure 1:**
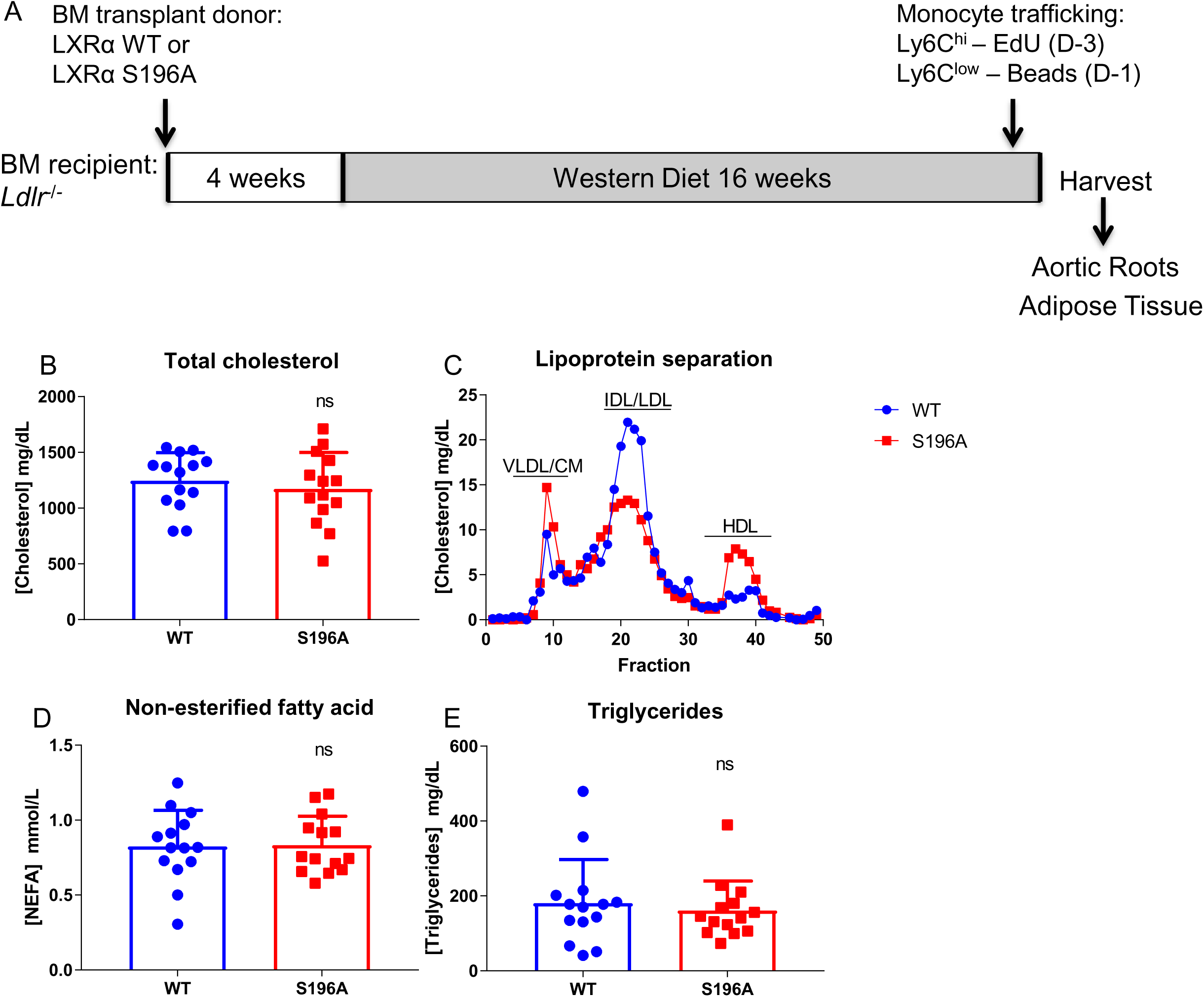
LXRα S196A increases HDL-C and decreases LDL-C levels in plasma. (A) Schematic of the bone marrow (BM) transplant experimental design. (B) Plasma total cholesterol levels from LXRα WT and S196A mice. (C) Plasma lipoproteins distribution as measured by FPLC chromatography between LXRα WT and S196A mice. VLDL: very low-density lipoprotein; CM: Chylomicron; IDL: intermediate-density lipoprotein; LDL: low-density lipoprotein; HDL: high-density lipoprotein. (D) Plasma non-esterified fatty acids and (E) triglycerides concentrations in LXRα WT and S196A mice. Data are expressed as mean ± SD (n=14 per group).

### LXR*α* S196A expressed in bone marrow-derived cells affects multiple kinetic factors governing plaque macrophage content

The favorable lipoprotein profile exhibited by the LXR*α* S196A mice might lead to reduced lipid accumulation within cells of the atherosclerotic plaque. Consistent with this, we found a ∼40 % reduction in the neutral lipid content within the LXR*α* S196A compared to WT plaque as measured by Oil Red O staining (Figs. 2A and S1B). We also measured plaque area and total macrophage content by CD68 staining. We observed trends towards a reduction in total plaque area (Fig. S1C) and CD68+ cell area (Figs. 2B and S1D), whereas necrotic area was unchanged (Fig. 2C). Since plaque macrophage content reflects multiple kinetic factors including monocyte recruitment, macrophage proliferation and apoptosis, we also measured each of these parameters. To determine the Ly6C^high^ monocytes (precursors of pro-inflammatory M1 macrophages in progressing plaques) and Ly6C^low^ monocytes (potential precursors of anti-inflammatory M2 macrophages) content, mice were injected with EdU (5-ethynyl-2’-deoxyuridine) and fluorescent latex beads, 3 days or 1 day before harvest, respectively (21). EdU is incorporated into the DNA of proliferating Ly6C^high^ monocytes while fluorescent beads are engulfed by Ly6C^low^ monocytes. Quantification of EdU- or bead-labeled monocytes infiltrating into the plaques revealed a significant decrease (∼50%) in the recruitment of inflammation-prone Ly6C^high^ monocytes into the plaques of LXR*α* S196A relative to LXR*α* WT mice (Fig. 2D), despite no changes in circulating monocyte number (Figs. S2A-D). In contrast, we did not observe a change in the recruitment of Ly6C^low^ monocytes into the plaque between LXR*α* S196A and WT mice (Fig.2E). Smooth muscle cells in the plaque under high cholesterol can transdifferentiate into macrophage-like CD68+ cells (22), and potentially contribute to the CD68 content. Therefore, we measured the number of cells expressing alpha smooth muscle actin (SMaA), a marker of smooth muscle cells in LXR*α* S196A and WT plaques and no significant difference was observed (Fig. S1E).

**Figure 2:**
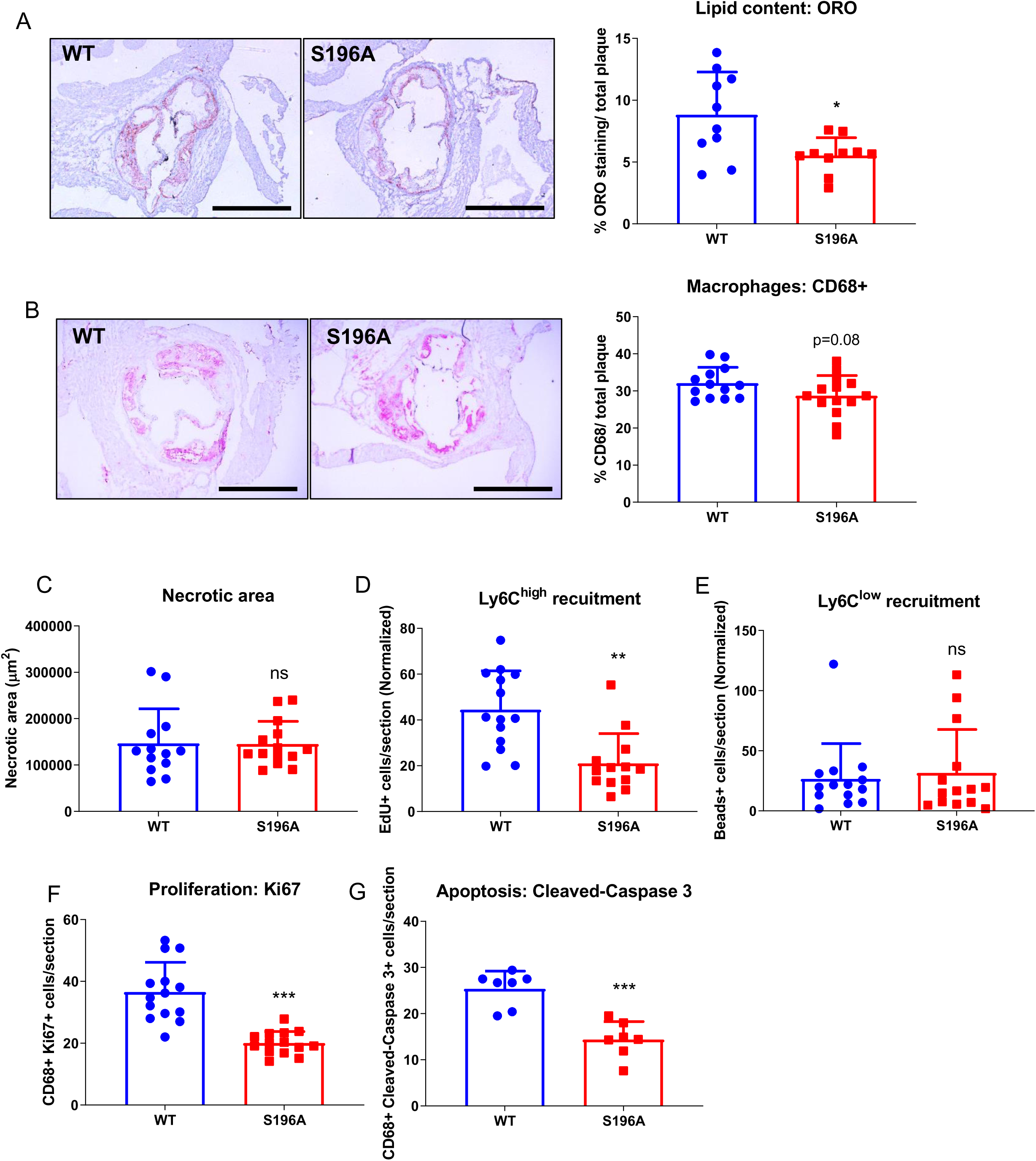
LXRα S196A decreases lipid accumulation in plaque macrophages, reduces monocyte recruitment and macrophage proliferation, as well as decreases macrophage apoptosis in atherosclerotic plaques. (A-B) Representative images and quantification of (A) Oil Red O, (B) CD68 immunostaining, and (C) necrotic area of aortic root sections from LXRα WT and S196A mice. Histological quantification of monocyte recruitment: (D) EdU positive Ly6C^high^ and (E) beads positive Ly6C^low^ cells normalized to the number of positive cells in the blood. (F) Ki67 (proliferation), and (G) Cleaved-Caspase 3 (apoptosis) were measured in the CD68+ cells. Data are expressed as mean ± SD (n=14 except for ORO n=10 and cleaved-Caspase 3 n=7). T test; *P<0.05, **P<0.01, and ***P < 0.001.

We did observe a ∼40% decrease in macrophage proliferation within the plaque in LXR*α* S196A relative to WT by co-staining with the proliferation marker Ki67 and the macrophage marker CD68 (Fig. 2F). Similarly, we observed a ∼40% reduction in apoptosis in LXR*α* S196A compared to WT as measured by cleaved-caspase 3 and CD68 co-staining (Fig. 2G), and would result in macrophage accumulation in the plaque. This could explain why we do not observe a significant reduction in macrophage content in the plaque despite a decrease in both, Ly6C-high monocyte recruitment and macrophage proliferation in S196A plaques. These striking kinetic differences in plaque characteristics between the S196A and WT in the bone marrow transplant model suggests that expression of S196A within myeloid cells is reducing macrophage proliferation, decreasing macrophage recruitment and apoptosis within the plaque microenvironment.

### LXR*α* S196A promotes expression of genes regulating mitochondrial activity and decreases inflammatory pathways in the plaque

To further elucidate the mechanisms governing macrophage inflammatory response in atherosclerosis, we examined gene expression changes from LXR*α* WT and LXR*α* S196A plaque CD68+ cells collected by laser-capture microdissection (23). RNA sequencing (RNA-seq) analysis revealed significant changes in the expression of multiple transcripts. We found that LXR*α* S196A induced 701 and repressed 346 genes in plaque CD68+ cells compared to LXR*α* WT (Fold change > 1.5, p-value < 0.05) (Figs. 3A, S3 and TableS1). Ingenuity Pathway Analysis (IPA) (24) of the genes upregulated by LXR*α* S196A identified mitochondrial function and oxidative phosphorylation (Oxphos) as the top enriched pathways, followed by sirtuin signaling, the TCA cycle and fatty acid oxidation. This suggests that mitochondrial activity and energy metabolism, characteristics of metabolic pathways upregulated in anti-inflammatory macrophages, are induced within the LXR*α* S196A CD68+ cells relative to WT (Fig. 3B-C) (25, 26). Genes downregulated by S196A were associated with inflammatory pathways including phagocytosis in macrophages, nuclear factor of activated T cells (NFAT), Th2 and NFκB signaling (Fig. 3B). Transcription factor analysis via IPA revealed that the majority of the genes upregulated in LXR*α* S196A compared to WT CD68+ cells are associated with PGC1α (PPARGC1A), and to lesser extent by TP53, ESRRα, NUR77 (NR4A1) and mineralocorticoid receptor (NR3C2) (Fig. 3D). In support of skewing macrophages to an anti-inflammatory phenotype, PGC1*α* is a co-activator of LXR*α* (27) and a regulator of mitochondrial biogenesis and function (28, 29). PGC1*α* is activated by deacetylation through sirtuins (30), and this may be strengthened through increased sirtuin signaling identified by IAP in LXR*α* S196A. Additionally, ESRRα is required for mitochondrial biogenesis and activation of mitochondrial genes (31). NUR77 and TP53 control cell cycle and cell death processes (32, 33). This is consistent with decreased proliferation and apoptosis of macrophages in the plaques expressing LXR*α* S196A. Genes downregulated by LXR*α* S196A compared to WT are enriched for members of the signal transducer and activator of transcription family (STAT6, STAT1, STAT3), known to regulate gene expression (Fig. 3D) (34–37). Thus, CD68+ cells within the plaque expressing LXR*α* S196A exhibited an increase in the expression of genes involved in mitochondrial metabolic activity and decreased in genes associated with inflammation, consistent with a more anti-inflammatory macrophage phenotype.

**Figure 3:**
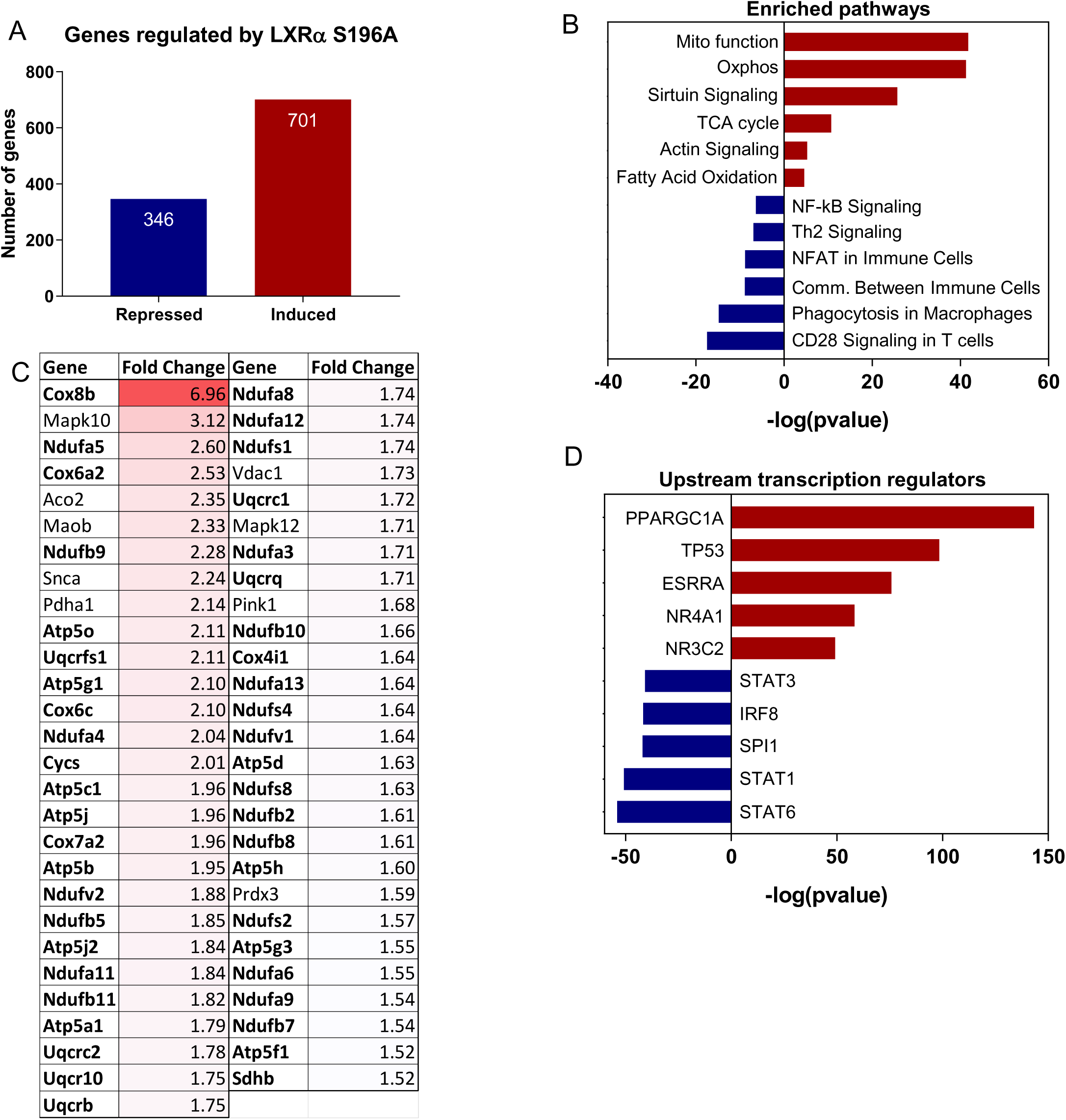
LXR*α* S196A promotes expression of genes regulating mitochondrial activity and decreases inflammatory pathways in plaque macrophages. (A) Number of genes regulated by LXRα S196A versus WT by RNAseq from plaque laser captured microdissected CD68+ cells (Fold change > 1.5, p-value < 0.05). (B) Enriched pathways in S196A after differential gene expression (DGE) analysis using Ingenuity Pathway Analysis (IPA). (D) Genes and fold change of upregulated mitochondrial function-associated genes. Genes in bold are involved in mitochondrial oxidative phosphorylation pathway. (D) Transcriptional regulators associate with upregulated or downregulated genes were determined by IPA transcription factor analysis.

### LXR*α* S196A increases mitochondrial respiration, ATP production and mitochondrial biogenesis in M1 and M2 BMDMs

Given the relationship of LXR*α* S196A to altered expression of genes in plaque CD68+ cells involved in mitochondrial oxidative phosphorylation, we assesses whether LXR*α* S196A enhanced mitochondrial respiration. In bone marrow derived macrophages (BMDM) derived from LXR*α* WT and LXR*α* S196A, we determined the effect of LXR*α* phosphorylation on mitochondrial function by Seahorse XF Extracellular Flux analysis of the oxygen consumption rate (OCR) in BMDM under pro- and anti-inflammatory conditions. In M1 and M2 BMDMs LXR*α* S196A showed higher basal respiration, ATP production and maximal respiration rate compared to control cells (Fig. 4A), suggesting that reduction of LXR*α* phosphorylation promotes electron transport chain capacity in the mitochondria.

**Figure 4:**
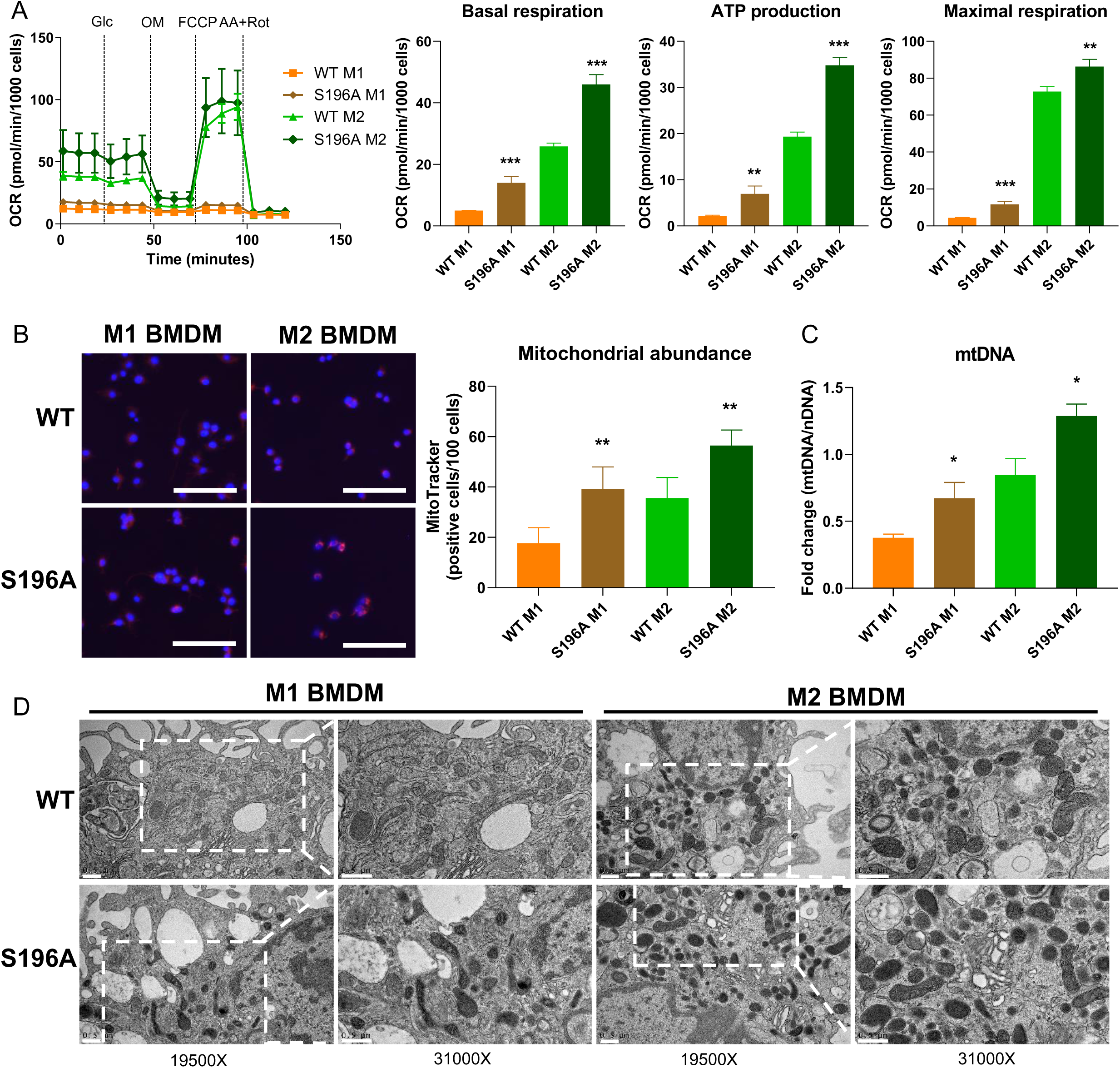
LXRα S196A BMDMs increase mitochondrial abundance and activity. (A) Oxygen consumption rate (OCR) after sequential injection of glucose (Glc), oligomycin (OM), carbonyl cyanide 4-trifluoromethoxyphenyl hydrazine (FCCP), and antimycin plus rotenone (AA+Rot) in LXRα S196A versus WT BMDMs polarized to M1 and M2. OXPHOS parameters: basal respiration, ATP production and maximal respiration, derived from OCR values. (B) Cell imaging of mitochondrial abundance was visualized with MitoTracker Red and quantified (scale bars: 100µm). (C) Mitochondrial abundance in LXRα WT and S196A BMDMs determined by the expression of mitochondrial (mt) DNA normalized to nuclear (n) DNA. (D) Transmission electron micrographs of LXRα S196A versus WT BMDMs polarized to M1 and M2 (scale bars: 0.5 µm). Magnification at 19,500X and 31,000X is shown. Data are expressed as mean ± SD (n=3). T test; *P<0.05, **P<0.01 and ***P<0.001.

One possibility for this could be by increasing mitochondria biogenesis in LXR*α* S196A cells. To examine this, we quantified mitochondrial abundance in LXR*α* WT and S196A BMDMs by labeling the mitochondria within live cells using the mitochondrial-specific probe, MitoTracker (38). Consistent with the enhanced mitochondrial activity in LXR*α* S196A, we found increased MitoTracker positive cells and an increase in mitochondrial (mt) DNA in M1 and M2 LXR*α* S196A cells compared to LXR*α* WT (Fig. 4B-C).

Since LXR*α* S196A gene expression in plaque CD68+ cells converge on the mitochondrion, we evaluated mitochondrial morphology, which is crucial for the maintenance of the mitochondrial membrane potential and ATP production (39) by performing transmission electron microscopy of LXR*α* WT and S196A M1 and M2 BMDMs (Fig. 4D). We observed alterations in mitochondrial morphology and ultrastructural changes in M1 LXR*α* WT BMDMs (less electron dense cristae) as compared with M1 LXR*α* S196A BMDMs (more electron dense cristae). The mitochondria observed in M1 LXR*α* S196A BMDMs look similar to mitochondria found in M2 BMDMs from both WT and LXR*α* S196A. This is consistent with the mitochondria from M1 LXR*α* S196A producing ATP through oxidative phosphorylation (Fig 4A), a mechanism that is normally found in the mitochondria of anti-inflammatory macrophages (40). Collectively, these data demonstrate that reduction of LXR*α* phosphorylation drives mitochondrial respiration processes that are akin to an anti-inflammatory metabolic response likely through its modulation of multiple mitochondrial genes (Fig. 3B), and enhanced mitochondrial biogenesis (Figs. 4B-C).

### LXR*α* S196A protects from diet-induced obesity

Obesity and atherosclerosis share pathological features including lipid accumulation, inflammation and cytokine production mediated by both innate and adaptive immune cells that accumulate within their respective tissues (10, 41–43). We therefore examined whether LXR*α* S196A affected fat deposition and immune cell recruitment in adipose tissue in the same mice as those analyzed for atherosclerosis progression.

Remarkably, LXR*α* S196A mice showed less weight gain during the experiment (Fig. S4A). They weighed 15% less than WT mice after 16 weeks of western diet feeding (Fig. 5A) and had equivalent food intake during the course of the experiment (Fig. S4B). This was observed in two independent cohorts (not shown). We confirmed that total body fat mass was lower (8.2 g vs 10.8 g) in LXR*α* S196A compared to control WT mice by Dual energy X-ray absorptiometry (DEXA) scanning (Fig. 5B). No differences were observed in bone mineral density and lean body mass between LXR*α* S196A and LXR*α* WT mice (Fig. S4C). LXR*α* S196A mice relative to WT also showed reduced VAT, in particular perigonadal white adipose tissue mass (pWAT: 1.32 g vs 1.75 g) (Fig. 5C). This was associated with a decrease in adipocyte size in LXR*α* S196A compared to WT mice (2471 vs 3570 µm^2^) (Figs. 5D-E), consistent with less lipid accumulation. The number of macrophages determined by F4/80+ cell staining was similar in S196A compared to WT pWAT (Figs. S4D-E), although a trend toward a decrease in the number of crown-like structures, a hallmark of the pro-inflammatory processes in adipose tissue (11, 44), was observed (Fig. S4F).

**Figure 5:**
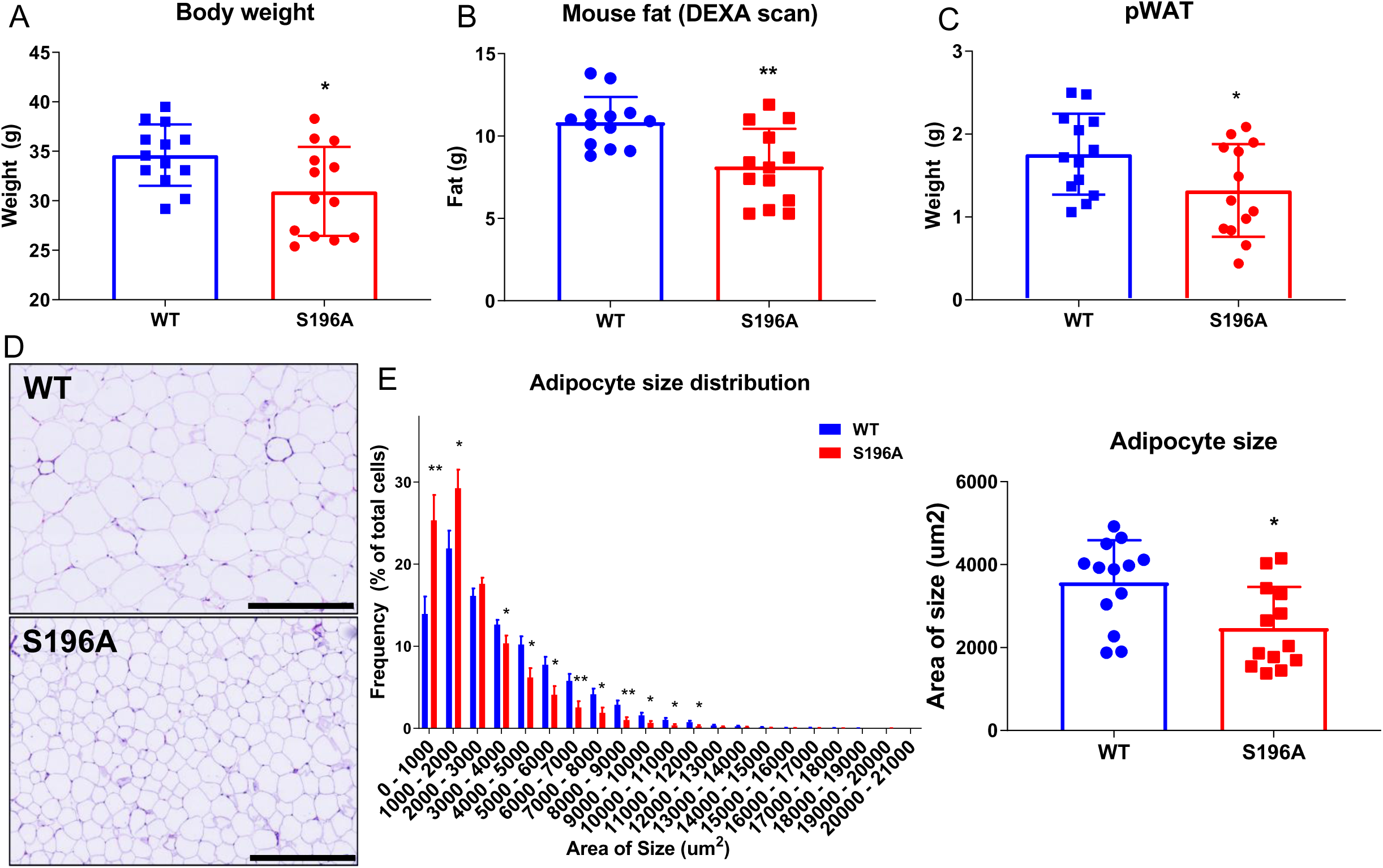
LXRα S196A reduces body weight and fat accumulation in mice. (A) Body weight of LXRα WT and S196A mice. (B) The total body fat was measured by DEXA scan. (C) The perigonadal white adipose tissue (pWAT) was collected and weighed. (D) Representative images of hematoxylin and eosin stained pWAT from LXRα WT and S196A mice. (E) Quantification of the frequency of adipocytes at indicated size ranges and adipocyte size in LXRα WT and S196A mice. Scale bars: 400µm. Data are expressed as mean ± SD (n=14 per group). T test; *P<0.05 and **P<0.01.

A possibility for the adipocyte phenotype is that the myeloid cells from the LXR*α* S196A signal to the adipocytes and decrease the expression of genes involved in lipid accumulation or increase the expression of genes involved in lipolysis. We therefore determined expression in adipocytes from mice reconstituted with bone marrow from either WT or S196A of *Fabp4* (Fatty acid binding protein 4), which promotes fatty acid uptake, and *Plin2* (Perilipin2), which coat intracellular lipid droplets to develop and maintain adipose tissue. We also examined the expression in adipocytes of lipolysis activators *Atgl* (Adipose triglyceride lipase) and *Hsl* (Hormone-sensitive lipase), which hydrolyze stored triglycerides in adipose tissue. The expression in adipocytes of none of these gene were affected in LXR*α* S196A compared to WT mice (Fig. S5A-B). We also determined the expression *Adipoq* (adiponectin), a hormone secreted by adipocytes (adipokine) with anti-inflammatory activities that regulate energy homeostasis, glucose and lipid metabolism. Although the expression of *Adipoq* mRNA did not differ significantly in the adipocytes between WT and S196A, we did find the level of adiponectin protein to be higher in the plasma of LXR*α* S196A compare to its wild type counterpart (Fig. S5C). This is consistent with the plasma levels of adiponectin being higher in lean versus obese rodent and humans (45, 46).

Given that brown adipose tissue (BAT) has the capacity to control energy homeostasis (47), we also examined whether BAT morphology and UCP1 levels were affected in LXR*α* S196A relative to LXR*α* WT mice. BAT tissue morphology and UCP1 protein abundance by immunohistochemistry (IHC) were unchanged in S196A versus WT mice on western diet (Fig. S5D). This suggests that BAT function is not contributing to the lower adipose deposition observed in LXR*α* S196A compared to WT. Also, liver weight was reduced in LXR*α* S196A compared to WT (Fig. S5B), but hepatic lipid droplet accumulation was comparable in LXR*α* S196A and WT mice (Figs. S5F-G). These data demonstrate that LXR*α* S196A expressed in the bone marrow protects *Ldlr^-/-^* mice from diet-induced obesity when challenged by a western diet.

### LXR*α* S196A ATMs are less inflammatory

Analysis of the immune cells present in the pWAT by flow cytometry revealed LXR*α* S196A affects various immune cell populations (Fig. S6A). Adipocyte tissue macrophages (ATMs) were decreased in S196A compared to WT (Fig. 6A). The number of pro-inflammatory FBC macrophages (M1-like: F4/80+CD11b+CD11c+) was reduced (Figs. 6B-C), while the number of anti-inflammatory FB macrophages (M2-like: F4/80+CD11b+CD11c-) was not changed (Fig. 6D). This results in a higher ratio of inflammation resolving FB *versus* inflammation promoting FBC macrophages in LXR*α* S196A compared to LXR*α* WT adipose tissue (Fig. 6E). Analysis of the percentage of dendritic cells (DC) and T cell populations did not demonstrate any significant changes between WT and LXR*α* S196A (Fig. S6B-G). Therefore, the LXR*α* S196A expressing mice resulted in fewer inflammatory ATM in pWAT, characteristic of a leaner phenotype.

**Figure 6:**
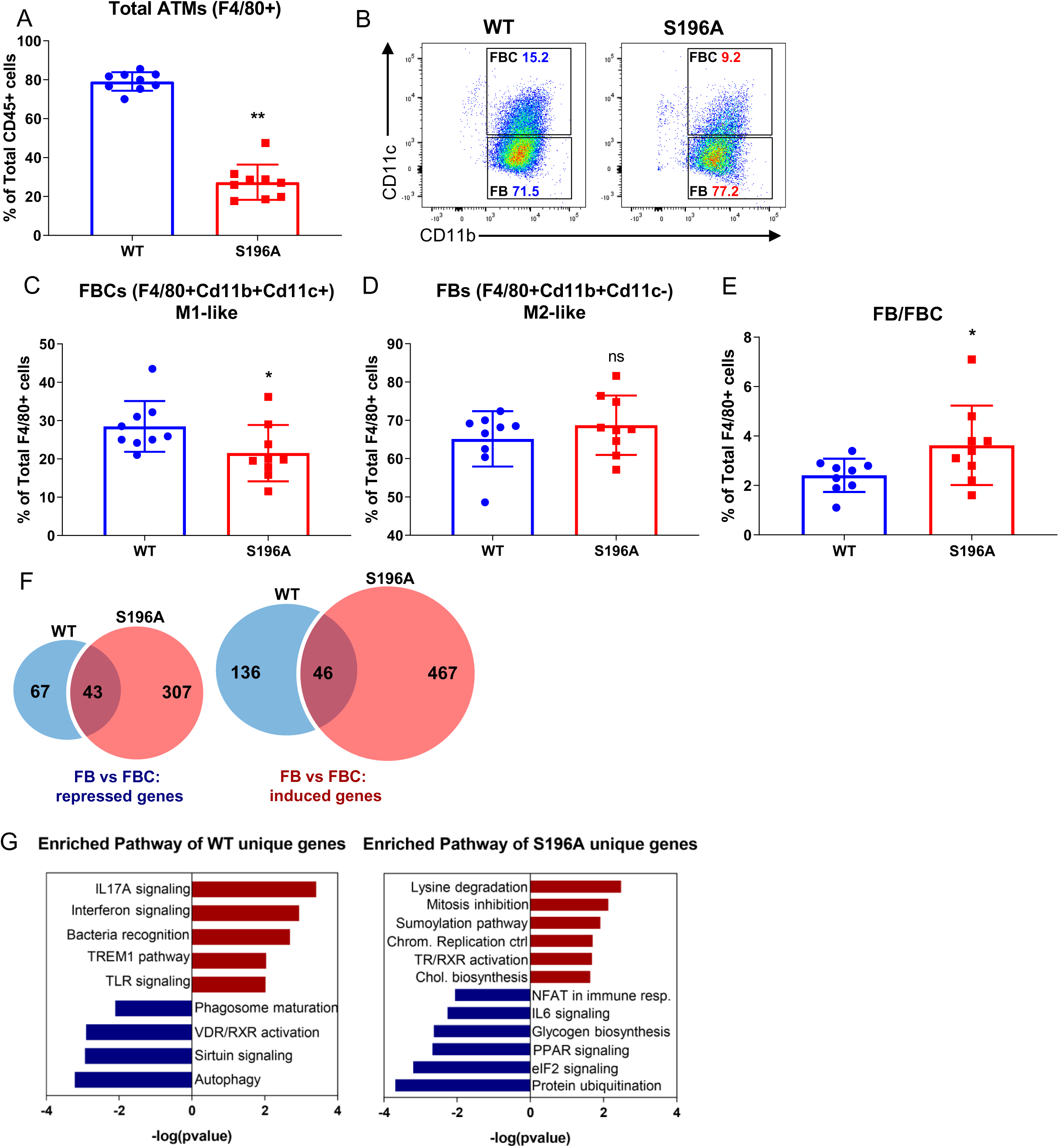
LXRα S196A ATMs are less inflammatory, and regulate pathways involved in metabolism and transcription. Immune cells in pWAT from LXRα WT and S196A were analyzed by flow cytometry: (A) Adipose tissue macrophages (ATM), (B) FBC and FB populations, (C) FBC (F4/80^+^, CD11b^+^, CD11c^+^) as a percentage of F4/80+ cells, and (D) FB (F4/80^+^, CD11b^+^, CD11c^-^) as a percentage of F4/80+ cells, and (E) ratio FBC/FB. (F-G) RNA-seq analysis of FB versus FBC from LXRα WT and S196A mice. (F) Number of genes repressed and induced in FB compared to FBC in LXRα WT and S196A (Fold Change >1.5; p<0.05). (G) Enriched pathways of genes differentially expressed in LXRα WT and S196A in the FB versus FBC. Data are expressed as mean ± SD (n=9 for flow cytometry data: the samples were all pooled for the RNA-seq). T test; *P<0.05 and **P<0.01.

To explore pathways underlying the changes observed in the ATM phenotype in LXR*α* S196A, FB and FBC were sorted from WT and S196A, and RNA-seq analysis was performed. We first analyzed genes in LXR*α* WT and LXR*α* S196A that were differentially expressed in FB *versus* FBC (Fold change > 1.5, p-value < 0.05). We then determined which genes were differentially induced or repressed between LXR*α* WT or LXR*α* S196A (Fig. 6F and Table S2) followed by IPA (Fig. 6-G). Enriched pathways in genes preferentially induced by LXR*α* WT compared to S196A in ATMs include pro-inflammatory factors, such as IL17A and interferon, TREM1 and TLR signaling pathways. In particular, RelA (p65 subunit of NF*κ*B) mRNA was 2.3 fold higher in WT versus S196A in ATMs (Table S3). Pathways associated with genes upregulated in LXR*α* S196A compared to WT include lysine degradation, with *Aldh7a1* (Aldehyde Dehydrogenase 7 Family Member A1) upregulated 8.4 fold (Table S3). ALDH7A1 protects cells from oxidative stress by metabolizing lipid peroxidation-derived aldehydes (48). Additional enriched pathways from genes preferentially induced in LXR*α* S196A ATMs include, sumoylation, in particular *Senp7* [SUMO-specific proteases 7; upregulated 5.8-fold in S196A versus WT (Table S3) and involved in the deconjugation of Sumo moieties (49)], a key determinant of transrepression by LXR*α* (50), and chromatin pathways, which have important ties to macrophage activation in atherosclerotic plaques (51) as well as lipid metabolism and cell cycle inhibitory pathways. An increase in the cell cycle inhibitory pathway in LXR*α* S196A would suggest decreased macrophage proliferation. Pathways downregulated in LXR*α* S196A include inflammatory processes (NFAT and IL6 signaling) (Fig. 6G and Table S3). Together, these data suggest that reduction of pS196 in ATMs decreases inflammatory mediated signaling to adipocytes. This could explain the decrease in lipid accumulation and in adipocyte hypertrophy in pWAT, leading to the reduction in obesity in LXR*α* S196A compared to WT mice.

### Conditioned media from LXR*α* S196A BMDMs reduces lipid accumulation in 3T3-L1 cells

Our previous results strongly suggest that signals from immune cells (most likely ATMs) to adipocytes are capable of reducing lipid accumulation in pWAT in mice reconstituted with LXR*α* S196A compared to WT (Fig. 5C-E). To test this, we developed an *in vitro* assay whereby 3T3-L1 cells are differentiated into adipocytes, treated for 24 hours with filtered conditioned media (CM) from either LXR*α* WT or S196A BMDMs under different inflammatory states (M0, M1 or M2), and stained for neutral lipids by Oil Red O (ORO) to determine lipid accumulation (Fig. 7A). We found that conditioned media from M1 WT BMDMs promoted more lipid accumulation in adipocytes than conditioned media from M1 S196A BMDMs, whereas conditioned media from M0 and M2 from either WT or S196A BMDMs showed no differences in lipid accumulation (Fig. 7B). This suggests M1 S196A cells secrete differing factors than M1 WT macrophages; a decrease in pro-inflammatory and/or an increase in anti-inflammatory factors, that promotes lipid accumulation.

**Figure 7:**
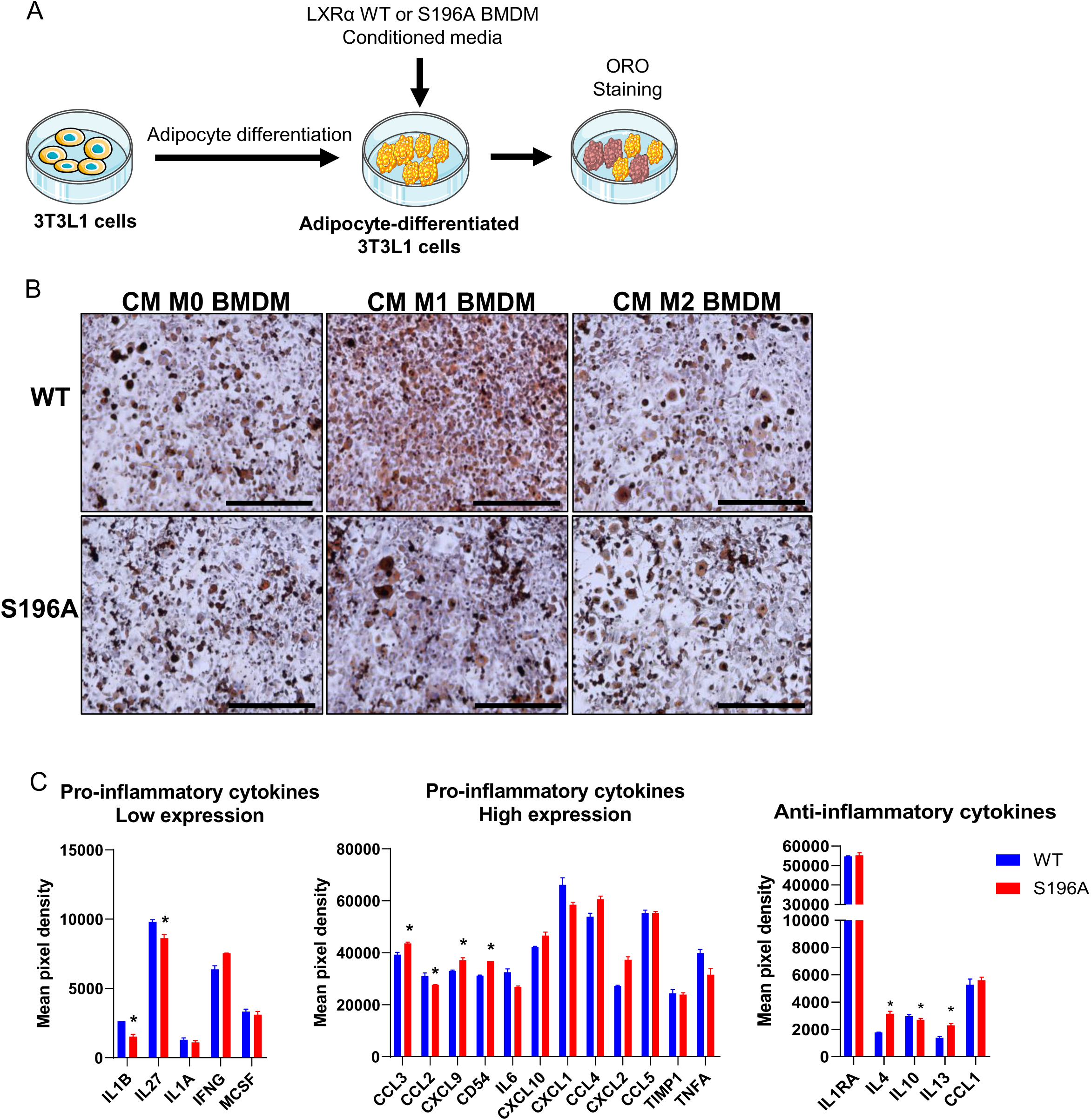
LXRα S196A BMDMs alter lipid accumulation of 3T3-L1 due to modulation of secreted-cytokines. (A) Experimental design: 3T3-L1 cells were differentiated into adipocytes and then treated with conditioned media from WT or S196A BMDMs (M0, M1, or M2) for 24h and neutral lipid accumulation was determined by Oil Red O (ORO) staining. (B) Shown are representative images of ORO staining of differentiated 3T3-L1 adipocytes treated with M0, M1 or M2 conditioned media from LXRα WT or S196A BMDMs. (C) Pro-inflammatory (low and high expression) and anti-inflammatory cytokines profiles of M1 WT and S196A BMDM. Scale bars: 400µm. Data are expressed as mean ± SD (n=3). T test; *P<0.05.

We therefore compared the level of cytokines in the media of M1 LXR*α* WT and S196A BMDMs using a cytokine array. We found in S196A relative to WT conditioned media a significant decrease in the pro-inflammatory cytokines IL1β, IL27 and CXCL2, as well as an increase in CCL3, CXCL9, and CD54 (Fig. 8C). The anti-inflammatory cytokine IL4, CCL1, and IL13 were increased in S196A conditioned media compared to WT BMDMs (Fig. 8C). We also found the mRNA expression of IL1β was lower in S196A versus WT BMDMs by RNA-seq (not shown). Our findings suggest that LXR*α* S196A M1 macrophages suppress lipid accumulation in adipocytes by reducing and/or enhancing the levels of pro-inflammatory and anti-inflammatory cytokines.

## DISCUSSION

We examined the effects of LXR*α* pS196 on atherosclerosis progression and obesity within the same mouse by expressing LXR*α* S196A in all myeloid cells in a bone marrow transplant model in *Ldlr*^-/-^ recipients. We found that expression of LXR*α* S196A increased HDL-C and reduced LDL-C levels. The increase in HDL-C is possibly through recruitment to the liver of LXR*α* S196A expressing resident macrophages (Kupffer cells) in the context of bone marrow transplantation after whole body irradiation to influence liver cells to either increase HDL-C particle secretion or decrease HDL-C uptake (52–54). Consistent with an increase in HDL-C in LXR*α* S196A, we observed a decrease in neutral lipid accumulation in plaque macrophages expressing S196A. We also found that expression of LXR*α* S196A reduced the recruitment of inflammation-prone Ly6C^high^ monocytes, which in turn would reduce these monocytes becoming macrophages, and local macrophage proliferation, thus blunting atherosclerotic plaque formation, although this reduction was opposed by decreased macrophage apoptosis.

We also observed from the CD68+ cells laser captured from the plaque of LXR*α* S196A mice an anti-inflammatory gene expression signature associated with the upregulation of genes involved in oxidative phosphorylation, and mitochondrial activity (Fig. 3B). PGC1*α*, a transcriptional coactivator that coordinates mitochondrial biogenesis (28, 29, 55) and a LXR*α* coactivator (27), was the top putative transcriptional regulator in plaque CD68+ cells expressing LXR*α* S196A. PCG1 is activated by deacetylation via sirutins (30), and one of the top IPA pathways induced by LXR*α* S196A in CD68+ cells is sirutin signaling. This suggests that the non-phosphorylated form of LXR*α* at S196 may help activate PGC1 via deacetylation and preferentially utilize PGC1*α* as a coactivator in this context. In fact, the amount of transcriptional coactivation by PGC1*α* of LXR*α* in cultured cells was greater when cells were treated with both LXR and RXR ligands (27), which we reported reduced LXR*α* S196 phosphorylation and induced LXR*α* non-phosphorylation-dependent target gene expression compared to treatment with either ligand alone (56).

Consistent with changes in genes linked to mitochondrial activity, we demonstrated in S196A BMDMs polarized *in vitro* to the M1 or M2 states, upregulation of basal respiration, ATP production (Fig. 4A), and increased mitochondrial abundance (Fig. 4B-C), and mitochondria morphology reminiscent of M2 BMDMs (Fig 4D). This indicates that both subsets of S196A macrophages utilize fatty acid metabolism to a greater extent than their WT counterparts. Since the rewiring of macrophage metabolism from glycolytic to oxidative phosphorylation is a hallmark of the anti-inflammatory response by M2 macrophages, our findings suggest that S196A expression in CD68+ cells in the plaque promotes the acquisition of the characteristics of anti-inflammatory macrophages (25).

Surprisingly, we also found that expression of LXR*α* S196A in the bone marrow significantly reduced mouse weight, total fat, pWAT, and adipocyte size in pWAT compared to WT (Fig. 5). This phenotype was not reported in macrophage-specific S196A expressing mice in the *Ldlr*^-/-^ background (17), suggesting that other bone marrow-derived cells in combination with macrophages are promoting this phenotype. The decrease in adipocyte size in LXR*α* S196A was associated with reduced numbers of pro-inflammatory ATMs in the adipose tissue (Fig. 6A-E), and a large change in the LXR*α* S196A transcriptomes between ATM subtypes that indicated a less inflammatory phenotype. We did not observe any expression of UCP1 in the pWAT by qPCR or IHC (not shown), suggesting the effect of LXR*α* S196A from bone marrow-derived cells on pWAT was not a result of conversion into beige fat via UCP1 (57). Rather, we suggest the reduction in adipocyte size is a result of a decline in pro-inflammatory ATM infiltration and a decrease in the secretion of inflammatory factors (and/or an increase in the secretion of anti-inflammatory molecules) from LXR*α* S196A compared WT ATMs.

As noted above, our studies indicated that LXR*α* S196A promoted an anti-inflammatory phenotype within the ATMs of adipose tissue. Previous studies demonstrated that reconstitution of wild type mice with LXR*α*/β knock-out bone marrow, which would be predicted to exacerbate obesity due to a reduction in the LXRs anti-inflammatory response, had no effect on diet-induced obesity (58). However, in *ob/ob* mice that were LXR*α*/β deficient, there was enhanced inflammation and increased macrophage content in adipose tissue, although the mice did not become more obese (59). Thus, it appears that depending on the physiological context, the effects of LXRs in general, and LXR*α* pS196 in particular, on inflammation and obesity vary. It will be interesting to see whether expression of LXR*α* S196A in the bone marrow of *ob/ob* mice would ameliorate obesity, or whether the effect on obesity of LXR*α* S196A in other genetic backgrounds would be the same, since *Ldlr*^-/-^ compared to *Apoe^-/-^* mice are more prone to diet-induced obesity (60, 61) through pathways affecting leptin and AMPK (62).

Given that LXR*α* S196A in our bone marrow transplant model is expressed only in myeloid cells and not in the adipocytes, we hypothesized that in the S196A mice, factors secreted by ATMs in WT are reduced in their abundance if pro-inflammatory, or increased in their levels if anti-inflammatory, which could drive the lipid accumulation in adipocytes. In fact, incubation of 3T3-L1 with conditioned media from M1 LXR*α* S196A BMDMs showed reduced neutral lipid staining compared to M1 LXR*α* WT BMDM conditioned media, consistent with LXR*α* S196A BMDMs having reduced levels of pro-inflammatory (and/or increased levels of anti-inflammatory) secreted factors (Fig. 7B). The myeloid specific expression of LXR*α* S196A was not reported to reduce obesity in the *Ldlr^-/-^* background (17), yet conditioned media from LXR*α* S196A M1 BMDM compared to WT suppressed lipid accumulation in adipocytes in culture. This suggests differences in the abundance and/or activity of inflammatory mediators secreted by LXR*α* S196A M1 BMDMs *in vitro* or LXR*α* S196A when expressed in the bone marrow transplant model relative to the myeloid-specific LXR*α* S196A expression *in vivo*.

It has been previously demonstrated that LXR*α* inhibits the transcription of pro-inflammatory cytokines, such as IL1β, IL6, IL18 and TNF*α* (63, 64), although the effect of LXR*α* S196A on transrepression of genes expressing inflammatory factors remains unclear. In principle, LXR*α* S196A could increase transrepression, thus further inhibiting expression of pro-inflammatory targets (65). We demonstrated that multiple cytokines were differentially secreted by the inhibition of LXR*α* pS196, including a decrease in the pro-inflammatory cytokines IL1β, and an increase in the anti-inflammatory factor IL4 and IL13 (Fig. 7C). Given that IL4 and IL13 promote M2 polarization (65), this suggests that LXR*α* S196A cells could more readily assume M2-like characteristics relative to WT cells. Indeed, the metabolic alterations observed in S196A macrophages (e.g., upregulation of oxidative phosphorylation and mitochondrial biogenesis) are consistent with a more pronounced anti-inflammatory M2 phenotype (65). IL1β is also an attractive candidate since it is a pro-inflammatory factor downregulated in LXR*α* S196A compared to WT M1 conditioned media. A clinical trial (CANTOS) has demonstrated a reduction in atherosclerosis using a therapeutic monoclonal antibody targeting IL1β (66). IL1β is also an established factor involved in macrophage-adipocyte crosstalk and can promote the development of obesity (67). Furthermore, IL1β mRNA expression was reduced in LXR*α* S196A versus WT BMDMs by RNA-seq (not shown). Further experiments using neutralizing antibodies or genetic deletions will be required to define the factor(s) released by macrophages responsible for reduced lipid accumulation in adipocytes by macrophages expressing S196A.

Our findings complement a recent elegant study by Gage *et al.* that demonstrated a myeloid-specific knock-in of LXR*α* S196A resulted in greater macrophage proliferation via the LXR*α* S196A dependent upregulation of FOXM1, and increased atherosclerosis (17). The difference in the phenotypes between the myeloid-specific expression of S196A versus our bone marrow transplant model, in which the entire complement of hematopoietic cells express LXR*α* S196A suggest that LXR*α* S196A expression within additional bone marrow-derived cells restrain macrophage proliferation in the plaque to deter atherosclerosis. Recent advances in single cell RNA-seq demonstrated that different leukocyte clusters with distinct phenotype (including T cells, B cells, macrophages, monocytes and NK cells populations) were present in the plaque of *Ldlr^-/-^* mice (68). Elucidating which cell subset influences macrophage proliferation within the plaque remains an open question and one we are actively pursuing.

In summary, we showed that reducing LXR*α* pS196 in bone marrow cells of *Ldlr^-/-^* mice reduced VAT weight and attenuated atherosclerosis. This was accompanied by a shift in macrophages to a less inflammatory state in LXR*α* S196A compared to WT in both tissue microenvironments through reprograming of LXR*α*-dependent transcription. However, the specific genes that change in expression were largely non-overlapping between plaques and VAT macrophages (TableS4). This is similar to recent single-cell RNA sequencing of macrophages from plaques and adipose tissue in western diet fed mice (8, 12).

Reducing LXR*α* pS196 in the bone marrow results in favorable effects on macrophage metabolism that are beneficial in reducing atherosclerosis and obesity. This is reminiscent of the effects of exercise on reducing macrophage inflammation (69, 70). We speculate that exercise could lower LXR*α* phosphorylation within myeloid cells to impart a more anti-inflammatory and metabolically favorable gene expression program by LXR*α*, thus helping to ameliorate atherosclerosis and obesity.

## ACKNOWLEDGEMENTS

We thank Linda Chenane and Delphine Dehée (Garabedian lab) for their help with staining of the plaques. We thank Hannah Weber and Susan Ha for critically reading the manuscript. We also thank Dr. Shruti Rawal (Fisher lab) for the smooth muscle cell actin staining protocol, and Dr. Ravichandran Ramasamy for insight into the mitochondrial and adipocyte phenotypes. This work was support by grants from the NIH (R01HL117226, MJG, EAF), and an American Heart Association postdoctoral fellowship (19POST34410010, MV) and Jan Vilcek/David Goldfarb Fellowship (MV). We thank the Scientific Cores and Shared Resources at NYU Langone Health supported by the Laura and Isaac Perlmuter Cancer Center (NCI P30CA16087), specifically the High Throughput Biology Laboratory, Cytometry and Cell Sorting Laboratory, Experimental Pathology Research Laboratory, Genome Technology Center, Applied Bioinformatics Laboratory and Microscopy Laboratory.

## CONTRIBUTIONS

M.V. designed and conducted experiments and analyzed the data. She collaborated with E.S., and C.R. on the atherosclerosis model, with H.R.C, and I.J.G on flow cytometry of adipose tissue macrophages, with T.J. and T.J.B. on flow cytometry of blood, with R.R., C.A.N., and M.L.G. on the mouse experiments. E.J.B. conducted RNA-seq analysis on adipose tissue macropahges. I.P. provided the mice and critical comments on data. M.V., E.A.F and M.J.G. analyzed the data and wrote the manuscript. E.A.F. and M.J.G. developed the concept, designed experiments and directed the study. All the authors provided critical comments on the manuscript.

## COMPETEING INTERESTS

The authors have no competing interests.

## MATERIALS AND CORRESPODANCE

Materials request and correspondence should be addressed to Michael J. Garabedian, NYU School of Medicine, 450 E. 29th Street, New York, NY 10016 or Edward A. Fisher, NYU School of Medicine, SB 705, 435 E. 30^th^ St., New York, NY 10016

## MATERIALS AND METHODS

### Animals

C57BL/6 LXR*α* S196A mice were generated by Ozgene, and have been described (17, 19). The mice were cared for in accordance with the National Institutes of Health guidelines and the NYU Institutional Animal Care and Use Committee. Bone marrow from male LXR*α* WT or LXR*α* S196A were transplanted into 10 week old male *Ldlr^-/-^* mice (Jackson Laboratory; Stock No: 002207). After 4 weeks after the bone marrow transplant, the mice were placed on a Western diet (42% kcal fat, 0.3% kcal cholesterol, Research Diets) for 16 weeks to allow development of moderately advanced atherosclerotic plaques. To study monocyte recruitment, Edu (1mg/30g mouse weight) was injected intraperitoneally (IP) 3 days before harvest to label Ly6C^high^ monocytes, and circulating Ly6C^low^ monocytes were labeled by injecting mice in the retro-orbital vein with 250 µL of 1µm Fluoresbrite fluorescein isothiocyanate-dyed (YG) plain microspheres (Polysciences Inc.) diluted in PBS (1:4) 1 day before harvesting as described (21). Mice were weighed once a week and on harvest day. Mice were anesthetized with xylazine/ketamine, and blood was collected via cardiac puncture for plasma analyses. Mice were perfused with 10% sucrose/saline. Organs weight was measured. Aortic roots were dissected and embedded in optimal cutting temperature (OCT) compound medium and frozen immediately, and stored at −80°C until further use. pWAT was collected and digested for immune cells analysis, and adipocytes were collected in Trizol (Invitrogen). Liver, pWAT and BAT were fixed in formalin and then embedded in paraffin for further IHC studies. The data presented are from male mice to focus on a single experimental cohort with similar body size and hormonal milieu.

### Mouse genotyping

Genotyping was performed by mouse tail DNA PCR analysis. Genotyping was performed using the following PCR primers: R2: 5′-AAGCATGACCTGCACACAAG-3′; WT: 5′-GGTGTCCCCAAGGGTGTCCT-3′; and SA: 5′-GGTGTCCCCAAGGGTGTCCG-3′. Primers R2 and WT identify the wild-type allele (642-bp) and primers R2 and SA identify mutant allele of LXR*α* S196A global knock-in mice (656-bp).

### Blood adiponectin and plasma lipoprotein measures

Blood samples were centrifuged at 10,000 rpm for 10 minutes and the supernatants collected as plasma. Plasma adiponectin concentration was quantified using the adiponectin mouse ELISA kit (Thermo Fisher Scientific). Total cholesterol, triglyceride, non-esterified fatty acid and HDL-C concentrations in plasma were measured using colorimetric assays (Wako Diagnostics). For plasma lipoprotein analysis: for each group, equal volume of plasma per mice were pooled. Lipoproteins were separated by fast-performance liquid chromatography (Superose 6 10/300 GL column, GE Healthcare, PA, USA) on a Shimadzu HPLC system. Following separation, fractions were collected and cholesterol content was quantified by colorimetric assays.

### Body fat composition

We assessed the total fat mass, lean mass, and bone mineral density (BMD) of mice after 10 weeks of western diet feeding using the LUNAR PIXIMUS bone densitometer (Lunar Corp.). Mice were anesthetized by continuous-inhalation of isoflurane and placed in a prone position on the detector tray to scan the entire mouse body. This non-invasive technique provides quantitative data on the fat tissue content, the lean tissue content, and BMD.

### Histochemical analyses

To measure lesions in the aortic root, the heart and proximal aorta were excised and the apex and lower half of the ventricles were removed. Tissues were sectioned at 6µm intervals using a cryostat (for frozen blocks) or a microtome (for paraffin blocks). For frozen tissue sections, slides were fixed in acetone for 10 minutes at −20 °C. For paraffin sections, slides were dewaxed in xylene, rehydrated and boiled for 20 minutes in citrate buffer (10mM, pH 6.0) for antigen retrieval. After washing three times with phosphate-buffered saline, tissue sections were incubated with primary antibodies diluted in blocking solution (10% BSA and horse serum in PBS) overnight at 4°C in a humidified chamber. Aortic roots frozen in OCT were stained for CD68 (1:250, MCA1957, Biorad), BAT was stained for UCP1 (1:300, ab23841, abcam), and pWAT was stained for F4/80 (1:200, MCA497RT, Biorad). Sections were washed and incubated with the appropriate biotinylated secondary antibodies (1:1,000, Vector laboratories), followed by washing and visualization using Vectastain ABC kit (Vector Laboratories) for CD68 or followed by horseradish peroxidase streptavidin (Vector Laboratories) and visualized using the DAB (3,3′-diaminobenzidine) peroxidase substrate kit (Vector Laboratories). The reaction mixture was counterstained with hematoxylin. BAT, liver, and pWAT were also stained for hematoxylin and eosin (H&E). Aortic root sections were stained for collagen content using the Pico Sirus Red Stain for 90 minutes following the manufacturer instructions (Poly Sciences), and imaged under linear polarized light with a Zeiss AxioPlan Upright. Oil Red O staining was used to detect lipid accumulation in aortic root plaques. Formaldehyde-fixed frozen sections of the aortic arch were stained for 15 minutes in 0.3% Oil Red O (Sigma-Aldrich) dissolved in 60% isopropanol and then counterstained with hematoxylin (Vector Laboratories). Microscopic images were taken at 10X, or 4X for aortic root sections using the EVOS FL COLOR system (Thermo Fisher Scientific). Plaque and CD68 staining morphometric measurements were performed using Image Pro Plus software (Micro Optical Solutions). Measurement of adipocyte size in pWAT and lipid droplets in the liver was performed by automatic counting and area measurement with Image J software using the MRI adipocyte tool.

### Immunofluorescence on tissues

Recruited Ly6C^high^ EdU positive cells in the aortic root plaques were detected using Click-iT EDU Imaging Kit (MP 10338). Recruitment of Ly6C^low^ beads were counted in the plaques using a fluorescent microscope, EVOS FL COLOR (Thermo Fisher Scientific). Apoptosis in the plaque was analyzed by staining the plaques with anti-cleaved caspase 3 (1:100, 9664, Cell signaling). Cell proliferation was analyzed by staining the plaques for Ki67 (1:100, ab1667, abcam). M1, M2 macrophages were measured with iNOS-AF594 (1:100, ab209027, abcam), Arginase 1-AF488 (1:100, 53-3697-82, Invitrogen) and alpha smooth muscle actin with SMaA-AF488 (1:100, 53-9760-82, Invitrogen). Aortic root plaques were also stained for CD68 (1:250, MCA1957, Biorad) and DAPI (ProLong™ Gold Antifade Mountant with DAPI, P36931, Thermo Fisher Scientific). Number of double positive cells was measured for each mouse.

### Hematology parameters by flow cytometry

Blood was collected before sacrificing the mice by tail bleeding. Total white blood cells (WBCs) were determined using the Genesis Hematology System (Oxford Science, Oxford, CT). Red blood cells were lysed with red blood cells lysis buffer (Sigma-Aldrich) and cells were stained with Percp/Cy5.5 anti-mouse CD45, PE anti-mouse CD115, and APC anti-mouse Ly6C/6G (Biolegend). Monocytes and neutrophils were identified by flow cytometry using a LSRII analyzer and analyzed using FlowJo v10.

### Laser capture microdissection and RNA-seq analysis

6 µm sections of aortic roots were collected on Pen membrane Frame Slides (Arcturus). CD68+ cells were isolated from atherosclerotic plaques by laser capture microdissection performed under RNase-free conditions (71, 72). Aortic root sections were stained with hematoxylin-eosin and cells were captured from approximately 36 frozen sections. After laser capture microdissection, RNA was isolated using the PicoPure RNA isolation kit (Thermo Fisher scientific), and quality and quantity were determined using an Agilent 2100 Bioanalyzer (Agilent Technologies). RNA-seq libraries were prepared using the Clontech SMARTer Stranded Total RNA-Seq Kit - Pico Input Mammalian following the manufacturer’s protocol. Libraries were purified using AMPure beads, pooled equimolarly, and run on a HiSeq 2500, paired end reads. FASTQ files were obtained and STAR 2.5 was used for read mapping against the mm10 reference genome. BAM alignment files were processed using Featurecounts 1.0.5. DESeq2 (3.4) was used to identify differentially expressed genes (DEG). Genes with a p-value < 0.05 were determined to be differentially expressed (Fold change >1.5) and the Ingenuity Pathway Analysis (Qiagen) was used for pathway and transcription factor analysis.

### Bone marrow derived macrophages cell culture

Bone marrow cells were isolated by flushing cells from the femurs and tibiae of mice. Cells were differentiated into bone marrow derived macrophages (BMDMs) in 4.5 g/L glucose DMEM (Lonza) with 20% fetal bovine serum (FBS), 1% penicillin/streptomycin, and murine M-CSF (10 ng/mL; PeproTech) at 37 °C and 5% CO_2_ for 7 days. The BMDMs were treated with LPS (10ng /mL, Sigma-Aldrich) and murine IFNγ (20 ng/mL, BD Biosciences) for M1 polarization, with murine IL4 (15 ng/mL; BD Biosciences) for M2 polarization or without treatment for control M0, for 24 hours.

### Mitochondrial activity: Seahorse assay

Oxygen consumption was measured in a Seahorse XF24 Analyzer (Agilent). Differentiated BMDMs were seeded at 0.25×10^6^ cells/well in XF24-well plates (Agilent) and polarized for M0, M1, and M2 for 24 hr, as described above. Wells without cells were included as a background control. Sensor plates were calibrated overnight in a CO_2_-free incubator at 37°C using 1 mL/well of XF calibrant solution (Agilent). After polarization, culture medium was replaced with glucose-free DMEM (Agilent) medium supplemented with 2 µM L-glutamine, 1 µM pyruvate and 10 mM glucose (Agilent) and cells were incubated 1 hr in a CO_2_-free incubator at 37°C. Injection ports were loaded with 10X injection mixes to obtain a final concentration in each well of 20 mM glucose after the first injection, 1.5 µM oligomycin after the second injection, 1.5 µM FCCP after the third injection, and 0.5 µM rotenone plus 0.5 µM antimycin A after the fourth injection. After the run, supernatants were carefully aspirated and cells were fixed with methanol and stained for DAPI (Invitrogen). Cells were imaged and counted using the CellInsight CX7 LZR High Content Analysis Platform (Thermo Fisher Scientific). OCR values were normalized to cell number for each well.

### Determination of mitochondrial abundance: MitoTracker

BMDMs were seeded at 1×10^5^ cells/well in 8 wells Nunc Lab-Tek II Chamber Slide (Thermo Scientific). The next day, cells were polarized for M0, M1, and M2 as described above. After 24 hours, cells were treated with 100 nM MitoTracker Red CMXRos for 30 minutes at 37°C, fixed in methanol and stained for DAPI (Invitrogen). Images were taken using an EVOS m7000 Imaging System.

### Determination of mitochondrial abundance: mitochondrial DNA measure

BMDMs were seeded at 0.6×10^6^ cells/mL in 12-well plates and differentiated as described above. The next day, total DNA was extracted from BMDMs using a DNeasy Blood & Tissue Kit (Qiagen) and 25 ng of DNA was used for qPCR using Fast SybrGreen Master Mix (Applied Biosystems) on a QuantStudio 6 Flex (Applied Biosystems). Mitochondrial DNA (mtDNA) expression was normalized by nuclear DNA (nDNA) expression. Sequences of primers used are mtDNA F: 5’-GTACCGCAAGGGAAAGATGA-3’; mtDNA R: 5’ACCAAGCTCGTTAGGCTTTTC-3’; nDNA F: 5’-GCCAGCCTCTCCTGATTTTAGTGT-3’; nDNA R: 5’-GGGAACACAAAAGACCTCTTCTGG-3’.

### Electron Microscopy

M0, M1 and M2 LXR*α* WT and S196A BMDM were fixed with 2.5% glutaraldehyde and 2% paraformaldehyde in 0.1 M cacodylate buffer, and postfixed in 1% OsO_4_ in 0.1 M cacodylate buffer with 1% potassium ferrocyanide for 1 h on ice. The cells were stained *en bloc* with 0.25% uranyl acetate then dehydrated in graded series of ethanol on ice followed by one wash with 100% ethanol and two washes with propylene oxide (5 min each) and embedded with EMbed 812 (Electron Microscopy Sciences, Hatfield, PA). Sections were cut at 60 nm on a Leica EM UC6 ultramicrotome, and picked up on copper grids, stained with 3% uranyl acetate for 15 min and Reynolds lead citrate stain for 5 min. Grids were viewed using a Philips CM12 TEM (Philips) transmission electron microscope and photographed using a Gatan 4k x 2.7k digital camera (Gatan Inc.).

### 3T3-L1 cell culture and differentiation

3T3-L1 cells were cultured in 4.5 g/L glucose DMEM (Lonza) with 10% FBS, 1% penicillin/streptomycin (complete DMEM) at 37 °C and 5% CO_2_. 3T3-L1 preadipocytes were induced to differentiate by incubation with differentiation medium containing 0.5 mM IBMX (Sigma-Aldrich), 1 µg/mL insulin (Gibco), 0.25 µM dexamethasone (Sigma-Aldrich) and 2 µM rosiglitazone (Cayman Chemical Company) in complete DMEM for 2 days. The cells were then incubated with complete media containing 10 µg/mL insulin for 3 days. Then conditioned media from M0, M1, or M2 BMDMs was filtered and added to the differentiated 3T3-L1 for 24 hours. Lipid accumulation was observed by staining with Oil Red O (ORO) staining: cells were fixed with 4% formaldehyde for 15 minutes, incubated with 60% isopropanol (Fisher Chemical) for 5 minutes, stained with 0.3% Oil Red O (Sigma-Aldrich) in 60% isopropanol for 30 minutes at room temperature and counterstained with hematoxylin (Vector Laboratories). Images were taken using the EVOS FL COLOR imaging system.

### Determination of secreted cytokines

Cytokine levels in cell culture supernatants samples were determined using the Proteome Profiler Mouse Cytokine Array Panel A (R&D Systems) following the manufacturer recommendations.

### Separation of Stromal Vascular Cells (SVC)

Isolation was performed as described by Weisberg *et al* (73). Perigonadal fat pads were minced, placed in DMEM supplemented with 10 mg/ml fatty acid-poor BSA, and centrifuged at 1000 × *g* for 10 minutes. A LPS-depleted collagenase mixture (Liberase^TM^, Roche Applied Science) at a concentration of 0.03 mg/ml and 50 units/ml DNase I (Sigma) was added to the tissue, and the samples were incubated at 37 °C in a rotating shaker for 45 minutes. Then, the samples were passed through a 70-µm nylon cell strainer (Corning). The suspension was centrifuged at 1000 × *g* for 10 minutes, and the pelleted cells were collected as the SVC. The floating cells were collected as the adipocytes. The SVC were resuspended in red blood cells lysis buffer (Sigma-Aldrich) and incubated at room temperature for 5 minutes and washed in PBS.

### RT-qPCR of adipocytes

Total RNA was extracted from adipocytes with TRIzol (Invitrogen) and was reverse transcribed into cDNA using the Verso cDNA Synthesis Kit (Thermo Fisher scientific) according to the manufacturer’s instructions. Quantitative real-time PCR was performed on the QuantStudio 6 Flex (Applied Biosystems) using SYBR Green Fast Master Mix (Applied Biosystems). Gene-expression levels were calculated using the ΔΔCt method after their normalization to the expression levels of Cyclophilin A1 (F: 5’-GGCCGATGACGAGCCC-3’, R: 5’-TGTCTTTGGAACTTTGTCTGCAA-3’). The sequences of the mouse primers used for qPCR are: Adipoq F: 5’-TGTTCCTCTTAATCCTGCCCA-3’, 5’-CCAACCTGCACAAGTTCCCTT-3’; Fabp4 F: 5’-AAGGTGAAGAGCATCATAACCCT-3’, R: 5’-TCACGCCTTTCATAACACATTCC-3’; Plin2 F: 5’-TCTGCGGCCATGACAAGTG-3’, R: 5’-GCAGGCATAGGTATTGGCAAC-3’; Atgl F: 5’-ATGTTCCCGAGGGAGACCAA-3’, R: 5’-GAGGCTCCGTAGATGTGAGTG-3’; Hsl F: 5’-GATTTACGCACGATGACACAGT-3’, R: 5’-ACCTGCAAAGACATTAGACAGC-3’.

### Immune cell profiling

SVC were resuspended in 2% Fc Block (553142, BD Pharmingen) and blocked for 30 minutes. Then fluorophore-conjugated primary antibodies were incubated for 30 minutes: PE/Cy7 anti-mouse CD45 (102114, Biolegend), APC anti-mouse F4/80 (MCA497APC, Biorad), PE anti-mouse CD11b (RM2804, Invitrogen), PE-Texas Red anti-mouse CD11c (MCD11C17, Invitrogen), BV786 anti-mouse CD3 (564379, BD Biosciences), PerCP/Cy5.5 anti-mouse CD4 (100540, Biolegend), APC-Cy7 anti-mouse CD8 (557654, BD Biosciences), and BV510 anti-mouse CD25 (740106, BD Biosciences) in FACS buffer (1X HBSS (Thermo Fisher Scientific), 2% BSA and 0.5 mM EDTA). After one wash in FACS buffer, cells were resuspended in FACS buffer containing 1 µg/mL DAPI (Invitrogen). Immune cells profiling was performed using a FACSAria II cell sorter (BD Biosciences) and data were analyzed with FlowJo v10. Adipose tissue macrophages (FB and FBC) were sorted and collected.

### Adipose tissue macrophages RNA-seq

ATM RNA was extracted with RNeasy micro kit (Qiagen), and quality and quantity were determined using an Agilent 2100 Bioanalyzer (Agilent Technologies). RNA-seq libraries were prepared using the Trio Low Input RNA-seq library preparation kit (NuGEN) following the manufacturer’s protocol. Libraries were purified using AMPure beads, pooled equimolarly, and run on a HiSeq 2500, paired end reads. Reads were mapped to the mm10 genome using STAR 2.5. Differential expression was assessed using Cufflinks v 2.2.1, using the cuffdiff module and the parameter of dispersion-method poisson to estimate p-values, using the Gencode vM18 gene annotation. Genes with a p-value < 0.05 and a Fold change > 1.5 were considered likely to be significantly differentially expressed and run in Ingenuity Pathway Analysis (Qiagen) for pathway analysis; genes with an FPKM of 0 were excluded from further analysis.

### Statistical analysis

Statistical analyses were performed using GraphPad Prism software. Data are reported as mean ± SEM. Number of replicates (n) and statistical tests used are described in the figure legends, with a p value < 0.05 being considered significant and levels of significance denoted as *p < 0.05; **p < 0.01; and ***p < 0.001. In general, a two-tailed unpaired Student’s t test was used when comparing two groups, and a two-way ANOVA was used when comparing interactions between genotype (LXR*α* WT or LXR*α* S196A) and stimulus (M0, M1, M2). For *in vivo* experiments, n = number of animals.

---

**Supplemental Figure 1:**
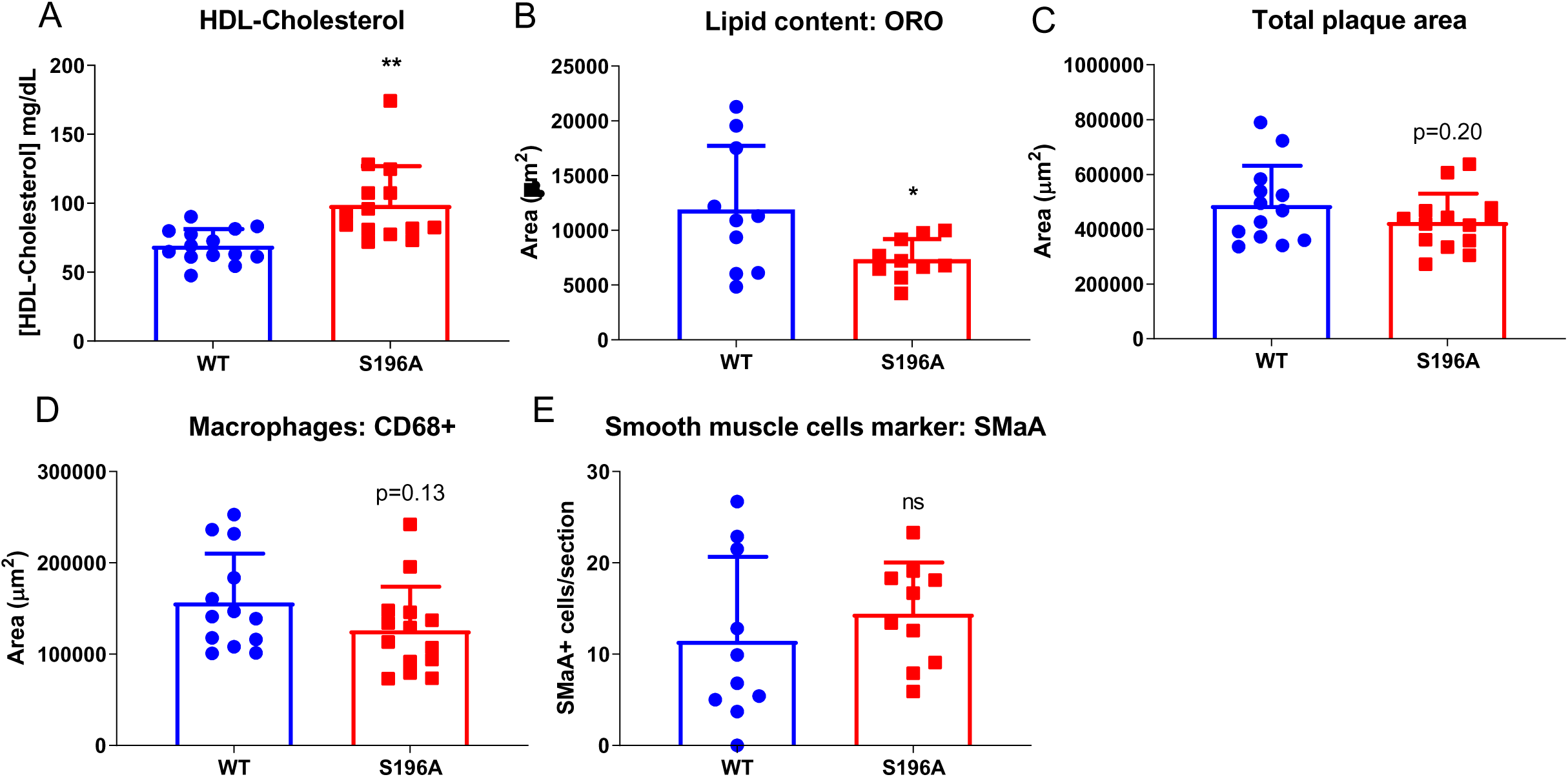
Effect of LXRα S196A on HDL-Cholesterol and aortic plaque characteristics. (A) HDL-Cholesterol measured by ELISA. (B) Lipid content (Oil Red O), (C) total plaque area quantification, (D) area of macrophages (CD68+) were measured after staining of aortic sections. Number of cells per section expressing (E) iNOS (M1 marker), (F) Arginase 1 (M2 marker), and (G) SMaA (smooth muscle cell actin marker) were determined after staining of aortic sections. Data are expressed as mean ± SD (n=14 per group, except for ORO, iNOS, Arginase1 and SMaA n=10). T test; *P<0.05 and **P<0.01.

**Supplemental Figure 2:**
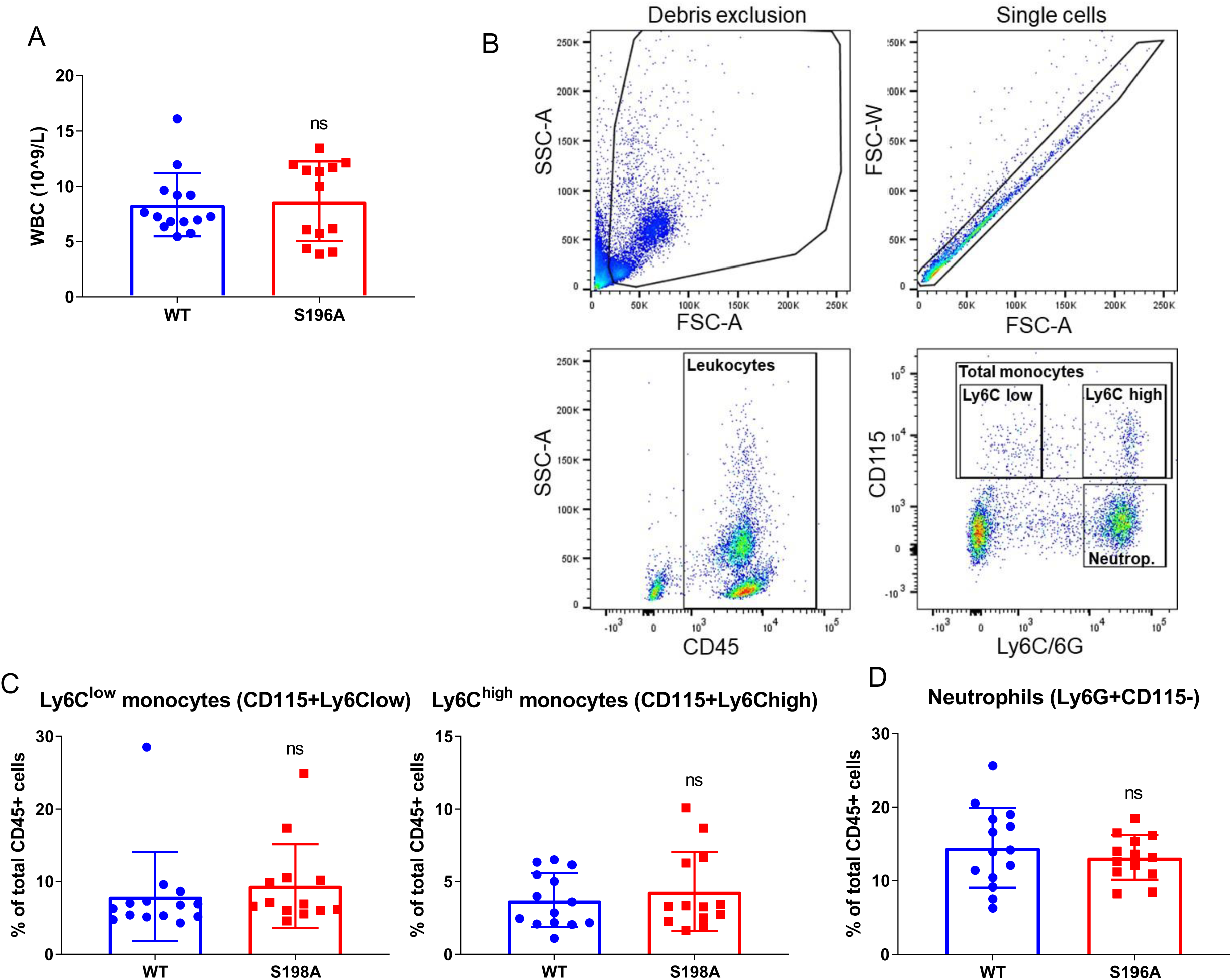
Effect of LXRα S196A on monocytes and neutrophils in the blood. (A) Total number of white blood cells (WBC). (B) Gating strategy for monocytes and neutrophils quantification by flow cytometry. (C) Ly6C-high and Ly6C-low monocytes, and (D) neutrophils were measured by flow cytometry. Data are expressed as mean ± SD (n=14 per group). T test; **P<0.01.

**Supplemental Figure 3:**
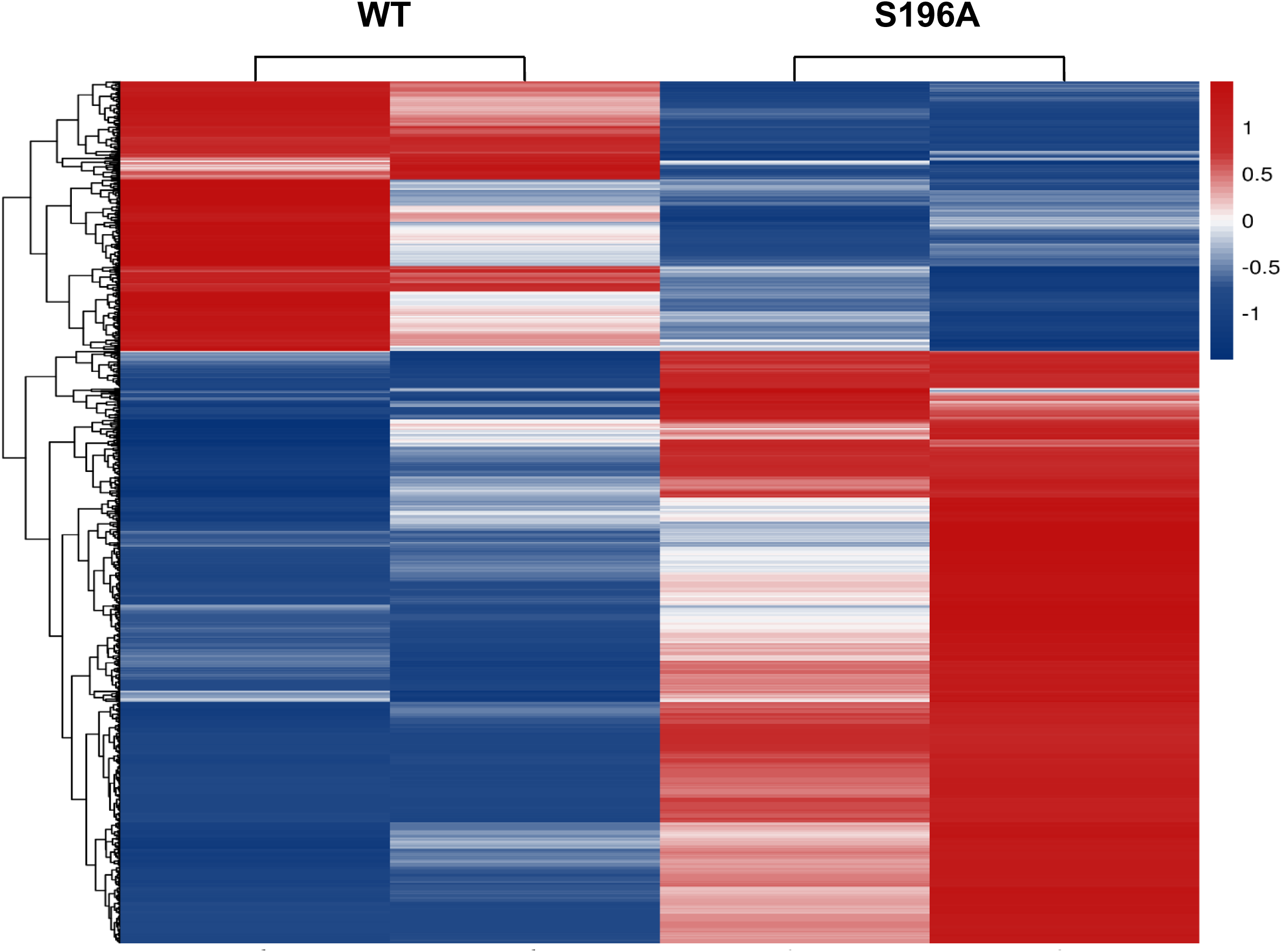
Heatmap of plaque macrophages captured by LCM from WT and LXRα S196A (RNAseq): 1133 significant genes; Fold change > 1.5; p<0.05.

**Supplemental Figure 4:**
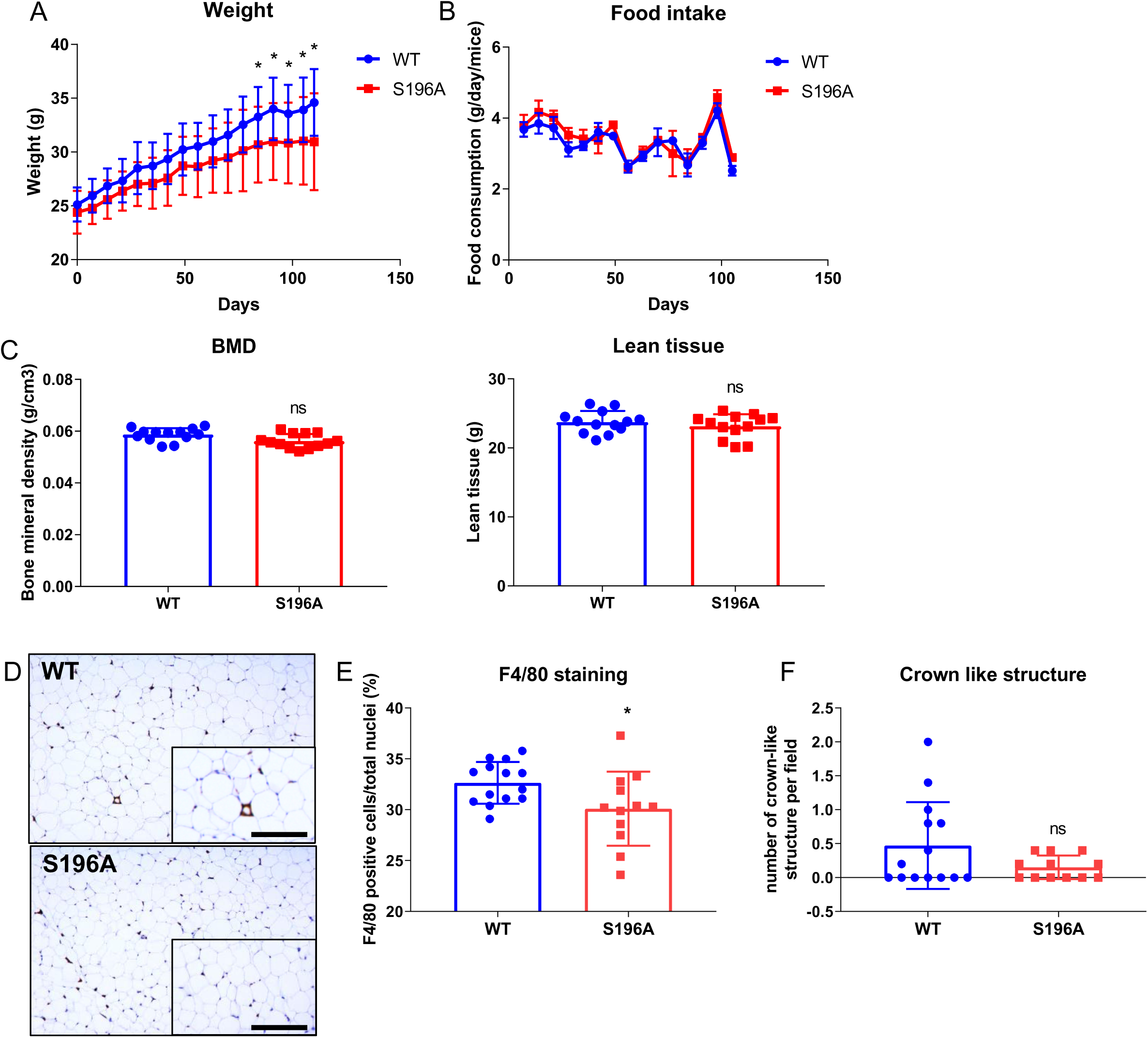
Effects of LXRα S196A bone marrow transplantation on diet induced obesity. (A) Mouse body weight, (B) food intake, (C) bone mineral density and lean tissue quantification by DEXA scan were determined from mice reconstituted with bone marrow from WT and S196A mice. (D) Representative F4/80 staining images (scale bar = 400 µm), (E) quantification of F4/80 staining and (F) of crown-like structures of pWAT sections. Data are expressed as mean ± SD (n=14 per group). T test; *P<0.05.

**Supplemental Figure 5:**
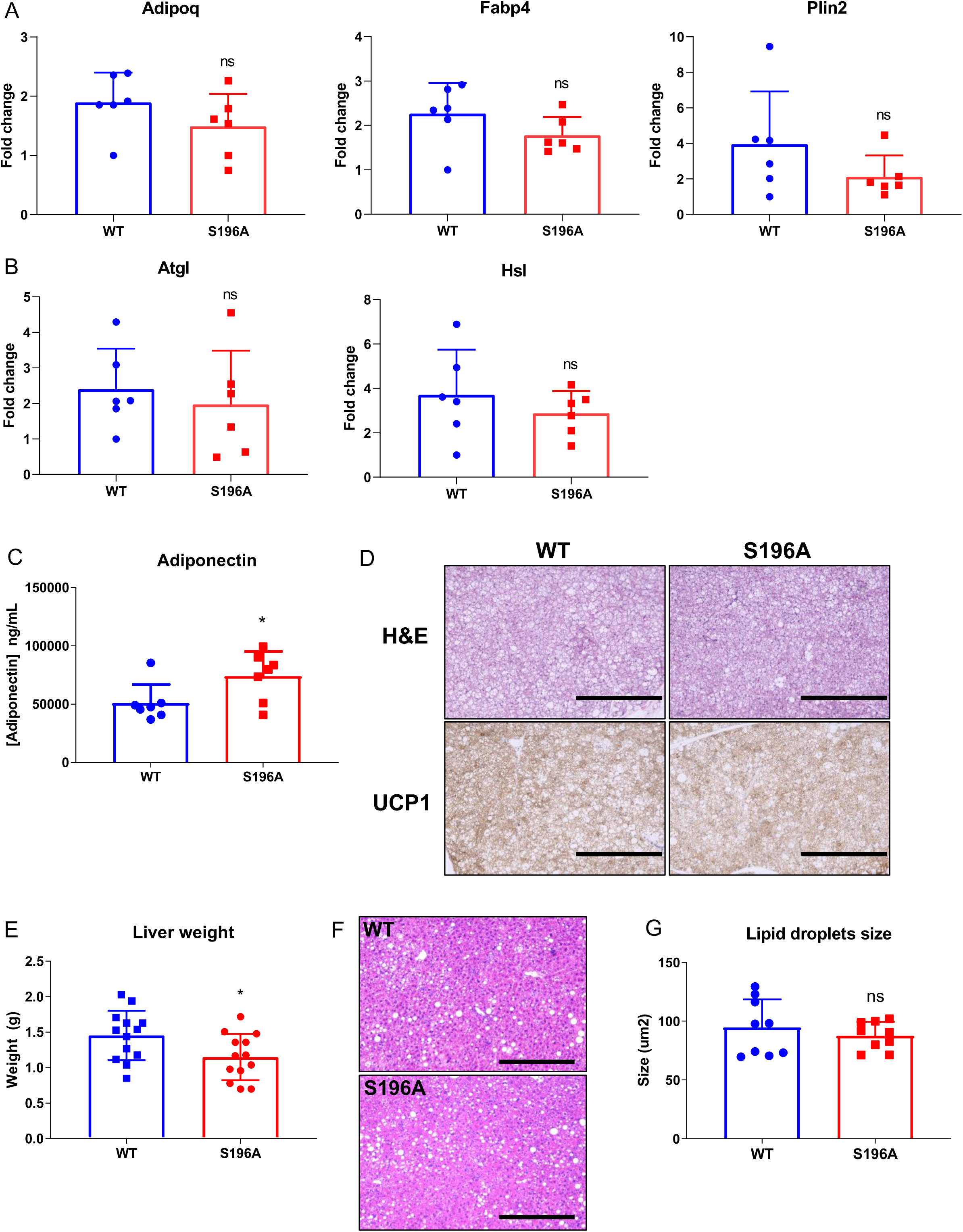
LXRα S196A increase adiponectin level but doesn’t affect BAT or liver phenotype. (A) Gene expression of adipocyte differentiation marker: *Adipoq* (Adiponectin), *Fabp4* (Fatty acid binding protein 4) and P*lin2* (Perilipin2), and (B) lipolysis activators: *Atgl* (Adipose triglyceride lipase) and *Hsl* (Hormone-sensitive lipase) (C) Adiponectin levels and (D) representative image of hematoxylin and eosin and UCP1 staining of brown adipose tissue (BAT) (scale bars: 400µm). (E) Liver weight, (F) images of representative hematoxylin and eosin stained sections (scale bar = 400 µm), and (G) quantification of lipid droplets from hematoxylin and eosin stained liver sections. Data are expressed as mean ± SD (n=6 for gene expression, n=7 for adiponectin level, n=14 for liver weight and n=8 for lipid droplet quantification in liver). T test; *P<0.05 and **P<0.01.

**Supplemental Figure 6:**
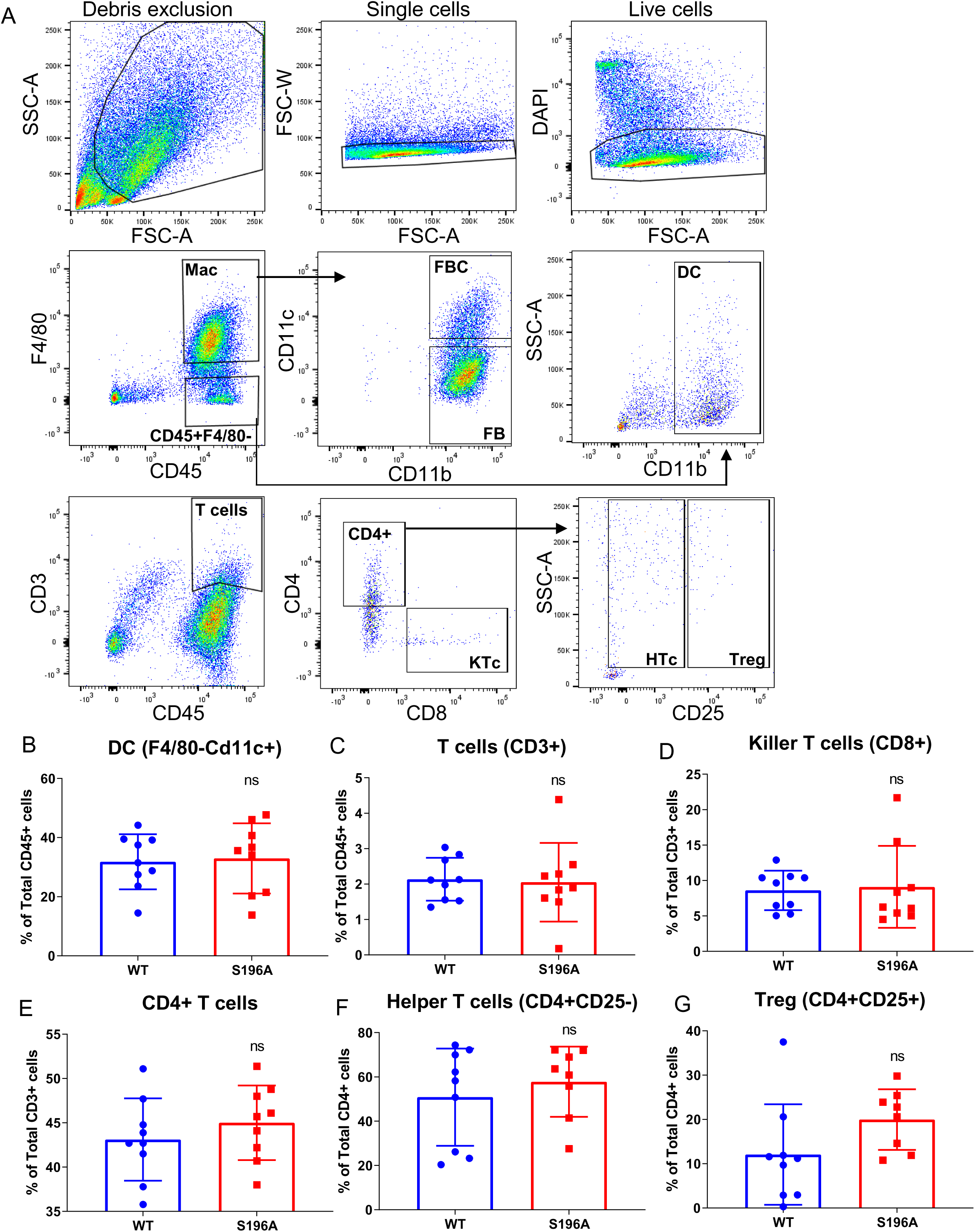
Effects of LXRα S196A bone marrow transplantation on immune cells in adipose tissue. (A) Gating strategy for immune cells population. (B-G) Quantification of immune cells by flow cytometry in perigonadal adipose tissue from WT and S196A mice. (B) Dendritic cells (DC), (C) total T cells (CD3^+^), (D) Killer T cells (CD8^+^), (E) CD4^+^ T cells, (F) Helper T cells (CD4^+^CD25^-^) and (G) Treg (CD4^+^C25^+^) populations. Data are expressed as mean ± SD (n=9 per group).

Supplementary Table 1: Genes differentially expressed in LXRα S196A versus WT from plaque CD68+ cells

Supplementary Table 2: Genes within IPA pathways in LXRα S196A versus WT from plaque CD68+ cells

Supplementary Table 3: Genes differentially expressed in ATMs from LXR WT and S196A

Supplementary Table 4: Genes within IPA pathways unique to LXRαWT and S196A in ATMs

Supplementary Table 5: Genes common between plaque CD68+ cells and ATMs

