## Supplementary Tables 1-5 for "Inhibiting LXR*α* Phosphorylation in Hematopoietic Cells Reduces Inflammation and Attenuates Atherosclerosis and Obesity"

TABLE 1
Genes differentially expressed in LXRα S196A versus WT from plaque CD68+ cells

| S196A vs WT | | |
|---|---|---|
| **gene** | **pvalue** | **FC** |
| Atp2a2 | 0 | 4.408 |
| Myom1 | 0 | 4.627 |
| Dsp | 0 | 6.063 |
| Ldb3 | 0 | 6.498 |
| Podn | 0 | 6.635 |
| Rbm20 | 0 | 7.210 |
| Akap6 | 0 | 7.568 |
| Angpt1 | 0 | 7.890 |
| Eef1a2 | 0 | 8.000 |
| Tcap | 0 | 8.168 |
| Fsd2 | 0 | 8.282 |
| Fhod3 | 0 | 8.398 |
| Stard10 | 0 | 8.754 |
| Popdc2 | 0 | 8.877 |
| Ppargc1a | 0 | 9.063 |
| Scn5a | 0 | 9.190 |
| Ckm | 0 | 9.254 |
| Nebl | 0 | 10.411 |
| Alpk2 | 0 | 10.411 |
| Adprhl1 | 0 | 10.483 |
| Fndc5 | 0 | 10.483 |
| Pygm | 0 | 10.703 |
| Ttn | 0 | 11.236 |
| Art3 | 0 | 13.454 |
| Sbk3 | 0 | 14.320 |
| Corin | 0 | 18.252 |
| Csrp3 | 0 | 20.535 |
| Nppb | 0 | 21.112 |
| Txlnb | 0 | 25.281 |
| Srl | 0 | 25.634 |
| Cmya5 | 0 | 26.723 |
| Lmod2 | 0 | 27.858 |
| Casq2 | 0 | 29.446 |
| Ckmt2 | 0 | 30.484 |
| Mybpc3 | 0 | 35.753 |
| Mylk3 | 0 | 36.002 |
| Pln | 0 | 39.671 |
| Hrc | 0 | 39.947 |
| Kcnj3 | 0 | 40.504 |
| Obscn | 0 | 47.835 |
| Actn2 | 0 | 54.192 |
| Ryr2 | 0 | 54.948 |
| Myom2 | 0 | 60.548 |

| | | |
|---|---|---|
| Tnnt2 | 0 | 73.009 |
| Mb | 0 | 74.028 |
| Myl4 | 0 | 91.139 |
| Myh6 | 0 | 604.668 |
| Pde4dip | 1E-15 | 2.848 |
| Nexn | 1E-15 | 4.500 |
| Myo18b | 1E-15 | 5.502 |
| Mylk4 | 1E-15 | 7.464 |
| Lmo7 | 2E-15 | 4.659 |
| Frmd5 | 1.6E-14 | 6.589 |
| Chrm2 | 1.7E-14 | 5.938 |
| Atp1b1 | 2.8E-14 | 5.736 |
| Rbm24 | 2.9E-14 | 7.210 |
| Trim55 | 3.8E-14 | 7.160 |
| Tmem182 | 5.3E-14 | 7.111 |
| Cox8b | 7.3E-14 | 6.964 |
| Fgf12 | 8.4E-14 | 6.964 |
| Cap2 | 9.5E-14 | 6.774 |
| Cryab | 1.03E-13 | 4.790 |
| Dkk3 | 1.16E-13 | 3.074 |
| Klhl31 | 1.68E-13 | 6.821 |
| Alas2 | 2.51E-13 | 5.696 |
| Tbx5 | 5.68E-13 | 6.543 |
| Mypn | 1.14E-12 | 6.409 |
| Mlip | 1.35E-12 | 6.277 |
| Tmod1 | 1.94E-12 | 4.823 |
| Cd5l | 4.33E-12 | 0.409 |
| Ppp1r3c | 4.39E-12 | 5.979 |
| Ldhb | 6.73E-12 | 5.063 |
| Smyd1 | 7.96E-12 | 5.897 |
| 1110002E2 | 1.21E-11 | 5.856 |
| Perm1 | 1.32E-11 | 5.816 |
| Nrap | 1.75E-11 | 5.134 |
| Hfe2 | 3.25E-11 | 5.657 |
| Abcc8 | 5.48E-11 | 5.540 |
| Alpk3 | 6.35E-11 | 5.426 |
| Clip4 | 7.79E-11 | 4.857 |
| Fbln1 | 8.72E-11 | 3.784 |
| Usp13 | 9.98E-11 | 4.691 |
| Eno3 | 1.25E-10 | 4.757 |
| Pam | 1.25E-10 | 2.713 |
| Ank1 | 1.57E-10 | 4.627 |
| Vldlr | 2.09E-10 | 4.084 |
| Kcnk3 | 2.16E-10 | 5.242 |
| Tmem163 | 2.23E-10 | 5.242 |

| | | |
|---|---|---|
| Sparcl1 | 3.69E-10 | 4.691 |
| Nudt4 | 6.87E-10 | 2.868 |
| Pfkm | 7.54E-10 | 4.028 |
| Art1 | 1E-09 | 4.925 |
| Cacna1h | 1.01E-09 | 4.891 |
| Lrrc2 | 1.02E-09 | 4.790 |
| Atp1a2 | 1.5E-09 | 3.945 |
| Xirp1 | 2.12E-09 | 4.377 |
| Sptb | 2.29E-09 | 3.918 |
| Ndrg2 | 2.34E-09 | 4.141 |
| Prob1 | 2.63E-09 | 4.724 |
| Erbb4 | 3.02E-09 | 4.595 |
| Trim54 | 3.15E-09 | 4.691 |
| Ctnna3 | 5.02E-09 | 4.438 |
| Ppp2r3a | 5.21E-09 | 3.053 |
| Myh14 | 6.16E-09 | 4.563 |
| Trim72 | 6.9E-09 | 4.532 |
| Tmem38a | 8.39E-09 | 4.347 |
| Sh3bgr | 9.02E-09 | 4.438 |
| Dsc2 | 1.15E-08 | 4.377 |
| Scd4 | 1.25E-08 | 4.408 |
| Fbxo40 | 1.27E-08 | 4.199 |
| Adcy1 | 1.61E-08 | 4.347 |
| Xirp2 | 2.13E-08 | 4.287 |
| Chchd10 | 2.62E-08 | 4.170 |
| Kcna2 | 2.69E-08 | 0.275 |
| Ccl6 | 3.03E-08 | 0.435 |
| Unc45b | 3.16E-08 | 4.028 |
| AC157017.[illegible] | 4.9E-08 | 4.141 |
| Gsn | 7.69E-08 | 1.945 |
| Mhrt | 8.06E-08 | 4.028 |
| Mfn2 | 1E-07 | 2.297 |
| Filip1 | 1.07E-07 | 3.531 |
| Igf2 | 1.39E-07 | 3.655 |
| Scn7a | 1.57E-07 | 3.918 |
| Trim63 | 1.67E-07 | 3.891 |
| Tecrl | 1.7E-07 | 3.864 |
| Aco2 | 1.71E-07 | 2.346 |
| Ccdc141 | 1.73E-07 | 2.657 |
| Parm1 | 2.27E-07 | 2.908 |
| Coro6 | 2.35E-07 | 3.630 |
| Ank3 | 2.47E-07 | 3.758 |
| Mmp15 | 2.63E-07 | 3.811 |
| Ptgfr | 2.92E-07 | 3.784 |
| Rp1 | 3.57E-07 | 3.732 |

| | | |
|---|---|---|
| Kcng2 | 4.09E-07 | 3.732 |
| Prkaa2 | 6.06E-07 | 3.531 |
| Fhl2 | 6.27E-07 | 3.531 |
| Ank2 | 6.6E-07 | 2.412 |
| Trdn | 6.62E-07 | 3.555 |
| Comp | 7.37E-07 | 2.250 |
| Lrrc10 | 8.55E-07 | 3.555 |
| Igfbp5 | 9.53E-07 | 2.173 |
| Coq8a | 1.63E-06 | 3.387 |
| Tnni3 | 1.68E-06 | 3.387 |
| Acacb | 1.74E-06 | 3.249 |
| Dmpk | 1.82E-06 | 2.908 |
| Klhl41 | 1.88E-06 | 3.411 |
| Ehbp1 | 2.13E-06 | 2.445 |
| Mybphl | 2.37E-06 | 3.317 |
| March1 | 2.51E-06 | 0.470 |
| Wnk2 | 3.01E-06 | 3.294 |
| Abca8a | 3.18E-06 | 3.387 |
| Gpc1 | 3.45E-06 | 2.462 |
| Hspb6 | 3.79E-06 | 3.204 |
| Clec1b | 4.13E-06 | 0.361 |
| Fgf1 | 4.16E-06 | 3.053 |
| Ctsc | 4.24E-06 | 0.525 |
| mt-Tl1 | 4.33E-06 | 3.317 |
| Tnni3k | 4.94E-06 | 3.204 |
| Sgcg | 5.29E-06 | 3.294 |
| Doc2g | 5.93E-06 | 3.249 |
| Epha4 | 6.23E-06 | 2.888 |
| Neb | 6.37E-06 | 3.204 |
| Ces1d | 6.61E-06 | 3.160 |
| Cacna1c | 6.63E-06 | 2.567 |
| Lmntd1 | 6.91E-06 | 3.138 |
| Fabp3 | 6.97E-06 | 3.117 |
| Ppp1r14c | 7.56E-06 | 3.160 |
| Slc2a4 | 7.81E-06 | 3.160 |
| Rhobtb1 | 8.01E-06 | 2.828 |
| Mapk10 | 8.03E-06 | 3.117 |
| Hba-a2 | 8.2E-06 | 3.160 |
| Fitm2 | 8.64E-06 | 2.828 |
| Gm13340 | 8.74E-06 | 2.928 |
| Ak1 | 9.13E-06 | 2.445 |
| Gm15543 | 9.37E-06 | 3.074 |
| Ndufa5 | 9.89E-06 | 2.603 |
| Lgr6 | 1.02E-05 | 3.053 |
| Abi3bp | 1.03E-05 | 2.204 |

| | | |
|---|---|---|
| Slc47a1 | 1.22E-05 | 3.053 |
| Dmd | 1.25E-05 | 2.071 |
| Vegfa | 1.32E-05 | 2.378 |
| mt-Co3 | 1.33E-05 | 2.789 |
| Idh3a | 1.37E-05 | 2.144 |
| Gm28437 | 1.4E-05 | 2.949 |
| Chpt1 | 1.67E-05 | 2.329 |
| mt-Nd3 | 1.69E-05 | 3.074 |
| Krt222 | 1.71E-05 | 2.969 |
| Atp5b | 1.83E-05 | 1.945 |
| Lrrc39 | 1.84E-05 | 3.031 |
| Kctd12 | 1.92E-05 | 0.470 |
| Pdha1 | 1.93E-05 | 2.144 |
| Dnaja4 | 2.08E-05 | 2.732 |
| Gja1 | 2.11E-05 | 1.905 |
| mt-Nd6 | 2.56E-05 | 2.099 |
| Clec12a | 2.79E-05 | 0.483 |
| Tnnc1 | 2.85E-05 | 2.848 |
| St18 | 2.9E-05 | 0.406 |
| Idh2 | 3E-05 | 2.282 |
| Npr3 | 3.24E-05 | 2.549 |
| Ndufb9 | 3.62E-05 | 2.282 |
| Slc17a7 | 3.63E-05 | 2.828 |
| Gm37829 | 3.82E-05 | 2.868 |
| Pygb | 3.97E-05 | 2.144 |
| Etfdh | 4.01E-05 | 2.189 |
| Fam174b | 4.04E-05 | 2.445 |
| Ndrg4 | 4.08E-05 | 2.567 |
| Myoz2 | 4.15E-05 | 2.770 |
| Flnc | 4.68E-05 | 2.129 |
| Ccl8 | 4.71E-05 | 0.356 |
| Cox6c | 5.11E-05 | 2.099 |
| Clic5 | 5.28E-05 | 2.297 |
| Atp5a1 | 5.35E-05 | 1.790 |
| H2-M2 | 5.76E-05 | 0.398 |
| Got1 | 5.85E-05 | 2.189 |
| Pilra | 5.94E-05 | 0.415 |
| Nipsnap2 | 7.8E-05 | 2.313 |
| Zbtb16 | 7.85E-05 | 2.329 |
| Gnao1 | 7.96E-05 | 2.329 |
| Kcnj11 | 8.38E-05 | 2.657 |
| Sox9 | 0.000105 | 2.173 |
| Fnbp1l | 0.000115 | 0.503 |
| Hmcn1 | 0.000116 | 2.282 |
| Plce1 | 0.000116 | 2.056 |

| | | |
|---|---|---|
| Adcy5 | 0.000118 | 2.266 |
| Rnf207 | 0.000124 | 2.676 |
| Cd74 | 0.000125 | 0.595 |
| Hspb8 | 0.000126 | 2.497 |
| Acsl1 | 0.000132 | 2.042 |
| Apobec2 | 0.000134 | 2.567 |
| Uqcrfs1 | 0.000137 | 2.114 |
| Asb14 | 0.000142 | 2.567 |
| Acadm | 0.00015 | 1.959 |
| Cox6a2 | 0.000152 | 2.532 |
| Phyh | 0.000153 | 2.266 |
| Pltp | 0.000154 | 0.611 |
| Prelp | 0.000159 | 1.647 |
| Cacnb2 | 0.00016 | 2.639 |
| Actc1 | 0.000162 | 2.497 |
| Muc16 | 0.000164 | 2.567 |
| Grm1 | 0.000165 | 2.532 |
| Rrad | 0.000165 | 2.567 |
| Cd4 | 0.00017 | 0.438 |
| Nppa | 0.000177 | 2.479 |
| A2m | 0.000185 | 2.514 |
| Rassf4 | 0.00019 | 0.503 |
| Acadl | 0.000197 | 1.803 |
| Ednra | 0.000201 | 2.639 |
| Atp5c1 | 0.000202 | 1.959 |
| Tnfrsf11b | 0.000209 | 1.905 |
| Mdh2 | 0.000212 | 1.866 |
| Esrrg | 0.000222 | 2.479 |
| Ccl9 | 0.000222 | 0.518 |
| Hectd2os | 0.000223 | 2.479 |
| Ryr3 | 0.000225 | 2.479 |
| Abcg3 | 0.000227 | 0.470 |
| Tbx20 | 0.00023 | 1.705 |
| Slc16a1 | 0.000232 | 2.282 |
| Pacsin3 | 0.000235 | 2.621 |
| Perp | 0.00024 | 2.603 |
| Slc40a1 | 0.000242 | 0.532 |
| Pgam2 | 0.00025 | 2.428 |
| Cd55 | 0.00026 | 0.607 |
| Yipf7 | 0.000268 | 2.532 |
| Dtna | 0.000279 | 2.549 |
| Il18bp | 0.000285 | 0.559 |
| Uqcrh | 0.000288 | 2.028 |
| Crip2 | 0.000303 | 1.905 |
| Fry | 0.000305 | 1.647 |

| | | |
|---|---|---|
| Hcls1 | 0.000308 | 0.543 |
| Inpp5a | 0.000312 | 1.972 |
| Usp28 | 0.00032 | 2.250 |
| Atp5j | 0.000321 | 1.959 |
| Mfn1 | 0.000329 | 1.853 |
| Gm28661 | 0.000331 | 2.445 |
| Pdk1 | 0.000336 | 2.235 |
| Cd300ld | 0.000337 | 0.582 |
| Vdac1 | 0.000343 | 1.729 |
| Atp5o | 0.000347 | 2.114 |
| Rasgrp1 | 0.000349 | 0.398 |
| C1qa | 0.000357 | 0.599 |
| Ddo | 0.000361 | 2.412 |
| Ddx60 | 0.000379 | 0.486 |
| Wfdc17 | 0.000382 | 0.529 |
| Pdk2 | 0.000383 | 2.428 |
| Bves | 0.000393 | 2.362 |
| Cd86 | 0.000403 | 0.435 |
| Grb14 | 0.000404 | 2.346 |
| Gm45191 | 0.000413 | 0.401 |
| Tlr1 | 0.000424 | 0.432 |
| Efcab2 | 0.000429 | 2.445 |
| Lars2 | 0.000436 | 1.474 |
| Myl2 | 0.000442 | 2.445 |
| Gpx3 | 0.000445 | 2.428 |
| Phkg1 | 0.000458 | 2.497 |
| Abcc9 | 0.000464 | 2.219 |
| Pde1c | 0.000484 | 2.479 |
| Itgb1bp2 | 0.000524 | 2.462 |
| Cped1 | 0.000527 | 1.879 |
| Bank1 | 0.000537 | 0.406 |
| Dpep1 | 0.000546 | 2.346 |
| Car14 | 0.000548 | 2.297 |
| Ppp1r12b | 0.000556 | 1.670 |
| Slc25a4 | 0.00056 | 2.445 |
| Atp9a | 0.00057 | 2.362 |
| Gm28439 | 0.000571 | 2.395 |
| Mmp12 | 0.000573 | 0.616 |
| Basp1 | 0.000574 | 0.582 |
| Kcnq1 | 0.000575 | 2.428 |
| Wdfy3 | 0.000597 | 0.559 |
| Osbpl6 | 0.000599 | 2.329 |
| Cavin4 | 0.000601 | 2.378 |
| Pik3ap1 | 0.000608 | 0.559 |
| Ptprc | 0.000611 | 0.624 |

| | | |
|---|---|---|
| mt-Tm | 0.000622 | 2.346 |
| Lyve1 | 0.000637 | 2.445 |
| Cxadr | 0.00064 | 2.362 |
| Emilin2 | 0.000645 | 0.543 |
| Zbtb20 | 0.000665 | 1.516 |
| Iqgap2 | 0.000682 | 0.607 |
| Pip5k1b | 0.000703 | 2.412 |
| H2-Aa | 0.000722 | 0.586 |
| Ankrd1 | 0.000739 | 2.313 |
| Acat1 | 0.000746 | 1.986 |
| Acot13 | 0.000766 | 2.042 |
| Tagap | 0.000767 | 0.420 |
| Ibsp | 0.000767 | 2.329 |
| Lum | 0.000799 | 1.591 |
| Myh7 | 0.000819 | 2.266 |
| Lyn | 0.000822 | 0.603 |
| Itih5 | 0.000827 | 2.235 |
| Aldh1a1 | 0.000828 | 2.173 |
| Cacna2d1 | 0.000834 | 1.753 |
| Daam2 | 0.000875 | 2.235 |
| Ptprj | 0.000876 | 0.570 |
| Casz1 | 0.000887 | 2.378 |
| Dock2 | 0.000888 | 0.624 |
| Tmsb4x | 0.000898 | 0.651 |
| Maf | 0.000944 | 0.547 |
| Stab2 | 0.000951 | 0.536 |
| Lrrc3b | 0.000963 | 2.189 |
| Fgd2 | 0.000993 | 0.444 |
| Lcp1 | 0.000997 | 0.683 |
| Gm6377 | 0.000998 | 0.451 |
| Fras1 | 0.001009 | 2.219 |
| Mgp | 0.001013 | 1.659 |
| Acadvl | 0.001025 | 1.853 |
| Pcp4l1 | 0.001034 | 2.266 |
| mt-Ti | 0.00104 | 2.297 |
| Kcnh2 | 0.001091 | 2.329 |
| Acp5 | 0.001101 | 0.551 |
| AB124611 | 0.001108 | 0.536 |
| Msr1 | 0.001118 | 0.595 |
| Pdzd2 | 0.001128 | 1.828 |
| Jam2 | 0.00113 | 1.803 |
| Epb41l3 | 0.001133 | 0.624 |
| Dixdc1 | 0.001138 | 2.329 |
| Marcks | 0.001158 | 0.629 |
| Lamb2 | 0.001217 | 1.569 |

| | | |
|---|---|---|
| Maob | 0.001234 | 2.329 |
| Adcy6 | 0.001267 | 1.840 |
| Cyth4 | 0.001274 | 0.629 |
| Csdc2 | 0.001275 | 2.297 |
| Tenm3 | 0.001288 | 2.056 |
| Slc8a1 | 0.001306 | 1.705 |
| Uqcrc1 | 0.001316 | 1.717 |
| Svil | 0.001326 | 1.526 |
| C1qc | 0.00134 | 0.655 |
| C1qb | 0.001344 | 0.683 |
| Pdzrn4 | 0.001351 | 2.129 |
| Plin4 | 0.001358 | 1.919 |
| Coq9 | 0.001409 | 2.014 |
| Nr1d1 | 0.001433 | 1.866 |
| CT572985.1 | 0.001455 | 2.158 |
| Sbk2 | 0.001488 | 2.250 |
| Clec4n | 0.001501 | 0.470 |
| Lrmp | 0.001505 | 0.486 |
| Crispld1 | 0.001532 | 1.892 |
| Snca | 0.001554 | 2.235 |
| Ndufa4 | 0.001554 | 2.042 |
| Dbp | 0.001561 | 2.114 |
| Olfr111 | 0.001611 | 0.493 |
| Sla | 0.001615 | 0.559 |
| Stk39 | 0.001623 | 2.042 |
| Ccdc148 | 0.001652 | 0.540 |
| Rbpms | 0.001655 | 1.765 |
| Sema3d | 0.001669 | 2.250 |
| Cox7a2 | 0.001685 | 1.959 |
| Lcp2 | 0.001693 | 0.559 |
| Pkp2 | 0.001694 | 2.158 |
| Fxyd1 | 0.0017 | 2.173 |
| Dlc1 | 0.001709 | 1.717 |
| Cycs | 0.001723 | 2.014 |
| Itga7 | 0.001762 | 2.219 |
| Egln1 | 0.001762 | 1.840 |
| Ndufv2 | 0.001871 | 1.879 |
| Arl4a | 0.001882 | 2.056 |
| Pkia | 0.001889 | 2.189 |
| Ear2 | 0.001904 | 0.603 |
| Hcn1 | 0.001904 | 2.144 |
| Clu | 0.001919 | 1.625 |
| Uqcrc2 | 0.001964 | 1.778 |
| Col6a3 | 0.001982 | 1.444 |
| Abca9 | 0.001987 | 0.543 |

| | | |
|---|---|---|
| Hydin | 0.002045 | 2.158 |
| Aldh1l2 | 0.002058 | 2.129 |
| Ndufs1 | 0.002074 | 1.741 |
| C3 | 0.002109 | 1.495 |
| Cs | 0.002186 | 1.613 |
| Myo1b | 0.00219 | 0.518 |
| Tmtc1 | 0.00219 | 1.625 |
| Myo9a | 0.002191 | 0.678 |
| Btnl9 | 0.00228 | 2.028 |
| Myl9 | 0.002339 | 1.892 |
| Tbxas1 | 0.002377 | 0.633 |
| Tpm1 | 0.002377 | 2.219 |
| Lst1 | 0.002389 | 0.454 |
| Gm45531 | 0.002397 | 2.014 |
| Gpihbp1 | 0.002406 | 2.028 |
| Synpo2 | 0.002437 | 1.765 |
| Prox1 | 0.002489 | 1.932 |
| Peg3 | 0.002506 | 1.986 |
| Shtn1 | 0.002541 | 0.620 |
| Slc9a9 | 0.002567 | 0.521 |
| Rassf2 | 0.002593 | 0.570 |
| Syde2 | 0.002616 | 2.189 |
| Fat3 | 0.002633 | 0.500 |
| Acaa2 | 0.002643 | 1.866 |
| Tuba4a | 0.002696 | 2.042 |
| Jph2 | 0.002726 | 2.014 |
| Gm15564 | 0.002779 | 1.454 |
| Gm13054 | 0.002823 | 2.099 |
| Lrrc14b | 0.002882 | 2.173 |
| 3425401B1 | 0.002927 | 2.173 |
| Ly9 | 0.002992 | 0.646 |
| Ptgds | 0.003014 | 1.972 |
| mt-Co2 | 0.003051 | 2.056 |
| Hfe | 0.003073 | 0.563 |
| Trp53inp2 | 0.003132 | 1.625 |
| Gsta4 | 0.003152 | 2.158 |
| Smarcd3 | 0.003158 | 2.085 |
| Synpo2l | 0.003221 | 1.945 |
| Rfxank | 0.003251 | 2.114 |
| 4930523C0 | 0.00327 | 1.602 |
| H2-Ab1 | 0.003304 | 0.629 |
| Fabp3-ps1 | 0.003312 | 2.000 |
| Pld4 | 0.003325 | 0.525 |
| Dhrs7c | 0.003342 | 1.959 |
| Entpd5 | 0.003365 | 1.919 |

| | | |
|---|---|---|
| Vav1 | 0.003375 | 0.611 |
| Hhatl | 0.003402 | 1.959 |
| Fermt2 | 0.003478 | 1.591 |
| Hmox1 | 0.003496 | 0.674 |
| AC123062.1 | 0.003504 | 0.480 |
| Prkag2 | 0.003527 | 1.815 |
| Cd83 | 0.003568 | 0.559 |
| Trabd2b | 0.003601 | 1.765 |
| Scd1 | 0.003709 | 0.664 |
| Ano4 | 0.003734 | 1.972 |
| Hopx | 0.003751 | 2.129 |
| Fmo2 | 0.003791 | 1.659 |
| Atp5g1 | 0.00386 | 2.099 |
| Grn | 0.00387 | 0.702 |
| Fyco1 | 0.003894 | 1.516 |
| Ly86 | 0.003894 | 0.582 |
| Tln2 | 0.003898 | 1.717 |
| Sspn | 0.003981 | 1.879 |
| C4b | 0.003999 | 1.625 |
| Ogn | 0.004066 | 2.042 |
| Hpgds | 0.004076 | 0.642 |
| D10Jhu81e | 0.004132 | 2.000 |
| B3galnt1 | 0.004169 | 0.543 |
| Ndufb5 | 0.00417 | 1.853 |
| Pdlim5 | 0.004178 | 1.495 |
| Gm13341 | 0.004181 | 2.085 |
| Abca1 | 0.004301 | 0.732 |
| Ogdh | 0.004398 | 1.444 |
| Abra | 0.004454 | 2.042 |
| Rasgef1b | 0.004518 | 0.629 |
| Gucy1a3 | 0.004636 | 1.932 |
| Fam180a | 0.004655 | 2.085 |
| Tppp | 0.00469 | 2.085 |
| Cd180 | 0.004697 | 0.629 |
| H2-Eb1 | 0.004727 | 0.620 |
| Rab8b | 0.004768 | 0.688 |
| Frzb | 0.00484 | 1.602 |
| Cfp | 0.00489 | 0.611 |
| Ablim3 | 0.004905 | 2.014 |
| Lair1 | 0.004976 | 0.669 |
| Irs1 | 0.004979 | 2.056 |
| Evl | 0.004989 | 0.669 |
| Sema3c | 0.005003 | 2.014 |
| Ifi207 | 0.00502 | 0.678 |
| Uqcrb | 0.005038 | 1.753 |

| | | |
|---|---|---|
| Rarb | 0.005063 | 2.000 |
| Apbb1ip | 0.005099 | 0.595 |
| Arhgap4 | 0.005115 | 0.497 |
| Mcoln3 | 0.005127 | 0.540 |
| 2310043P1 | 0.005157 | 0.511 |
| Ano1 | 0.005176 | 1.647 |
| Bdh1 | 0.005241 | 1.919 |
| Mrgpre | 0.005276 | 0.497 |
| Fkbp3 | 0.005347 | 1.803 |
| Atp8a1 | 0.00535 | 0.664 |
| Sync | 0.005384 | 2.000 |
| Aldh5a1 | 0.005398 | 2.056 |
| Themis2 | 0.005446 | 0.590 |
| Spock2 | 0.005472 | 2.014 |
| Itga8 | 0.005499 | 1.636 |
| Angptl7 | 0.005533 | 1.959 |
| Gm43185 | 0.005601 | 2.056 |
| Sucla2 | 0.005648 | 1.682 |
| Serpine2 | 0.005664 | 1.505 |
| Pla2g5 | 0.00568 | 1.892 |
| mt-Nd4l | 0.005732 | 1.892 |
| Rgs5 | 0.005746 | 1.840 |
| Ildr2 | 0.0058 | 1.705 |
| Tead1 | 0.005802 | 1.580 |
| Pink1 | 0.005881 | 1.682 |
| Ptpn6 | 0.00591 | 0.611 |
| Cilp2 | 0.006038 | 2.042 |
| Stap1 | 0.006121 | 0.525 |
| Timp3 | 0.006162 | 1.454 |
| Tnxb | 0.006165 | 1.879 |
| Tuba8 | 0.006179 | 1.840 |
| Homer1 | 0.006185 | 1.682 |
| Sirpa | 0.006236 | 0.722 |
| Fxyd6 | 0.006283 | 2.014 |
| Fabp4 | 0.006305 | 1.613 |
| Ap1s2 | 0.006305 | 0.620 |
| Csf1r | 0.006358 | 0.669 |
| Ndufs4 | 0.006361 | 1.636 |
| Nov | 0.006373 | 1.693 |
| Atcayos | 0.006414 | 1.879 |
| Rps6ka1 | 0.006454 | 0.595 |
| Fh1 | 0.006454 | 1.866 |
| Ndufb10 | 0.006505 | 1.659 |
| Apoe | 0.006509 | 0.712 |
| Clec4a3 | 0.006569 | 0.529 |

| | | |
|---|---|---|
| Gbp3 | 0.006571 | 0.595 |
| Chchd3 | 0.006646 | 1.729 |
| Ndufb11 | 0.006668 | 1.815 |
| S100a1 | 0.006694 | 1.613 |
| Atp5k | 0.006749 | 1.765 |
| Sphkap | 0.006754 | 1.828 |
| Clec4a2 | 0.006759 | 0.493 |
| Magi3 | 0.006831 | 1.705 |
| Serpina3g | 0.006949 | 0.595 |
| Coro1a | 0.00701 | 0.611 |
| Cnksr3 | 0.007101 | 0.543 |
| Actb | 0.007135 | 0.737 |
| Syt7 | 0.007182 | 1.905 |
| Anxa8 | 0.007186 | 1.625 |
| Camta1 | 0.0072 | 1.879 |
| Tcf21 | 0.007215 | 2.014 |
| Atp5j2 | 0.007278 | 1.840 |
| Cd59a | 0.007285 | 1.682 |
| Cspg4 | 0.007302 | 1.537 |
| Ccnd2 | 0.007319 | 1.569 |
| Col6a1 | 0.007364 | 1.414 |
| Tnc | 0.007524 | 1.558 |
| Col14a1 | 0.007545 | 1.464 |
| Cox4i1 | 0.007556 | 1.636 |
| Nckap1l | 0.007583 | 0.712 |
| Gramd1b | 0.0077 | 0.651 |
| Tfpi | 0.007756 | 1.670 |
| Cotl1 | 0.007759 | 0.697 |
| Ech1 | 0.007795 | 1.693 |
| Ms4a7 | 0.007803 | 0.678 |
| Pcsk6 | 0.007828 | 1.959 |
| Stab1 | 0.007895 | 0.697 |
| P2ry14 | 0.007913 | 0.603 |
| Hba-a1 | 0.007922 | 1.892 |
| Bnip3 | 0.008013 | 1.945 |
| Cox5b | 0.008095 | 1.705 |
| Nfia | 0.008119 | 1.516 |
| Scn4a | 0.008258 | 1.866 |
| Hspb7 | 0.008291 | 1.986 |
| Cttnbp2nl | 0.008355 | 0.678 |
| Dynll2 | 0.008366 | 1.670 |
| Ndufs2 | 0.008386 | 1.569 |
| Sez6l2 | 0.008439 | 0.518 |
| Acyp2 | 0.008471 | 1.986 |
| Oxnad1 | 0.008563 | 1.945 |

| | | |
|---|---|---|
| Cd2ap | 0.008579 | 0.655 |
| Synm | 0.008659 | 1.591 |
| Hexb | 0.00866 | 0.727 |
| Cacna1d | 0.008726 | 0.595 |
| Obsl1 | 0.008773 | 1.919 |
| Arhgap15 | 0.008796 | 0.582 |
| Myocd | 0.008841 | 1.853 |
| Igsf23 | 0.008852 | 1.778 |
| Ikzf1 | 0.00888 | 0.624 |
| Tpm2 | 0.009044 | 1.558 |
| C130080G1 | 0.009147 | 1.778 |
| Ppara | 0.009147 | 1.778 |
| Sln | 0.009239 | 1.753 |
| Apol11b | 0.009278 | 1.892 |
| C530008M | 0.009299 | 1.932 |
| Myo5b | 0.009381 | 1.765 |
| Adamts8 | 0.009396 | 1.892 |
| Dpf3 | 0.009412 | 1.840 |
| mt-Nd2 | 0.009454 | 1.959 |
| Tgfbr3 | 0.009481 | 1.647 |
| Ddx3y | 0.009522 | 0.669 |
| Cxcl10 | 0.00959 | 0.532 |
| Plekha6 | 0.009607 | 1.815 |
| Cfl2 | 0.009685 | 1.682 |
| Asb15 | 0.009709 | 1.765 |
| Ptbp3 | 0.009709 | 0.722 |
| Slfn2 | 0.009743 | 0.683 |
| Rilpl1 | 0.009768 | 1.778 |
| Ccl3 | 0.009776 | 0.555 |
| Ndrg1 | 0.00992 | 0.629 |
| Fcer1g | 0.009952 | 0.693 |
| Kif21a | 0.009986 | 1.840 |
| Chst2 | 0.009988 | 0.563 |
| Slc5a3 | 0.010014 | 1.647 |
| Neo1 | 0.010033 | 1.569 |
| Gm42418 | 0.01014 | 1.329 |
| Ablim2 | 0.010307 | 1.932 |
| Arap1 | 0.01032 | 0.664 |
| Angptl2 | 0.010438 | 1.580 |
| Plekha7 | 0.010482 | 1.945 |
| Pid1 | 0.010513 | 0.629 |
| Ndufa8 | 0.010515 | 1.741 |
| Samsn1 | 0.01054 | 0.540 |
| Ndufa11 | 0.010726 | 1.840 |
| Fam49a | 0.010805 | 0.646 |

| | | |
|---|---|---|
| Unc93b1 | 0.010838 | 0.678 |
| Sort1 | 0.010882 | 1.454 |
| Kcnd2 | 0.011013 | 1.741 |
| Kcnb1 | 0.011044 | 1.932 |
| Pik3cg | 0.011234 | 0.590 |
| Rgl1 | 0.011236 | 0.693 |
| Gstm2 | 0.011237 | 1.828 |
| Mmp8 | 0.011397 | 0.555 |
| Itgam | 0.011493 | 0.651 |
| Schip1 | 0.011518 | 1.932 |
| Serpina3n | 0.011664 | 1.347 |
| Bola3 | 0.011749 | 1.803 |
| Isca1 | 0.011835 | 1.753 |
| Slit3 | 0.01186 | 1.670 |
| Atp5d | 0.011871 | 1.625 |
| Mmp27 | 0.011947 | 0.566 |
| Pik3cd | 0.012045 | 0.651 |
| Asb10 | 0.012049 | 1.717 |
| Zfp385b | 0.012049 | 1.717 |
| Blnk | 0.012067 | 0.655 |
| D330045A2 | 0.012166 | 0.566 |
| Trak2 | 0.012202 | 1.424 |
| Sox6 | 0.012301 | 1.717 |
| Apod | 0.012331 | 1.840 |
| Limd2 | 0.012356 | 0.529 |
| Grk2 | 0.012399 | 0.678 |
| Lima1 | 0.012477 | 0.693 |
| Lacc1 | 0.012495 | 0.620 |
| Pik3ip1 | 0.012668 | 1.905 |
| Ifit1 | 0.012706 | 0.559 |
| C7 | 0.012706 | 1.705 |
| Inpp5d | 0.012735 | 0.688 |
| Slamf9 | 0.012964 | 0.532 |
| Uqcr10 | 0.012997 | 1.753 |
| mt-Cytb | 0.013034 | 1.905 |
| Vegfb | 0.013073 | 1.693 |
| Dlat | 0.013081 | 1.636 |
| Cd38 | 0.0131 | 0.674 |
| Dock10 | 0.013135 | 0.707 |
| Hist1h1d | 0.013168 | 0.607 |
| Slc25a12 | 0.01335 | 1.602 |
| Lynx1 | 0.013392 | 1.729 |
| Sgca | 0.013396 | 1.853 |
| Gm10435 | 0.013397 | 1.705 |
| Gm5532 | 0.013397 | 1.705 |

| | | |
|---|---|---|
| Slc25a23 | 0.013472 | 1.879 |
| mt-Tq | 0.01351 | 1.741 |
| Slc25a3 | 0.013586 | 1.414 |
| Smtn | 0.01362 | 1.485 |
| Strip2 | 0.013651 | 1.866 |
| Cand2 | 0.013653 | 1.905 |
| Atp5f1 | 0.013729 | 1.516 |
| Ccng1 | 0.013807 | 1.516 |
| Abcc3 | 0.013843 | 0.693 |
| Fam49b | 0.013855 | 0.664 |
| Sdc3 | 0.013888 | 0.742 |
| Igsf6 | 0.013967 | 0.683 |
| A130071D0 | 0.014004 | 0.629 |
| Fosb | 0.014026 | 1.892 |
| Pm20d2 | 0.014134 | 1.892 |
| Sfrp1 | 0.01424 | 0.655 |
| Uqcrq | 0.014259 | 1.705 |
| Bmp2k | 0.014362 | 0.674 |
| Pfn1 | 0.014521 | 0.722 |
| Tlr7 | 0.014534 | 0.669 |
| Slco3a1 | 0.014543 | 1.537 |
| Bach2os | 0.014616 | 0.586 |
| Pdgfrb | 0.01463 | 1.495 |
| Actr3 | 0.014655 | 0.742 |
| Rabgap1l | 0.014672 | 1.495 |
| Ccl2 | 0.014685 | 0.599 |
| Mmp2 | 0.014711 | 1.444 |
| mt-Tp | 0.014752 | 1.790 |
| Gatm | 0.014826 | 0.590 |
| Cybb | 0.014879 | 0.747 |
| Tecr | 0.014935 | 1.693 |
| Plod2 | 0.014987 | 1.548 |
| Serp1 | 0.015012 | 0.646 |
| Sh3bp5 | 0.01503 | 0.595 |
| Kdr | 0.01507 | 1.613 |
| Reck | 0.015201 | 1.591 |
| Gys1 | 0.015208 | 1.778 |
| Ccdc157 | 0.015253 | 1.803 |
| Stard9 | 0.01531 | 0.683 |
| Prss23 | 0.01532 | 1.659 |
| Mical3 | 0.015336 | 1.485 |
| Kcnj10 | 0.015393 | 0.532 |
| Lipg | 0.015412 | 0.555 |
| Ncapg2 | 0.01545 | 0.637 |
| Msi2 | 0.015459 | 1.444 |

| | | |
|---|---|---|
| Adamtsl3 | 0.015493 | 1.526 |
| Hivep3 | 0.015533 | 0.646 |
| Irf5 | 0.015546 | 0.664 |
| Cndp2 | 0.015554 | 0.717 |
| A630001G2 | 0.015642 | 0.570 |
| Decr1 | 0.015671 | 1.705 |
| Tcea3 | 0.015719 | 1.853 |
| Nap1l5 | 0.015721 | 1.670 |
| Acad11 | 0.015739 | 1.682 |
| Zfp9 | 0.015772 | 1.693 |
| Ebi3 | 0.015805 | 0.566 |
| Adam17 | 0.015858 | 0.712 |
| Met | 0.015907 | 0.582 |
| 5430419D1 | 0.01592 | 1.670 |
| Adarb1 | 0.01603 | 1.454 |
| Gstk1 | 0.01606 | 1.879 |
| Rxfp1 | 0.016095 | 1.670 |
| Dcn | 0.016139 | 1.347 |
| Atp13a2 | 0.016268 | 0.678 |
| Ndufa12 | 0.016288 | 1.741 |
| Plagl2 | 0.016301 | 0.607 |
| Dnm1 | 0.016327 | 1.729 |
| Pdzrn3 | 0.016375 | 1.558 |
| Mdh1 | 0.016396 | 1.866 |
| Des | 0.016508 | 1.659 |
| Myl7 | 0.016521 | 1.647 |
| Fsd1l | 0.016542 | 1.790 |
| Cep250 | 0.016548 | 0.683 |
| Gcnt1 | 0.016588 | 0.664 |
| Enpep | 0.016593 | 1.705 |
| Nkx2-5 | 0.016605 | 1.705 |
| Atp5h | 0.016695 | 1.602 |
| Bzw2 | 0.016726 | 1.729 |
| Ptn | 0.01683 | 1.853 |
| Zfp651 | 0.016897 | 1.741 |
| Gm45221 | 0.016962 | 0.586 |
| Nlrp1b | 0.016971 | 0.590 |
| Gm44745 | 0.016977 | 0.540 |
| Slc16a6 | 0.017149 | 0.559 |
| Tnfrsf26 | 0.01722 | 0.555 |
| mt-Nd4 | 0.017311 | 1.853 |
| Bche | 0.017517 | 1.647 |
| Smpx | 0.017547 | 1.636 |
| Ndufv1 | 0.01761 | 1.636 |
| Arhgef9 | 0.017645 | 1.840 |

| | | |
|---|---|---|
| mt-Tw | 0.017684 | 1.705 |
| Lpxn | 0.017726 | 0.678 |
| Dgat2 | 0.017749 | 1.840 |
| Suclg1 | 0.017839 | 1.591 |
| Abi3 | 0.01802 | 0.660 |
| Lpcat2 | 0.018146 | 0.629 |
| Ptpn11 | 0.018181 | 1.505 |
| Mtus2 | 0.018191 | 1.853 |
| Fkbp4 | 0.0182 | 1.516 |
| Myzap | 0.018422 | 1.778 |
| Tmem109 | 0.018487 | 1.647 |
| Marveld1 | 0.018513 | 1.495 |
| Ndufa3 | 0.018547 | 1.705 |
| Pkig | 0.018621 | 1.682 |
| Ar | 0.018647 | 1.840 |
| Svip | 0.018671 | 1.840 |
| Ncf1 | 0.018741 | 0.637 |
| Enah | 0.01895 | 1.602 |
| Aspn | 0.01901 | 1.840 |
| Cnst | 0.019013 | 1.613 |
| Hadhb | 0.019173 | 1.591 |
| Gm8995 | 0.01929 | 0.737 |
| Wdfy4 | 0.01934 | 0.732 |
| Ifit2 | 0.019424 | 0.616 |
| Dusp6 | 0.019471 | 0.664 |
| Mir133a-1h | 0.019507 | 1.636 |
| Slfn8 | 0.019654 | 0.642 |
| Sptbn1 | 0.019684 | 1.320 |
| Abcg1 | 0.019788 | 0.763 |
| Spi1 | 0.019886 | 0.688 |
| Bmp3 | 0.019926 | 1.625 |
| 9330158H0 | 0.019939 | 1.625 |
| Pgm2l1 | 0.020052 | 0.642 |
| Pyurf | 0.020095 | 1.828 |
| Fgd4 | 0.020221 | 0.722 |
| Bex1 | 0.02027 | 1.625 |
| Tank | 0.020296 | 0.697 |
| Osgepl1 | 0.020466 | 1.803 |
| Syk | 0.020534 | 0.737 |
| Mrps23 | 0.020694 | 1.741 |
| Slc8b1 | 0.020769 | 0.646 |
| Tfdp2 | 0.020786 | 1.613 |
| Mrpl42 | 0.02079 | 1.636 |
| Arpc4 | 0.020852 | 0.717 |
| Nlrp10 | 0.020905 | 1.753 |

| | | |
|---|---|---|
| Cdh13 | 0.02091 | 1.385 |
| Cep128 | 0.020919 | 0.629 |
| Gm10222 | 0.021181 | 1.828 |
| Eif2ak2 | 0.02136 | 0.683 |
| Fcgr1 | 0.021458 | 0.664 |
| Selenbp1 | 0.021536 | 1.803 |
| Htra3 | 0.021714 | 1.693 |
| Pdhb | 0.02184 | 1.613 |
| Npy | 0.021918 | 1.659 |
| Slc43a2 | 0.021923 | 0.697 |
| Samhd1 | 0.021986 | 0.717 |
| mt-Co1 | 0.022005 | 1.815 |
| mt-Nd1 | 0.022018 | 1.815 |
| Zfp217 | 0.022043 | 0.660 |
| Ehd1 | 0.022237 | 0.642 |
| Mlana | 0.022317 | 1.705 |
| Mir6236 | 0.022378 | 1.301 |
| Rab3il1 | 0.022381 | 0.688 |
| Nedd4 | 0.022435 | 1.376 |
| Ksr2 | 0.022443 | 0.599 |
| Scara3 | 0.022521 | 1.444 |
| Meis1 | 0.022523 | 1.753 |
| Dclre1c | 0.022673 | 0.629 |
| 1810011O1 | 0.022686 | 1.815 |
| Myo15b | 0.022763 | 0.595 |
| Cyfip1 | 0.022788 | 0.742 |
| C1s1 | 0.022883 | 1.495 |
| Nfasc | 0.023018 | 1.693 |
| P2rx7 | 0.023145 | 0.683 |
| Picalm | 0.023188 | 0.768 |
| Homer2 | 0.023229 | 1.659 |
| Psip1 | 0.023232 | 1.464 |
| Tmem26 | 0.023288 | 0.620 |
| Stk17b | 0.023326 | 0.669 |
| Ghitm | 0.023419 | 1.444 |
| D130040H2 | 0.023587 | 0.555 |
| Tlr8 | 0.023681 | 0.717 |
| Kcnd3 | 0.023681 | 1.591 |
| Tmem158 | 0.023779 | 0.559 |
| Efhd2 | 0.023783 | 0.732 |
| Csf2ra | 0.023803 | 0.702 |
| Fam110c | 0.023873 | 0.563 |
| Ptpn1 | 0.024029 | 0.732 |
| Itih2 | 0.024036 | 1.765 |
| Adam33 | 0.024157 | 1.682 |

| | | |
|---|---|---|
| mt-Nd5 | 0.024182 | 1.803 |
| Mylip | 0.024274 | 0.747 |
| Slc35g1 | 0.024317 | 0.570 |
| Clec7a | 0.024411 | 0.763 |
| Ndufa13 | 0.024583 | 1.636 |
| Ppm1h | 0.024598 | 0.693 |
| Fgr | 0.024947 | 0.642 |
| Fbxl22 | 0.024959 | 1.790 |
| Ecm2 | 0.025036 | 1.670 |
| Trerf1 | 0.025081 | 0.616 |
| Kcnj2 | 0.025124 | 1.729 |
| Atp5g3 | 0.025334 | 1.548 |
| Card14 | 0.025369 | 1.753 |
| Kcnip2 | 0.0255 | 1.580 |
| Mov10l1 | 0.0255 | 1.580 |
| Ighg2c | 0.025504 | 1.580 |
| Fyn | 0.025667 | 0.664 |
| Egr1 | 0.025788 | 1.505 |
| Eps8l2 | 0.025822 | 1.778 |
| Znrf2 | 0.025826 | 0.688 |
| Csf3r | 0.025891 | 0.595 |
| Ccdc50 | 0.025922 | 0.717 |
| Sdhb | 0.02596 | 1.516 |
| Ms4a6d | 0.026314 | 0.717 |
| Mrps6 | 0.026334 | 1.717 |
| Mefv | 0.026341 | 0.651 |
| Mpdz | 0.026352 | 1.505 |
| Scara5 | 0.026371 | 1.765 |
| Fgf2 | 0.026414 | 1.505 |
| Pkd1 | 0.026423 | 1.347 |
| Ednrb | 0.02677 | 1.717 |
| Evi2a | 0.026825 | 0.683 |
| Mov10 | 0.026851 | 0.578 |
| Klf15 | 0.026919 | 1.778 |
| Dab2 | 0.026957 | 0.712 |
| Mgst3 | 0.026969 | 1.765 |
| Cxcl13 | 0.026997 | 1.569 |
| Fmn1 | 0.027033 | 0.742 |
| Myc | 0.027234 | 0.624 |
| Dgki | 0.027297 | 0.570 |
| Zfhx4 | 0.027436 | 1.778 |
| Flrt2 | 0.027468 | 1.625 |
| Coro1b | 0.027475 | 0.678 |
| Bag2 | 0.027619 | 1.778 |
| Lims2 | 0.027638 | 1.741 |

| | | |
|---|---|---|
| Apobec3 | 0.027675 | 0.642 |
| Fhl3 | 0.027677 | 0.566 |
| Dhtkd1 | 0.02768 | 1.659 |
| Hadh | 0.027722 | 1.580 |
| Bsg | 0.027786 | 1.404 |
| Tarsl2 | 0.027805 | 1.705 |
| Camk1d | 0.027875 | 0.646 |
| Tns2 | 0.027977 | 1.474 |
| Atp6v0a4 | 0.028024 | 1.717 |
| Mafb | 0.028106 | 0.732 |
| Gm10925 | 0.028141 | 1.765 |
| Scrn1 | 0.028156 | 1.647 |
| Pea15a | 0.028168 | 0.732 |
| Igfbp6 | 0.028235 | 1.602 |
| Uty | 0.028357 | 0.664 |
| Ndufb7 | 0.028479 | 1.537 |
| Agmo | 0.028485 | 0.563 |
| Ndufs8 | 0.028558 | 1.625 |
| Dag1 | 0.028623 | 1.376 |
| Ppp1r3a | 0.028715 | 1.569 |
| Fgfrl1 | 0.028728 | 1.682 |
| Fcrls | 0.028818 | 0.570 |
| Msrb3 | 0.028904 | 1.414 |
| Rtn2 | 0.029229 | 1.670 |
| Ccser2 | 0.029334 | 1.414 |
| Il1rn | 0.029337 | 0.712 |
| mt-Rnr1 | 0.029406 | 1.765 |
| Slc25a10 | 0.029505 | 0.578 |
| Hsdl2 | 0.02951 | 1.537 |
| Upk3b | 0.029563 | 1.602 |
| Adpgk | 0.029569 | 0.651 |
| Stk32a | 0.029596 | 1.591 |
| Itih5l-ps | 0.029677 | 1.693 |
| Plxdc2 | 0.029709 | 1.454 |
| Hlf | 0.029813 | 1.705 |
| Gm42047 | 0.029857 | 0.590 |
| Map1lc3a | 0.029893 | 1.741 |
| Taok3 | 0.029929 | 0.742 |
| Oxct1 | 0.02998 | 1.357 |
| Zyx | 0.030083 | 0.712 |
| Prdx3 | 0.03013 | 1.591 |
| Fgf13 | 0.030242 | 1.729 |
| Zfp992 | 0.030274 | 0.570 |
| Cap1 | 0.030274 | 0.774 |
| Hmgb3 | 0.03033 | 1.741 |

| | | |
|---|---|---|
| Slc20a2 | 0.030378 | 1.591 |
| Idh3b | 0.030399 | 1.526 |
| Dock8 | 0.030481 | 0.697 |
| Ikbke | 0.030807 | 0.633 |
| Gprc5b | 0.031095 | 1.526 |
| Dchs2 | 0.031219 | 1.753 |
| Ptpn13 | 0.031227 | 1.753 |
| Tanc2 | 0.031345 | 0.732 |
| Pik3r5 | 0.031475 | 0.742 |
| Arhgap12 | 0.031522 | 0.683 |
| Sh2d1b1 | 0.031713 | 0.570 |
| Gpam | 0.031726 | 1.613 |
| 8430408G2 | 0.031748 | 1.613 |
| Slc28a2 | 0.031783 | 0.582 |
| Pfkfb3 | 0.031788 | 0.693 |
| mt-Atp6 | 0.031867 | 1.753 |
| Bbx | 0.031904 | 0.717 |
| Adgrg2 | 0.031934 | 1.729 |
| Ncf2 | 0.031968 | 0.717 |
| Sema3a | 0.032013 | 1.741 |
| Prepl | 0.032108 | 1.548 |
| Kcnj5 | 0.032145 | 1.548 |
| Dpt | 0.032172 | 1.434 |
| Scimp | 0.032251 | 0.586 |
| Slc11a1 | 0.032318 | 0.737 |
| Stk10 | 0.032325 | 0.732 |
| Tmem176b | 0.032353 | 0.669 |
| Got2 | 0.03262 | 1.591 |
| Acsl4 | 0.032843 | 0.712 |
| Dnajc13 | 0.032894 | 0.763 |
| Adra1b | 0.032927 | 1.602 |
| Serping1 | 0.032939 | 1.434 |
| Nrxn2 | 0.033011 | 0.607 |
| Fmod | 0.033017 | 1.454 |
| Aif1l | 0.033019 | 1.613 |
| Mrpl18 | 0.033036 | 1.537 |
| Etv1 | 0.033466 | 1.693 |
| Ndufa10 | 0.033478 | 1.474 |
| Prim1 | 0.033479 | 0.590 |
| Csk | 0.033679 | 0.697 |
| Ndufa9 | 0.033857 | 1.537 |
| Plxnc1 | 0.0339 | 0.732 |
| Ms4a14 | 0.034079 | 0.722 |
| Arhgap18 | 0.034205 | 0.722 |
| Nox4 | 0.034226 | 1.505 |

| | | |
|---|---|---|
| 3632451O0 | 0.034253 | 1.729 |
| Tlr9 | 0.034262 | 0.595 |
| n-R5s194 | 0.03429 | 1.717 |
| Pecam1 | 0.03435 | 0.774 |
| Itgbl1 | 0.034434 | 1.434 |
| Col6a2 | 0.034503 | 1.329 |
| Grb2 | 0.034582 | 0.737 |
| Chchd7 | 0.034868 | 1.729 |
| P2ry6 | 0.034997 | 0.669 |
| Fads3 | 0.035172 | 1.474 |
| Slc4a4 | 0.035254 | 1.729 |
| Fam107b | 0.035389 | 0.693 |
| Gbp5 | 0.035503 | 0.607 |
| Fyb | 0.035593 | 0.582 |
| Ptgfrn | 0.03578 | 1.376 |
| Crat | 0.035899 | 1.444 |
| Bpgm | 0.035917 | 1.558 |
| Sh3bgrl | 0.036071 | 0.758 |
| Gm15675 | 0.036117 | 0.629 |
| Ptpn22 | 0.036287 | 0.637 |
| Usp8 | 0.036291 | 0.758 |
| Gm29216 | 0.036374 | 1.717 |
| Tacc2 | 0.036432 | 1.485 |
| Bhlhe40 | 0.036479 | 1.338 |
| Ndufb8 | 0.036857 | 1.613 |
| Syt4 | 0.037198 | 1.580 |
| Hck | 0.037367 | 0.633 |
| Twsg1 | 0.037674 | 1.404 |
| Neurl3 | 0.03773 | 0.655 |
| Lama2 | 0.037748 | 1.602 |
| Chmp4c | 0.037775 | 1.717 |
| Irak4 | 0.038073 | 0.697 |
| Ndufa6 | 0.038109 | 1.548 |
| Pabpc4 | 0.038188 | 1.526 |
| Aldh6a1 | 0.03824 | 1.548 |
| Syn1 | 0.038384 | 0.629 |
| Abcd2 | 0.038484 | 0.655 |
| Arpc5 | 0.038529 | 0.742 |
| Cisd1 | 0.03853 | 1.670 |
| Fbxo32 | 0.038682 | 1.283 |
| Gm32849 | 0.03883 | 1.602 |
| Bod1 | 0.038868 | 1.670 |
| Swap70 | 0.039033 | 0.722 |
| Gdf10 | 0.039162 | 1.717 |
| Cdh2 | 0.039241 | 1.464 |

| | | |
|---|---|---|
| Rab3b | 0.039384 | 1.569 |
| Gbp8 | 0.039431 | 0.582 |
| Pkib | 0.039442 | 0.678 |
| Zfp758 | 0.039445 | 0.611 |
| Cavin2 | 0.039486 | 1.485 |
| Insr | 0.039555 | 1.385 |
| Cdc42ep3 | 0.039636 | 1.537 |
| Xrcc1 | 0.039713 | 0.620 |
| Fgd3 | 0.039727 | 0.651 |
| Cmip | 0.040016 | 0.727 |
| Ahnak | 0.040023 | 1.292 |
| Hspb1 | 0.040074 | 1.602 |
| Dok2 | 0.040108 | 0.586 |
| Egln2 | 0.040279 | 0.620 |
| Aldoa | 0.040325 | 1.320 |
| Nfam1 | 0.040402 | 0.651 |
| Fes | 0.040455 | 0.702 |
| Nlrp1c-ps | 0.040659 | 0.603 |
| Sema4b | 0.040664 | 0.642 |
| Pgd | 0.040671 | 0.737 |
| AI662270 | 0.040788 | 0.633 |
| Ryk | 0.041233 | 1.548 |
| Gas7 | 0.041251 | 0.678 |
| Cpd | 0.041357 | 0.768 |
| Serpinf1 | 0.041464 | 1.516 |
| Kctd9 | 0.04147 | 1.516 |
| Hspb2 | 0.041963 | 1.647 |
| Nckap1 | 0.042054 | 1.376 |
| Tom1l2 | 0.042129 | 1.464 |
| Ivd | 0.042167 | 1.548 |
| Laptm5 | 0.042194 | 0.779 |
| Mapk12 | 0.042241 | 1.705 |
| Cacna1b | 0.042323 | 0.607 |
| Ppip5k1 | 0.042564 | 1.464 |
| Car8 | 0.042613 | 0.664 |
| Pcolce | 0.042817 | 1.474 |
| Plcl2 | 0.042827 | 0.737 |
| Bgn | 0.042969 | 1.320 |
| Fhl1 | 0.043137 | 1.454 |
| Tef | 0.043142 | 1.454 |
| Cfl1 | 0.043463 | 0.790 |
| Ndufb2 | 0.043517 | 1.613 |
| Ccdc88b | 0.043574 | 0.633 |
| 5430427O1 | 0.043612 | 0.616 |
| mt-Rnr2 | 0.043684 | 1.693 |

| | | |
|---|---|---|
| Serpinb6a | 0.0437 | 0.779 |
| Gpx1 | 0.043956 | 0.790 |
| Zfyve21 | 0.043983 | 1.693 |
| Zfp106 | 0.04411 | 1.292 |
| AI464131 | 0.044123 | 1.548 |
| Arhgdib | 0.044306 | 0.727 |
| Cobll1 | 0.044366 | 1.526 |
| Fam220a | 0.044426 | 1.613 |
| Map3k20 | 0.044531 | 1.310 |
| Narf | 0.044532 | 1.516 |
| C5ar1 | 0.044991 | 0.697 |
| CT010467.1 | 0.045126 | 1.320 |
| Sorbs2 | 0.045141 | 1.434 |
| Mxra8 | 0.045228 | 1.434 |
| Ticam2 | 0.045269 | 0.599 |
| Syn2 | 0.045348 | 1.569 |
| Ss18l2 | 0.045613 | 1.682 |
| Klhl13 | 0.045648 | 1.647 |
| Inpp5j | 0.045662 | 1.636 |
| Copz2 | 0.045739 | 1.526 |
| Dld | 0.046189 | 1.395 |
| Lgmn | 0.046355 | 0.790 |
| Cav3 | 0.046381 | 1.659 |
| Tmed4 | 0.046438 | 1.485 |
| Ahsg | 0.04664 | 1.485 |
| Nfkbiz | 0.046718 | 0.660 |
| Mpc2 | 0.046753 | 1.537 |
| Cd52 | 0.047017 | 0.646 |
| Gsg1l | 0.047313 | 1.464 |
| Tmem42 | 0.047629 | 1.580 |
| Wnt5a | 0.047658 | 1.569 |
| Eci1 | 0.04786 | 1.670 |
| Arhgap17 | 0.048108 | 0.737 |
| Prcp | 0.048162 | 0.712 |
| Fam105a | 0.048201 | 0.722 |
| Tor3a | 0.048209 | 0.693 |
| Hbb-bs | 0.048296 | 1.526 |
| Timm17a | 0.048512 | 1.537 |
| Antxr1 | 0.048515 | 1.366 |
| Trim14 | 0.048602 | 0.637 |
| Tpd52 | 0.048609 | 0.753 |
| Epb41 | 0.048748 | 1.376 |
| Ptgr2 | 0.048787 | 1.434 |
| Gm44746 | 0.048905 | 0.624 |
| Rnpep | 0.048942 | 0.722 |

| | | |
|---|---|---|
| Card11 | 0.049001 | 0.693 |
| Mdm1 | 0.049138 | 0.599 |
| Trpm2 | 0.049159 | 0.616 |
| Tnfrsf11a | 0.049211 | 0.717 |
| Ttbk2 | 0.04932 | 1.548 |
| Atp8a2 | 0.049379 | 1.625 |
| Itsn1 | 0.049416 | 0.732 |
| Igf1 | 0.049741 | 0.774 |
| Lrrc25 | 0.049996 | 0.624 |
| Itgb8 | 0.049996 | 1.558 |
| Tnfrsf1b | 0.050092 | 0.763 |
| Cdnf | 0.050111 | 1.613 |
| Rgs10 | 0.050177 | 0.722 |
| Csgalnact2 | 0.050272 | 1.385 |
| Spic | 0.050479 | 0.599 |
| Rnf213 | 0.050609 | 0.785 |
| Epsti1 | 0.050615 | 0.693 |
| Cox6b1 | 0.050744 | 1.495 |
| Akap1 | 0.050881 | 1.625 |
| Sh3bp1 | 0.050932 | 0.637 |

Supplementary Table 2:
Genes within IPA pathways in LXRα S196A versus WT from plaque CD68+ cells

| Mito. Function | |
|---|---|
| **Gene** | **Fold Change** |
| Cox8b | 6.96 |
| Mapk10 | 3.12 |
| Ndufa5 | 2.60 |
| Cox6a2 | 2.53 |
| Aco2 | 2.35 |
| Maob | 2.33 |
| Ndufb9 | 2.28 |
| Snca | 2.24 |
| Pdha1 | 2.14 |
| Atp5o | 2.11 |
| Uqcrfs1 | 2.11 |
| Atp5g1 | 2.10 |
| Cox6c | 2.10 |
| Ndufa4 | 2.04 |
| Cycs | 2.01 |
| Atp5c1 | 1.96 |
| Atp5j | 1.96 |
| Cox7a2 | 1.96 |
| Atp5b | 1.95 |
| Ndufv2 | 1.88 |
| Ndufb5 | 1.85 |
| Atp5j2 | 1.84 |
| Ndufa11 | 1.84 |
| Ndufb11 | 1.82 |
| Atp5a1 | 1.79 |
| Uqcrc2 | 1.78 |
| Uqcr10 | 1.75 |
| Uqcrb | 1.75 |
| Ndufa8 | 1.74 |
| Ndufa12 | 1.74 |
| Ndufs1 | 1.74 |
| Vdac1 | 1.73 |
| Uqcrc1 | 1.72 |
| Mapk12 | 1.71 |
| Ndufa3 | 1.71 |
| Uqcrq | 1.71 |
| Pink1 | 1.68 |
| Ndufb10 | 1.66 |
| Cox4i1 | 1.64 |
| Ndufa13 | 1.64 |
| Ndufs4 | 1.64 |
| Ndufv1 | 1.64 |

| Oxphos | |
|---|---|
| **Gene** | **Fold Change** |
| Cox8b | 6.96 |
| Ndufa5 | 2.60 |
| Cox6a2 | 2.53 |
| Ndufb9 | 2.28 |
| Atp5o | 2.11 |
| Uqcrfs1 | 2.11 |
| Atp5g1 | 2.10 |
| Cox6c | 2.10 |
| Ndufa4 | 2.04 |
| Cycs | 2.01 |
| Atp5c1 | 1.96 |
| Atp5j | 1.96 |
| Cox7a2 | 1.96 |
| Atp5b | 1.95 |
| Ndufv2 | 1.88 |
| Ndufb5 | 1.85 |
| Atp5j2 | 1.84 |
| Ndufa11 | 1.84 |
| Ndufb11 | 1.82 |
| Atp5a1 | 1.79 |
| Uqcrc2 | 1.78 |
| Uqcr10 | 1.75 |
| Uqcrb | 1.75 |
| Ndufa8 | 1.74 |
| Ndufa12 | 1.74 |
| Ndufs1 | 1.74 |
| Uqcrc1 | 1.72 |
| Ndufa3 | 1.71 |
| Uqcrq | 1.71 |
| Ndufb10 | 1.66 |
| Cox4i1 | 1.64 |
| Ndufa13 | 1.64 |
| Ndufs4 | 1.64 |
| Ndufv1 | 1.64 |
| Atp5d | 1.63 |
| Ndufs8 | 1.63 |
| Ndufb2 | 1.61 |
| Ndufb8 | 1.61 |
| Atp5h | 1.60 |
| Ndufs2 | 1.57 |
| Atp5g3 | 1.55 |
| Ndufa6 | 1.55 |

| Sirtuin Signaling | |
|---|---|
| **Gene** | **Fold Change** |
| Ppargc1a | 9.06 |
| Ldhb | 5.06 |
| Pfkm | 4.03 |
| Prkaa2 | 3.53 |
| Ndufa5 | 2.60 |
| Slc25a4 | 2.45 |
| Pgam2 | 2.43 |
| Idh2 | 2.28 |
| Ndufb9 | 2.28 |
| Pdk1 | 2.24 |
| Pdha1 | 2.14 |
| Uqcrfs1 | 2.11 |
| Atp5g1 | 2.10 |
| Ndufa4 | 2.04 |
| Tuba4a | 2.04 |
| Rarb | 2.00 |
| Atp5c1 | 1.96 |
| Atp5j | 1.96 |
| Atp5b | 1.95 |
| Ndufv2 | 1.88 |
| Ndufb5 | 1.85 |
| Ndufa11 | 1.84 |
| Tuba8 | 1.84 |
| Ndufb11 | 1.82 |
| Acadl | 1.80 |
| Atp5a1 | 1.79 |
| Ppara | 1.78 |
| Uqcrc2 | 1.78 |
| Map1lc3a | 1.74 |
| Ndufa8 | 1.74 |
| Ndufa12 | 1.74 |
| Ndufs1 | 1.74 |
| Vdac1 | 1.73 |
| Mapk12 | 1.71 |
| Ndufa3 | 1.71 |
| Ndufb10 | 1.66 |
| Ndufa13 | 1.64 |
| Ndufs4 | 1.64 |
| Ndufv1 | 1.64 |
| Atp5d | 1.63 |
| Ndufs8 | 1.63 |
| Ndufb2 | 1.61 |

| | |
|---|---|
| Atp5d | 1.63 |
| Ndufs8 | 1.63 |
| Ndufb2 | 1.61 |
| Ndufb8 | 1.61 |
| Atp5h | 1.60 |
| Prdx3 | 1.59 |
| Ndufs2 | 1.57 |
| Atp5g3 | 1.55 |
| Ndufa6 | 1.55 |
| Ndufa9 | 1.54 |
| Ndufb7 | 1.54 |
| Atp5f1 | 1.52 |
| Sdhb | 1.52 |

| | |
|---|---|
| Ndufa9 | 1.54 |
| Ndufb7 | 1.54 |
| Atp5f1 | 1.52 |
| Sdhb | 1.52 |

| | |
|---|---|
| Ndufb8 | 1.61 |
| Got2 | 1.59 |
| Ndufs2 | 1.57 |
| Bpgm | 1.56 |
| Ndufa6 | 1.55 |
| Ndufa9 | 1.54 |
| Ndufb7 | 1.54 |
| Timm17a | 1.54 |
| Atp5f1 | 1.52 |
| Sdhb | 1.52 |

| TCA Cycle | |
|---|---|
| **Gene** | **Fold Change** |
| Aco2 | 2.35 |
| Idh3a | 2.14 |
| Fh1 | 1.87 |
| Mdh1 | 1.87 |
| Mdh2 | 1.87 |
| Sucla2 | 1.68 |
| Dhtkd1 | 1.66 |
| Cs | 1.61 |
| Suclg1 | 1.59 |
| Idh3b | 1.53 |
| Sdhb | 1.52 |

| Actin Signaling | |
|---|---|
| **Gene** | **Fold Change** |
| Myh6 | 604.67 |
| Myl4 | 91.14 |
| Actn2 | 54.19 |
| Mylk3 | 36.00 |
| Ttn | 11.24 |
| Fgf12 | 6.96 |
| Myh14 | 4.56 |
| Fgf1 | 3.05 |
| Actc1 | 2.50 |
| Myl2 | 2.45 |
| Pip5k1b | 2.41 |
| Myh7 | 2.27 |
| Gsn | 1.95 |
| Myl9 | 1.89 |
| Fgf13 | 1.73 |
| Tln2 | 1.72 |
| Cfl2 | 1.68 |
| Ppp1r12b | 1.67 |
| Myl7 | 1.65 |
| Fgf2 | 1.51 |

| Fatty Acid Oxidation | |
|---|---|
| **Gene** | **Fold Change** |
| Acsl1 | 2.04 |
| Acadm | 1.96 |
| Acaa2 | 1.87 |
| Eci1 | 1.67 |
| Hadhb | 1.59 |
| Hadh | 1.58 |
| Ivd | 1.55 |

| Cd28 Signaling | |
|---|---|
| **Gene** | **Fold Change** |
| Actr3 | 0.742 |
| Arpc5 | 0.742 |
| Pik3r5 | 0.742 |
| Grb2 | 0.737 |
| Syk | 0.737 |
| Arpc4 | 0.717 |
| Csk | 0.697 |
| Card11 | 0.693 |
| Fcer1g | 0.693 |
| Fyn | 0.664 |
| Pik3cd | 0.651 |
| Ikbke | 0.633 |
| H2-Ab1 | 0.629 |
| Ptprc | 0.624 |
| H2-Eb1 | 0.62 |
| Ptpn6 | 0.611 |
| Vav1 | 0.611 |
| Pik3cg | 0.59 |
| H2-Aa | 0.586 |
| Lcp2 | 0.559 |
| Cd4 | 0.438 |
| Cd86 | 0.435 |

| Phagocytosis in Mac. | |
|---|---|
| **Gene** | **Fold Change** |
| Actr3 | 0.742 |
| Arpc5 | 0.742 |
| Actb | 0.737 |
| Syk | 0.737 |
| Arpc4 | 0.717 |
| Inpp5d | 0.688 |
| Hmox1 | 0.674 |
| Fcgr1 | 0.664 |
| Fyn | 0.664 |
| Fgr | 0.642 |
| Ncf1 | 0.637 |
| Hck | 0.633 |
| Vav1 | 0.611 |
| Lyn | 0.603 |
| Pik3cg | 0.59 |
| Fyb | 0.582 |
| Lcp2 | 0.559 |
| Pld4 | 0.525 |

| Comm. Betw. Imm. C. | |
|---|---|
| **Gene** | **Fold Change** |
| Tlr8 | 0.717 |
| Il1rn | 0.712 |
| Fcer1g | 0.693 |
| Tlr7 | 0.669 |
| H2-Eb1 | 0.62 |
| Tlr9 | 0.595 |
| Cd83 | 0.559 |
| Ccl3 | 0.555 |
| Cxcl10 | 0.532 |
| Ccl9 | 0.518 |
| Cd4 | 0.438 |
| Cd86 | 0.435 |
| Tlr1 | 0.432 |

| NFAT in immune resp. | |
|---|---|
| **Gene** | **Fold Change** |
| Pik3r5 | 0.742 |
| Grb2 | 0.737 |
| Syk | 0.737 |
| Fcer1g | 0.693 |
| Fcgr1 | 0.664 |
| Fyn | 0.664 |
| Blnk | 0.655 |
| Pik3cd | 0.651 |
| Ikbke | 0.633 |
| H2-Ab1 | 0.629 |
| H2-Eb1 | 0.62 |
| Lyn | 0.603 |
| Pik3cg | 0.59 |
| H2-Aa | 0.586 |
| Lcp2 | 0.559 |
| Cd4 | 0.438 |
| Cd86 | 0.435 |

| Th2 signaling | |
|---|---|
| **Gene** | **Fold Change** |
| Pik3r5 | 0.742 |
| Grb2 | 0.737 |
| Spi1 | 0.688 |
| Pik3cd | 0.651 |
| H2-Ab1 | 0.629 |
| Ikzf1 | 0.624 |
| H2-Eb1 | 0.62 |
| Vav1 | 0.611 |
| Pik3cg | 0.59 |
| H2-Aa | 0.586 |
| Maf | 0.547 |
| Cd4 | 0.438 |
| Cd86 | 0.435 |

| NFAT in immune resp. | |
|---|---|
| **Gene** | **Fold Change** |
| Pik3r5 | 0.742 |
| Tlr8 | 0.717 |
| Tnfrsf11a | 0.717 |
| Il1rn | 0.712 |
| Irak4 | 0.697 |
| Tank | 0.697 |
| Card11 | 0.693 |
| Fcer1g | 0.693 |
| Eif2ak2 | 0.683 |
| Tlr7 | 0.669 |
| Pik3cd | 0.651 |
| Tlr9 | 0.595 |
| Pik3cg | 0.59 |
| Tlr1 | 0.432 |

Supplementary Table 3: Genes differentially expressed in ATMs from LXR WT and S196A

| FB WT vs FBC WT | | |
|---|---|---|
| gene | pvalue | FC |
| Fabp5 | 0.01435 | 0.082625 |
| Mmp12 | 5.00E-05 | 0.083651 |
| Mical1 | 0.00995 | 0.087658 |
| Zranb3 | 0.0032 | 0.09101 |
| Itgax | 5.00E-05 | 0.094355 |
| Gpnmb | 5.00E-05 | 0.096862 |
| Dhrs9 | 0.0054 | 0.097883 |
| Il7r | 5.00E-05 | 0.098074 |
| St18 | 0.0386 | 0.115284 |
| Spp1 | 6.00E-04 | 0.120178 |
| Cadm1 | 0.0032 | 0.121517 |
| Cmbl | 0.0358 | 0.122567 |
| Myh15 | 0.00065 | 0.125804 |
| Adk | 0.0152 | 0.136709 |
| Rnf128 | 0.0147 | 0.140501 |
| Efr3b | 0.00505 | 0.142406 |
| Kcnn4 | 0.01095 | 0.145048 |
| Mad2l2 | 0.03675 | 0.149618 |
| Igf2bp2 | 0.0187 | 0.151089 |
| Rras2 | 0.01795 | 0.164021 |
| Ear2 | 4.00E-04 | 0.180829 |
| Bcl2 | 0.016 | 0.18198 |
| Fat3 | 0.00085 | 0.188205 |
| Atp6v0d2 | 5.00E-05 | 0.19039 |
| Ctsk | 0.0192 | 0.191757 |
| Plxdc1 | 0.0368 | 0.192576 |
| Med1 | 0.01605 | 0.192648 |
| Cd9 | 0.001 | 0.199667 |
| Phf8 | 0.03295 | 0.201108 |
| Rrm2 | 0.039 | 0.205881 |
| RP23-285C | 0.01005 | 0.207408 |
| Slc2a1 | 0.04895 | 0.211421 |
| Atp1a3 | 0.0041 | 0.220343 |
| Pcdh7 | 0.00745 | 0.220732 |
| Tnip3 | 5.00E-05 | 0.241339 |
| Dusp5 | 0.0449 | 0.243334 |
| Slc6a8 | 0.00095 | 0.248125 |
| Adam8 | 3.00E-04 | 0.25816 |
| Tnfaip2 | 0.02175 | 0.261101 |
| Ndrg1 | 0.02165 | 0.262498 |
| Nes | 0.00805 | 0.262711 |
| Lpl | 5.00E-05 | 0.263141 |
| Ahnak2 | 0.00505 | 0.267683 |
| Fam96a | 0.02375 | 0.273482 |
| Igf2r | 0.0011 | 0.274582 |
| Card11 | 0.0227 | 0.276787 |
| Tapt1 | 0.043 | 0.278191 |
| Lhfpl2 | 0.00565 | 0.282874 |
| Psat1 | 0.0247 | 0.290321 |
| Tsnax | 0.0294 | 0.293197 |

| FB S196A vs FBC S196A | | |
|---|---|---|
| gene | pvalue | FC |
| Tnip1 | 5.00E-05 | 0.006364 |
| Mrpl54 | 0.01665 | 0.019073 |
| Yipf1 | 5.00E-05 | 0.03966 |
| F7 | 0.03565 | 0.040351 |
| Dph6 | 0.00045 | 0.040824 |
| Mmp12 | 5.00E-05 | 0.05524 |
| Gnpda2 | 0.01485 | 0.05584 |
| Zranb3 | 0.00415 | 0.056565 |
| Lyrm2 | 0.023 | 0.056942 |
| Ykt6 | 0.00935 | 0.064599 |
| Il7r | 5.00E-05 | 0.065137 |
| Igf2bp2 | 0.0031 | 0.068139 |
| Stau2 | 0.00095 | 0.068547 |
| Unc119 | 0.04285 | 0.070228 |
| Gen1 | 0.00425 | 0.071123 |
| Xndc1 | 0.03385 | 0.073328 |
| Pcnxl3 | 0.0013 | 0.076628 |
| Siva1 | 0.01475 | 0.076911 |
| Tmem184b | 0.00385 | 0.081547 |
| Psmd10 | 0.0052 | 0.082331 |
| Gpnmb | 5.00E-05 | 0.084941 |
| Med10 | 0.02035 | 0.087007 |
| Spic | 0.01585 | 0.08922 |
| Fam220a | 0.0465 | 0.091289 |
| Acp5 | 0.04345 | 0.091356 |
| Ptgir | 0.0154 | 0.097516 |
| RP23-116E | 0.0383 | 0.105679 |
| Eif2s3x | 0.0035 | 0.108618 |
| Slc2a9 | 0.00135 | 0.109902 |
| Reep4 | 0.0326 | 0.111216 |
| Cbll1 | 0.0249 | 0.112818 |
| Cuta | 0.0139 | 0.113176 |
| Zfp597 | 0.0423 | 0.11332 |
| Timm17a | 0.0281 | 0.114094 |
| Car5b | 0.0209 | 0.115683 |
| Airn | 0.038 | 0.119035 |
| Kctd2 | 0.04405 | 0.120141 |
| Numb | 3.00E-04 | 0.120521 |
| Ggta1 | 0.00065 | 0.123461 |
| Parp6 | 0.0022 | 0.125185 |
| Psma5 | 0.01025 | 0.1257 |
| Brcc3 | 0.01535 | 0.125903 |
| Pcdh7 | 0.0038 | 0.126905 |
| Hras | 0.0227 | 0.131773 |
| Pdgfa | 0.03305 | 0.132251 |
| Nbeal2 | 0.01765 | 0.133854 |
| Rpf2 | 0.0394 | 0.134368 |
| Gart | 0.01855 | 0.134944 |
| Car13 | 0.02475 | 0.136328 |
| Exosc8 | 0.01225 | 0.138156 |

| Genes unique in | |
|---|---|
| gene | pvalue |
| Mical1 | 0.00995 |
| Dhrs9 | 0.0054 |
| St18 | 0.0386 |
| Spp1 | 6.00E-04 |
| Cmbl | 0.0358 |
| Myh15 | 0.00065 |
| Rnf128 | 0.0147 |
| Efr3b | 0.00505 |
| Mad2l2 | 0.03675 |
| Rras2 | 0.01795 |
| Fat3 | 0.00085 |
| Ctsk | 0.0192 |
| Plxdc1 | 0.0368 |
| Med1 | 0.01605 |
| Phf8 | 0.03295 |
| Rrm2 | 0.039 |
| RP23-285C | 0.01005 |
| Slc2a1 | 0.04895 |
| Tnip3 | 5.00E-05 |
| Tnfaip2 | 0.02175 |
| Ndrg1 | 0.02165 |
| Nes | 0.00805 |
| Fam96a | 0.02375 |
| Card11 | 0.0227 |
| Tapt1 | 0.043 |
| Lhfpl2 | 0.00565 |
| Psat1 | 0.0247 |
| Tsnax | 0.0294 |
| Cdk9 | 0.01925 |
| Cct3 | 0.0493 |
| RP24-234D | 0.0409 |
| Hist1h1c | 0.0344 |
| Tpx2 | 0.0498 |
| Anxa1 | 0.001 |
| Kansl3 | 0.02785 |
| Ftsj3 | 0.04085 |
| AC114920. | 0.0438 |
| Phlpp1 | 0.0405 |
| Tmx4 | 0.0204 |
| Asph | 0.00375 |
| Rangap1 | 0.04885 |
| Kif11 | 0.03825 |
| Myo1e | 0.00965 |
| Sema4d | 0.03995 |
| Tkt | 0.0399 |
| Prkch | 0.0228 |
| Rnh1 | 0.0155 |
| Mgat5 | 0.0449 |
| Tmem65 | 0.02775 |
| Acss1 | 0.04845 |

| | | |
|---|---|---|
| Cdk9 | 0.01925 | 0.29703 |
| Cct3 | 0.0493 | 0.297621 |
| RP24-234D | 0.0409 | 0.307262 |
| Hist1h1c | 0.0344 | 0.307596 |
| Tpx2 | 0.0498 | 0.316373 |
| Anxa1 | 0.001 | 0.317443 |
| Aldh1a2 | 0.01835 | 0.32023 |
| Kansl3 | 0.02785 | 0.330976 |
| Ftsj3 | 0.04085 | 0.33105 |
| AC114920.1 | 0.0438 | 0.336475 |
| Phlpp1 | 0.0405 | 0.336857 |
| Tmx4 | 0.0204 | 0.3382 |
| Asph | 0.00375 | 0.338472 |
| Rangap1 | 0.04885 | 0.34031 |
| Bhlhe40 | 0.00595 | 0.344955 |
| Kif11 | 0.03825 | 0.348022 |
| Lipa | 2.00E-04 | 0.360297 |
| Myo1e | 0.00965 | 0.367068 |
| Lgals3 | 0.0019 | 0.367916 |
| Sema4d | 0.03995 | 0.373736 |
| Anxa2 | 0.00425 | 0.376353 |
| Arhgap25 | 0.01355 | 0.376458 |
| Lyz1 | 0.0127 | 0.393319 |
| Atp6v1b2 | 0.00315 | 0.398494 |
| Tkt | 0.0399 | 0.398687 |
| Prkch | 0.0228 | 0.399348 |
| Cd180 | 0.00595 | 0.404595 |
| Rnh1 | 0.0155 | 0.405215 |
| Slc9a4 | 0.04555 | 0.407957 |
| Mgat5 | 0.0449 | 0.408738 |
| Capg | 0.0219 | 0.415189 |
| Tmem65 | 0.02775 | 0.415488 |
| Acss1 | 0.04845 | 0.415794 |
| Myo10 | 0.0401 | 0.4179 |
| Tomm22 | 0.04635 | 0.421239 |
| Slc37a2 | 0.01385 | 0.422045 |
| Frmd6 | 0.04195 | 0.423605 |
| Dpep2 | 0.02555 | 0.428465 |
| Ell2 | 0.04255 | 0.43632 |
| Spcs3 | 0.02915 | 0.436714 |
| Itga6 | 0.0444 | 0.43844 |
| Ctsz | 0.0183 | 0.442365 |
| Mkl2 | 0.0264 | 0.443523 |
| Efr3a | 0.036 | 0.444686 |
| Rgs1 | 0.0282 | 0.456818 |
| Cd36 | 0.0151 | 0.458407 |
| Ctsl | 0.01865 | 0.460068 |
| Ctss | 0.00275 | 0.462803 |
| Pa2g4 | 0.0351 | 0.469293 |
| Anp32b | 0.0202 | 0.469937 |
| Cd63 | 0.02655 | 0.474257 |
| Fabp4 | 0.04615 | 0.479938 |
| Abcg1 | 0.0153 | 0.486351 |

| | | |
|---|---|---|
| Vkorc1l1 | 0.00955 | 0.141436 |
| Fam78b | 0.01205 | 0.144517 |
| Lrrc61 | 0.0027 | 0.146319 |
| Coro2a | 0.0052 | 0.146631 |
| Cdh5 | 0.00695 | 0.146821 |
| Mrpl4 | 0.00695 | 0.146842 |
| Lrp12 | 0.0077 | 0.146975 |
| Cstb | 1.00E-04 | 0.147244 |
| Psma6 | 0.0113 | 0.14755 |
| Mrps18a | 0.0182 | 0.14875 |
| Ccl22 | 0.02135 | 0.14937 |
| AC157774.( | 0.02235 | 0.152498 |
| Mtmr9 | 0.01465 | 0.152974 |
| Itgax | 5.00E-05 | 0.152995 |
| Stx3 | 0.0052 | 0.153791 |
| Cope | 0.0055 | 0.154166 |
| Ext2 | 0.005 | 0.155466 |
| Pet100 | 0.0111 | 0.155552 |
| Ndufab1 | 0.02735 | 0.155593 |
| Tex30 | 0.03455 | 0.156325 |
| Ptchd1 | 0.029 | 0.15822 |
| Kcnn4 | 0.0241 | 0.160796 |
| Aamdc | 0.04955 | 0.160915 |
| L1cam | 0.0353 | 0.163192 |
| AC157561.9 | 3.00E-04 | 0.163436 |
| Cacfd1 | 0.0018 | 0.163626 |
| Fam213a | 0.01525 | 0.165829 |
| Ahnak2 | 0.00265 | 0.168332 |
| Gpr68 | 0.0231 | 0.168783 |
| Higd2a | 0.03635 | 0.170318 |
| Abcb8 | 0.02535 | 0.171291 |
| Hilpda | 0.0152 | 0.173012 |
| Eya4 | 0.01245 | 0.17351 |
| Apbb2 | 0.00695 | 0.173556 |
| Zfp946 | 0.0473 | 0.173878 |
| Uprt | 0.00625 | 0.175634 |
| Chn2 | 0.01755 | 0.176496 |
| Wdr91 | 0.00075 | 0.178844 |
| Hspb7 | 0.01875 | 0.18005 |
| Slfn10-ps | 0.0086 | 0.18021 |
| Glipr1 | 0.0293 | 0.180829 |
| Rnf146 | 0.03445 | 0.181422 |
| Atp6v0d2 | 5.00E-05 | 0.183168 |
| Sympk | 0.01255 | 0.184785 |
| Utp14a | 0.0057 | 0.185621 |
| Gdf15 | 0.0467 | 0.185824 |
| Sh2d1b1 | 0.03925 | 0.187497 |
| Cadm1 | 0.0476 | 0.188876 |
| Phf11a | 0.02085 | 0.191093 |
| Drosha | 0.00245 | 0.1913 |
| Plp2 | 0.04395 | 0.192007 |
| Phtf2 | 0.00235 | 0.192512 |
| Zfp748 | 0.0266 | 0.193765 |

| | |
|---|---|
| Myo10 | 0.0401 |
| Tomm22 | 0.04635 |
| Slc37a2 | 0.01385 |
| Frmd6 | 0.04195 |
| Ell2 | 0.04255 |
| Spcs3 | 0.02915 |
| Itga6 | 0.0444 |
| Ctsz | 0.0183 |
| Mkl2 | 0.0264 |
| Ctsl | 0.01865 |
| Pa2g4 | 0.0351 |
| Anp32b | 0.0202 |
| Cd63 | 0.02655 |
| Abcg1 | 0.0153 |
| Ldha | 0.03805 |
| Pon2 | 0.02185 |
| Dynlt3 | 0.04785 |
| C1qc | 0.04925 |
| Ifi207 | 0.0274 |
| Nfkbiz | 0.03225 |
| Notch2 | 0.0398 |
| Siglec1 | 0.03945 |
| Sbf2 | 0.03455 |
| Mtmr12 | 0.04575 |
| Ctsc | 0.0423 |
| Zfp652 | 0.0426 |
| Cited2 | 0.04705 |
| Clock | 0.0447 |
| Ifitm2 | 0.0439 |
| Flnb | 0.04755 |
| Sult1a1 | 0.03575 |
| Dip2b | 0.04895 |
| Usf3 | 0.0253 |
| Nr3c1 | 0.02015 |
| Gda | 0.0075 |
| Rnd3 | 0.0272 |
| Ighm | 0.0288 |
| Ophn1 | 0.02895 |
| Ssh2 | 0.0087 |
| Ubc | 0.0475 |
| Ankhd1 | 0.04 |
| Exoc6b | 0.04575 |
| Ttpal | 0.03335 |
| P3h2 | 0.03705 |
| Ogfrl1 | 0.03445 |
| Wdfy3 | 0.02355 |
| Fcgr2b | 0.01105 |
| Sulf2 | 0.02745 |
| Egfr | 0.02075 |
| Fkbp5 | 0.0148 |
| Man1a | 0.0101 |
| Muc6 | 0.02115 |
| Zbtb38 | 0.0174 |

| | | |
|---|---|---|
| Ldha | 0.03805 | 0.488588 |
| Pld3 | 0.0369 | 0.49545 |
| Ctsd | 0.01095 | 0.498965 |
| Pon2 | 0.02185 | 0.51656 |
| Dynlt3 | 0.04785 | 0.527928 |
| Anpep | 0.03885 | 0.543552 |
| Psap | 0.03 | 0.572203 |
| C1qc | 0.04925 | 1.687848 |
| Mrc1 | 0.0326 | 1.724018 |
| Ifi207 | 0.0274 | 1.772002 |
| Nfkbiz | 0.03225 | 1.773534 |
| Notch2 | 0.0398 | 1.843572 |
| Siglec1 | 0.03945 | 1.876398 |
| Sbf2 | 0.03455 | 1.891616 |
| Mtmr12 | 0.04575 | 1.893971 |
| Ctsc | 0.0423 | 1.897868 |
| Zfp652 | 0.0426 | 1.912071 |
| Cited2 | 0.04705 | 1.919737 |
| Rnase4 | 0.04085 | 1.950267 |
| Clock | 0.0447 | 1.954299 |
| Ifitm2 | 0.0439 | 1.980388 |
| Flnb | 0.04755 | 1.985241 |
| Klf4 | 0.0164 | 1.991165 |
| Sult1a1 | 0.03575 | 1.993011 |
| Dip2b | 0.04895 | 1.996266 |
| Usf3 | 0.0253 | 1.999579 |
| Nr3c1 | 0.02015 | 2.026289 |
| Abca9 | 0.0105 | 2.041203 |
| Gda | 0.0075 | 2.042576 |
| Rnd3 | 0.0272 | 2.053778 |
| Ighm | 0.0288 | 2.056527 |
| Gas6 | 0.0115 | 2.091227 |
| Ophn1 | 0.02895 | 2.096539 |
| Tgfbr2 | 0.01815 | 2.098982 |
| Vcam1 | 0.0114 | 2.108387 |
| Ssh2 | 0.0087 | 2.117101 |
| Psd3 | 0.02505 | 2.145167 |
| Ubc | 0.0475 | 2.147428 |
| Ankhd1 | 0.04 | 2.148054 |
| Exoc6b | 0.04575 | 2.150825 |
| Ttpal | 0.03335 | 2.150944 |
| Lrp6 | 0.00465 | 2.157798 |
| Rnf169 | 0.0066 | 2.170068 |
| P3h2 | 0.03705 | 2.170338 |
| Ogfrl1 | 0.03445 | 2.174344 |
| Filip1l | 0.01515 | 2.178719 |
| Wdfy3 | 0.02355 | 2.186055 |
| Fcgr2b | 0.01105 | 2.187329 |
| Sulf2 | 0.02745 | 2.200483 |
| Zfhx3 | 0.00985 | 2.206959 |
| Egfr | 0.02075 | 2.208398 |
| Fkbp5 | 0.0148 | 2.209684 |
| Man1a | 0.0101 | 2.224868 |

| | | |
|---|---|---|
| Zfp64 | 0.0313 | 0.194174 |
| Gyg | 0.0111 | 0.194502 |
| Prmt1 | 0.03245 | 0.195961 |
| Zbtb8os | 0.0485 | 0.197342 |
| Ikbkap | 0.02065 | 0.197453 |
| Adk | 0.03375 | 0.198189 |
| Ndor1 | 0.04945 | 0.19869 |
| Nudc | 0.03735 | 0.198706 |
| Timm9 | 0.0466 | 0.199749 |
| Mpv17 | 0.0277 | 0.200666 |
| Igf2r | 2.00E-04 | 0.202768 |
| Zfp236 | 0.0281 | 0.203153 |
| Swi5 | 0.04375 | 0.20447 |
| Slc39a14 | 0.0305 | 0.20546 |
| Sf3b6 | 0.0152 | 0.207662 |
| Il10 | 0.0029 | 0.208366 |
| Atp1a3 | 0.0029 | 0.208367 |
| Fmn1 | 0.01095 | 0.21076 |
| Ctnnbl1 | 0.04065 | 0.212106 |
| AC158971.1 | 0.04915 | 0.212246 |
| Trem2 | 0.0033 | 0.213054 |
| Dscr3 | 0.0466 | 0.2134 |
| Mrpl32 | 0.0325 | 0.214323 |
| Atic | 0.027 | 0.214371 |
| Fndc7 | 0.04565 | 0.216373 |
| D3Ertd751e | 0.04295 | 0.218564 |
| Ankrd33b | 0.0276 | 0.219185 |
| Tbcb | 0.0117 | 0.2193 |
| Aldh3b1 | 0.0148 | 0.219565 |
| Zdhhc13 | 0.04985 | 0.220013 |
| Blm | 0.04105 | 0.220112 |
| Nufip1 | 0.0287 | 0.221807 |
| Slc25a40 | 0.02375 | 0.221994 |
| Dpagt1 | 0.0402 | 0.222089 |
| Atox1 | 0.0241 | 0.222221 |
| Lrif1 | 0.02645 | 0.225717 |
| Fbxl14 | 0.0173 | 0.225771 |
| Brca1 | 0.0452 | 0.226609 |
| Fabp5 | 0.04035 | 0.227432 |
| Slc6a8 | 0.0023 | 0.228724 |
| Zscan21 | 0.03035 | 0.228808 |
| Dusp10 | 0.028 | 0.229102 |
| Tnfsf9 | 0.01925 | 0.229318 |
| Brca2 | 0.04655 | 0.229792 |
| Arhgap6 | 0.026 | 0.231525 |
| Timm29 | 0.04295 | 0.232031 |
| Lgals3 | 5.00E-05 | 0.232052 |
| Bhlhe41 | 0.02845 | 0.232972 |
| Rab35 | 0.0155 | 0.234934 |
| Pi4kb | 0.01975 | 0.237288 |
| Dcun1d5 | 0.03655 | 0.237638 |
| Pold3 | 0.01075 | 0.238118 |
| Cd200r4 | 0.04355 | 0.238817 |

| | |
|---|---|
| Hcfc1 | 0.04305 |
| Fbxo22 | 0.0399 |
| Rela | 0.04835 |
| Ccl7 | 0.0173 |
| Pacs1 | 0.0443 |
| Adgre4 | 0.0328 |
| Hpgd | 0.02215 |
| Ly6e | 0.00365 |
| Slc28a2 | 0.03445 |
| Oas2 | 0.0323 |
| Lyve1 | 0.00115 |
| Ppp5c | 0.04555 |
| Tlr9 | 0.04505 |
| Mylip | 0.03265 |
| C1qtnf1 | 0.01295 |
| Ogdh | 0.03955 |
| Zfp36l1 | 0.00865 |
| Smurf1 | 0.04385 |
| Kat2b | 0.03275 |
| Zfp710 | 0.00875 |
| Pou2f1 | 0.03535 |
| Odf2 | 0.04025 |
| Tppp | 0.03485 |
| Eaf1 | 0.0499 |
| Sema4a | 0.0411 |
| Zfp84 | 0.01375 |
| Pknox1 | 0.0192 |
| Tns1 | 0.00125 |
| Plekhg5 | 0.04575 |
| Cbr2 | 0.0057 |
| Dnase1l3 | 0.047 |
| Arid3a | 0.03365 |
| Trp53bp2 | 0.04535 |
| Cd38 | 0.0053 |
| Slc16a14 | 0.0419 |
| Sh3pxd2a | 0.0087 |
| Rreb1 | 0.0141 |
| Smad5 | 0.0113 |
| Btbd3 | 0.02715 |
| Esr1 | 0.0483 |
| Zswim8 | 0.00475 |
| Ttll4 | 0.02865 |
| Cog6 | 0.0144 |
| U2af1l4 | 0.0492 |
| Ifi208 | 0.0109 |
| Rara | 0.0024 |
| Cxcl10 | 0.01115 |
| Map7d3 | 0.0364 |
| Susd1 | 0.04585 |
| Zmym4 | 0.00715 |
| RP23-314A | 0.04985 |
| Hk3 | 0.00865 |
| RP23-200C | 0.0459 |

| | | |
|---|---|---|
| Muc6 | 0.02115 | 2.229005 |
| Zbtb38 | 0.0174 | 2.262927 |
| Hcfc1 | 0.04305 | 2.289575 |
| Jun | 0.00205 | 2.289797 |
| Fbxo22 | 0.0399 | 2.292068 |
| Egr1 | 0.0011 | 2.29423 |
| Rela | 0.04835 | 2.310075 |
| Igfbp4 | 0.0243 | 2.316843 |
| Ccl7 | 0.0173 | 2.333815 |
| Pacs1 | 0.0443 | 2.341203 |
| F13a1 | 0.00205 | 2.352199 |
| Adgre4 | 0.0328 | 2.361445 |
| Man2b2 | 0.03995 | 2.385365 |
| Hpgd | 0.02215 | 2.389171 |
| Ly6e | 0.00365 | 2.390513 |
| Thbs1 | 0.0347 | 2.406056 |
| Slc28a2 | 0.03445 | 2.442977 |
| Oas2 | 0.0323 | 2.447587 |
| Lyve1 | 0.00115 | 2.453481 |
| Ppp5c | 0.04555 | 2.453566 |
| Tlr9 | 0.04505 | 2.479433 |
| Mylip | 0.03265 | 2.515195 |
| C1qtnf1 | 0.01295 | 2.515543 |
| Tcof1 | 0.0186 | 2.516764 |
| Sema6d | 0.003 | 2.530461 |
| Ogdh | 0.03955 | 2.552003 |
| Zfp36l1 | 0.00865 | 2.552109 |
| Smurf1 | 0.04385 | 2.564629 |
| RP23-410L1 | 0.0038 | 2.57671 |
| Maf | 5.00E-04 | 2.629922 |
| Kat2b | 0.03275 | 2.655659 |
| Kbtbd11 | 0.0128 | 2.683043 |
| Zfp710 | 0.00875 | 2.700544 |
| Pou2f1 | 0.03535 | 2.724554 |
| Ifi213 | 0.01065 | 2.738071 |
| Lifr | 0.0013 | 2.743714 |
| Bank1 | 0.00895 | 2.78431 |
| Nhsl2 | 0.00205 | 2.799327 |
| Zfp329 | 0.0471 | 2.809727 |
| Odf2 | 0.04025 | 2.823785 |
| Tppp | 0.03485 | 2.872944 |
| Eaf1 | 0.0499 | 2.882159 |
| Sema4a | 0.0411 | 2.893569 |
| Tlr5 | 0.00625 | 2.906615 |
| Zfp84 | 0.01375 | 2.95525 |
| Pknox1 | 0.0192 | 2.968245 |
| Arhgef5 | 0.03055 | 2.978715 |
| Tns1 | 0.00125 | 3.043856 |
| F830016B0 | 0.0197 | 3.054254 |
| Plekhg5 | 0.04575 | 3.067365 |
| Cbr2 | 0.0057 | 3.084571 |
| Dnase1l3 | 0.047 | 3.086453 |
| Adrb2 | 0.0143 | 3.097255 |

| | | |
|---|---|---|
| Cd9 | 0.01795 | 0.23959 |
| Aldh1a2 | 0.00425 | 0.239869 |
| Csnk2a2 | 0.03285 | 0.240354 |
| Med6 | 0.0493 | 0.241163 |
| Jag1 | 0.0435 | 0.241377 |
| Sepw1 | 0.02325 | 0.241539 |
| Slc29a1 | 0.04055 | 0.242233 |
| Dlat | 0.03835 | 0.242395 |
| Ear2 | 0.00045 | 0.245083 |
| Adam8 | 0.0019 | 0.245579 |
| Man1a2 | 0.04135 | 0.246551 |
| Gdf3 | 0.01275 | 0.247399 |
| Evl | 0.00785 | 0.248711 |
| Mad2l1 | 0.03955 | 0.250678 |
| Trappc1 | 0.04825 | 0.251307 |
| Txn1 | 0.00195 | 0.25151 |
| Dpp9 | 0.04665 | 0.25462 |
| Ufc1 | 0.03085 | 0.254765 |
| Aip | 0.04915 | 0.255953 |
| Etaa1 | 0.04585 | 0.258051 |
| Leo1 | 0.0462 | 0.258293 |
| Acat1 | 0.0297 | 0.258397 |
| Tfdp2 | 0.02175 | 0.258402 |
| Chchd3 | 0.014 | 0.259084 |
| Pparg | 0.01365 | 0.260588 |
| Fuca2 | 0.0184 | 0.261065 |
| Trappc3 | 0.0163 | 0.261485 |
| Dot1l | 0.01515 | 0.261944 |
| Sh2b2 | 0.0234 | 0.26226 |
| Dusp5 | 0.03575 | 0.262704 |
| Pus7 | 0.04415 | 0.264117 |
| Ugp2 | 0.004 | 0.264896 |
| Fam20b | 0.03895 | 0.265941 |
| Smim15 | 0.01205 | 0.267988 |
| Mcm3 | 0.03565 | 0.26889 |
| Stard7 | 0.03165 | 0.270069 |
| Wipf2 | 0.043 | 0.270539 |
| Mcm9 | 0.04445 | 0.271112 |
| Synj2 | 0.0352 | 0.271987 |
| Traf1 | 0.0209 | 0.273083 |
| Csnk1g2 | 0.03995 | 0.276803 |
| Paip2b | 0.0499 | 0.278823 |
| Itpr1 | 0.0207 | 0.280799 |
| Ociad1 | 0.01225 | 0.282984 |
| Atp6v0e | 0.01365 | 0.283675 |
| Vipas39 | 0.03495 | 0.283732 |
| Mamld1 | 0.04475 | 0.283942 |
| Nostrin | 0.04275 | 0.284433 |
| Ddx47 | 0.03165 | 0.285357 |
| Mtch2 | 0.03275 | 0.286338 |
| Cul5 | 0.00785 | 0.286509 |
| Vps11 | 0.02385 | 0.28884 |
| Heg1 | 0.0148 | 0.289058 |

| | |
|---|---|
| Herc3 | 0.02285 |
| Adam22 | 0.0399 |
| Pml | 0.0124 |
| Mctp2 | 0.0169 |
| Ago3 | 0.0314 |
| Brms1l | 0.0378 |
| Etv5 | 0.0439 |
| Eral1 | 0.0404 |
| Extl2 | 0.03905 |
| Tnfsf10 | 0.03035 |
| Vps8 | 0.02055 |
| Iqsec2 | 0.02585 |
| Siglece | 0.0327 |
| Med25 | 0.0379 |
| Yeats2 | 0.03945 |
| Prelp | 0.0373 |
| RP23-173H | 0.00905 |
| Aldh6a1 | 0.00695 |
| Nid1 | 0.02495 |
| Ifit3b | 0.047 |
| Kremen1 | 0.0483 |
| Strn4 | 0.01805 |
| Tmem150b | 0.0457 |
| Saa3 | 0.04115 |
| Vcan | 0.0469 |
| Ssbp3 | 0.01435 |
| Mboat1 | 0.0403 |
| Fxyd1 | 0.048 |
| Fgf11 | 0.0174 |
| Ston1 | 0.0237 |
| RP24-309H | 0.0288 |
| Senp8 | 0.02195 |
| Myo1b | 0.0378 |
| Hmgb1 | 2.00E-04 |
| 0610009O2 | 0.01235 |
| Adhfe1 | 0.012 |
| Serpinb2 | 0.0447 |
| Dnah14 | 0.0392 |
| Alox15 | 0.0111 |
| Chil3 | 0.0156 |
| Ptprm | 0.00915 |
| RP23-149L1 | 0.0022 |
| Slc7a6 | 0.0061 |
| Slit3 | 0.0483 |
| Rgs18 | 5.00E-05 |
| Ticrr | 4.00E-04 |
| Nsg1 | 0.03295 |

| | | |
|---|---|---|
| Arid3a | 0.03365 | 3.110917 |
| Trp53bp2 | 0.04535 | 3.111068 |
| Cd38 | 0.0053 | 3.115362 |
| Foxred2 | 0.0108 | 3.137662 |
| Cd163 | 0.00015 | 3.172478 |
| Mgl2 | 0.00185 | 3.193083 |
| Slc16a14 | 0.0419 | 3.197579 |
| Sh3pxd2a | 0.0087 | 3.213198 |
| Rreb1 | 0.0141 | 3.23017 |
| Smad5 | 0.0113 | 3.233777 |
| Btbd3 | 0.02715 | 3.29393 |
| Cd33 | 0.0015 | 3.305687 |
| Esr1 | 0.0483 | 3.342089 |
| Wipf2 | 0.0415 | 3.406476 |
| Cmah | 0.0021 | 3.414442 |
| Zswim8 | 0.00475 | 3.417781 |
| Ttll4 | 0.02865 | 3.470708 |
| Cog6 | 0.0144 | 3.484472 |
| U2af1l4 | 0.0492 | 3.493444 |
| Ifi208 | 0.0109 | 3.569668 |
| Dph6 | 0.03515 | 3.623716 |
| Rara | 0.0024 | 3.62545 |
| Cxcl10 | 0.01115 | 3.637381 |
| Slc14a1 | 0.0111 | 3.641367 |
| Map7d3 | 0.0364 | 3.688234 |
| Susd1 | 0.04585 | 3.703425 |
| Zmym4 | 0.00715 | 3.824073 |
| RP23-314A | 0.04985 | 3.841421 |
| Hk3 | 0.00865 | 3.856174 |
| RP23-200C | 0.0459 | 3.867148 |
| Herc3 | 0.02285 | 3.908188 |
| Adam22 | 0.0399 | 3.985747 |
| Sh3d19 | 0.01395 | 3.98793 |
| Pml | 0.0124 | 4.024027 |
| Mctp2 | 0.0169 | 4.153881 |
| Ago3 | 0.0314 | 4.173044 |
| Brms1l | 0.0378 | 4.176487 |
| Etv5 | 0.0439 | 4.195172 |
| Eral1 | 0.0404 | 4.20006 |
| Extl2 | 0.03905 | 4.261289 |
| Tnfsf10 | 0.03035 | 4.271403 |
| Vps8 | 0.02055 | 4.304364 |
| Iqsec2 | 0.02585 | 4.316404 |
| Siglece | 0.0327 | 4.322302 |
| Med25 | 0.0379 | 4.346277 |
| Mcoln2 | 0.0083 | 4.352306 |
| Slfn1 | 0.0221 | 4.402399 |
| Yeats2 | 0.03945 | 4.578548 |
| Prelp | 0.0373 | 4.609116 |
| RP23-173H | 0.00905 | 4.721646 |
| Aldh6a1 | 0.00695 | 4.971909 |
| Nid1 | 0.02495 | 5.028751 |
| Ifit3b | 0.047 | 5.203347 |

| | | |
|---|---|---|
| Phf11b | 0.0416 | 0.289114 |
| Arpp19 | 0.0393 | 0.289581 |
| Prpf31 | 0.03615 | 0.289653 |
| Sntb2 | 0.0073 | 0.290631 |
| Chst11 | 0.0469 | 0.294476 |
| Tcirg1 | 0.00405 | 0.294612 |
| Ak2 | 0.01155 | 0.295154 |
| Ctss | 5.00E-05 | 0.296841 |
| Pdk1 | 0.0335 | 0.297186 |
| Mfsd12 | 0.01615 | 0.300644 |
| Ago4 | 0.00855 | 0.300724 |
| Leprot | 0.0128 | 0.301435 |
| Cul2 | 0.0326 | 0.30204 |
| Wsb2 | 0.01895 | 0.302631 |
| Fem1b | 0.00235 | 0.302805 |
| Dnajb11 | 0.0129 | 0.304309 |
| Rac2 | 0.01785 | 0.304529 |
| Reps1 | 0.04725 | 0.305114 |
| P4ha1 | 0.04265 | 0.307068 |
| Pik3cb | 0.0122 | 0.307381 |
| Stap1 | 0.0116 | 0.307677 |
| Ticam1 | 0.037 | 0.310305 |
| Actr1b | 0.0213 | 0.312319 |
| Zc3h12c | 0.01535 | 0.313411 |
| Fbxo33 | 0.0294 | 0.313655 |
| Tspan5 | 0.03465 | 0.313694 |
| Sod2 | 0.0291 | 0.315325 |
| Ccdc90b | 0.04695 | 0.315344 |
| Slc16a10 | 0.02055 | 0.316569 |
| Atg12 | 0.0265 | 0.317494 |
| RP23-167C | 0.03915 | 0.320456 |
| Ctsd | 0.00015 | 0.321259 |
| Ndufa4 | 0.0443 | 0.32164 |
| Slc7a1 | 0.0159 | 0.321908 |
| Coa5 | 0.00195 | 0.322216 |
| Fabp4 | 0.00925 | 0.323209 |
| Slc25a4 | 0.0319 | 0.324044 |
| Anxa2 | 0.0019 | 0.325762 |
| Sort1 | 0.0405 | 0.326142 |
| Tars | 0.0252 | 0.327487 |
| Abcd3 | 0.04425 | 0.328722 |
| Capg | 0.00525 | 0.3295 |
| Trap1 | 0.0278 | 0.330474 |
| Sparc | 0.0301 | 0.330685 |
| Anpep | 2.00E-04 | 0.332635 |
| Eif3h | 0.03405 | 0.333514 |
| Dpep2 | 0.0043 | 0.335055 |
| S100a11 | 0.0248 | 0.335129 |
| Lrpap1 | 0.0365 | 0.336832 |
| Plaur | 0.0399 | 0.33707 |
| Txndc9 | 0.03725 | 0.337488 |
| Txndc17 | 0.00865 | 0.337502 |
| Flrt2 | 0.03165 | 0.337748 |

| | | |
|---|---|---|
| Kremen1 | 0.0483 | 5.450959 |
| Strn4 | 0.01805 | 5.461738 |
| Tmem150b | 0.0457 | 5.506059 |
| Saa3 | 0.04115 | 5.588457 |
| Vcan | 0.0469 | 5.661365 |
| Ssbp3 | 0.01435 | 5.892623 |
| Mboat1 | 0.0403 | 6.15219 |
| Fxyd1 | 0.048 | 6.221144 |
| Fgf11 | 0.0174 | 6.292616 |
| Ston1 | 0.0237 | 6.460071 |
| RP24-309H | 0.0288 | 6.600796 |
| Senp8 | 0.02195 | 6.606563 |
| Myo1b | 0.0378 | 6.613849 |
| Hmgb1 | 2.00E-04 | 6.863764 |
| 0610009O2 | 0.01235 | 7.190988 |
| Il20rb | 0.03415 | 7.287985 |
| Adhfe1 | 0.012 | 7.666191 |
| Serpinb2 | 0.0447 | 7.826461 |
| Dnah14 | 0.0392 | 8.160797 |
| Alox15 | 0.0111 | 8.511067 |
| Chil3 | 0.0156 | 8.532509 |
| F5 | 0.00195 | 9.296966 |
| Ptprm | 0.00915 | 9.320582 |
| RP23-149L1 | 0.0022 | 9.549863 |
| Slc7a6 | 0.0061 | 10.94052 |
| Slit3 | 0.0483 | 12.01754 |
| Rgs18 | 5.00E-05 | 12.04765 |
| Ticrr | 4.00E-04 | 13.0607 |
| Rnase2a | 0.03425 | 14.871 |
| Nsg1 | 0.03295 | 16.52067 |

| | | |
|---|---|---|
| Atxn3 | 0.0399 | 0.339071 |
| Arl8b | 0.01585 | 0.33949 |
| Rnf4 | 0.0266 | 0.339789 |
| Rcan1 | 0.0291 | 0.34304 |
| Sike1 | 0.0302 | 0.343962 |
| RP24-482A | 0.0496 | 0.346746 |
| Ddx52 | 0.024 | 0.348111 |
| Cyp4v3 | 0.0167 | 0.348374 |
| Lrrc47 | 0.02985 | 0.351986 |
| Lpl | 0.00015 | 0.353531 |
| Elavl1 | 0.0347 | 0.355239 |
| Nlrp1c-ps | 0.0209 | 0.355362 |
| Tmem126a | 0.04955 | 0.356287 |
| Atp5c1 | 0.01845 | 0.35789 |
| Cd52 | 0.00925 | 0.359004 |
| Ctbs | 0.01945 | 0.35993 |
| Rilpl2 | 0.03965 | 0.36592 |
| Sdhd | 0.02175 | 0.36733 |
| Rap2a | 0.02875 | 0.367746 |
| Prelid3b | 0.02305 | 0.368179 |
| Dync1i2 | 0.0126 | 0.368286 |
| Fermt3 | 0.02105 | 0.368552 |
| Setd3 | 0.0375 | 0.368805 |
| Ube2z | 0.0372 | 0.372905 |
| Ipo8 | 0.03725 | 0.373927 |
| Bmi1 | 0.04285 | 0.375223 |
| Nek6 | 0.0224 | 0.380477 |
| Nt5c2 | 0.0292 | 0.382937 |
| Kpna6 | 0.0316 | 0.383958 |
| Mfhas1 | 0.03405 | 0.385855 |
| Smpdl3a | 0.00605 | 0.385951 |
| Ost4 | 0.03025 | 0.389493 |
| Vdac2 | 0.0191 | 0.396288 |
| Tbl1xr1 | 0.0113 | 0.399443 |
| Gas2l3 | 0.00405 | 0.401096 |
| Cd84 | 0.00505 | 0.401241 |
| Rps17 | 0.021 | 0.404859 |
| Cct4 | 0.0299 | 0.404988 |
| Ide | 0.04285 | 0.405451 |
| Bhlhe40 | 0.008 | 0.406924 |
| Nceh1 | 0.0097 | 0.408014 |
| Pla2g7 | 0.0116 | 0.408571 |
| Rab6a | 0.0341 | 0.408968 |
| Akr1a1 | 0.0068 | 0.40903 |
| Lipa | 9.00E-04 | 0.409365 |
| Psmd8 | 0.03585 | 0.411513 |
| Wdfy1 | 0.0376 | 0.412307 |
| Eif5a | 0.0334 | 0.413704 |
| Ppp2r5c | 0.02295 | 0.41586 |
| Amfr | 0.04615 | 0.425429 |
| Rpsa | 0.0168 | 0.429699 |
| Slc9a4 | 0.03865 | 0.4304 |
| Gnas | 0.01855 | 0.430531 |

| | | |
|---|---|---|
| Fxyd5 | 0.0118 | 0.431332 |
| Rps27a | 0.04455 | 0.43649 |
| Gusb | 0.018 | 0.4382 |
| Ccrl2 | 0.0158 | 0.440917 |
| Cdc27 | 0.03265 | 0.445813 |
| Cct5 | 0.0473 | 0.446893 |
| Tcp1 | 0.0287 | 0.449434 |
| Ifrd1 | 0.034 | 0.452168 |
| Atp6v1b2 | 0.0362 | 0.452469 |
| Rgs1 | 0.04075 | 0.455875 |
| Clta | 0.03765 | 0.461199 |
| Rpl32 | 0.0248 | 0.463159 |
| Il1b | 0.0151 | 0.463654 |
| Pttg1ip | 0.04405 | 0.471883 |
| Gfpt1 | 0.04625 | 0.473478 |
| Cd36 | 0.0176 | 0.47916 |
| Efhd2 | 0.0399 | 0.480664 |
| Vat1 | 0.04665 | 0.488459 |
| Psap | 0.0058 | 0.490281 |
| Cd180 | 0.0149 | 0.4943 |
| M6pr | 0.03035 | 0.495123 |
| Ptbp1 | 0.02835 | 0.496409 |
| Idh1 | 0.0479 | 0.497501 |
| Pkm | 0.049 | 0.501114 |
| Pld3 | 0.0399 | 0.503397 |
| Cxcl1 | 0.01935 | 0.504255 |
| Arhgap5 | 0.04635 | 0.506802 |
| Cat | 0.0474 | 0.521437 |
| Lyz1 | 0.0325 | 0.523874 |
| Pmp22 | 0.0389 | 0.527088 |
| Arhgap25 | 0.0425 | 0.530589 |
| Cd68 | 0.03135 | 0.533575 |
| Itgb2 | 0.03755 | 0.539211 |
| Cd83 | 0.0497 | 0.567596 |
| Ctsb | 0.03255 | 0.579054 |
| Zcchc6 | 0.0487 | 1.769383 |
| Vcam1 | 0.04325 | 1.811656 |
| Slc43a2 | 0.04465 | 1.822309 |
| Gas6 | 0.0361 | 1.82918 |
| Abca9 | 0.02835 | 1.831151 |
| Fosb | 0.02945 | 1.843142 |
| Parp14 | 0.04235 | 1.849355 |
| Cd300ld | 0.034 | 1.851085 |
| Zfhx3 | 0.03215 | 1.877779 |
| Sat1 | 0.046 | 1.90176 |
| Tgfbr2 | 0.03595 | 1.909542 |
| Klf4 | 0.0205 | 1.941305 |
| Sp1 | 0.03655 | 1.941379 |
| Ifi209 | 0.045 | 1.952689 |
| Arf3 | 0.0431 | 1.9572 |
| Frmd4b | 0.0323 | 1.957525 |
| Rbpj | 0.03105 | 1.959306 |
| Il6ra | 0.04515 | 1.981431 |

| | | |
|---|---|---|
| Slc9a9 | 0.0486 | 1.984589 |
| Rnase4 | 0.0463 | 2.007584 |
| Slfn5 | 0.03445 | 2.013227 |
| Gpatch8 | 0.03595 | 2.01352 |
| Samhd1 | 0.01925 | 2.027511 |
| Rb1cc1 | 0.038 | 2.037161 |
| Mrc1 | 0.0054 | 2.044857 |
| Man2a2 | 0.021 | 2.046459 |
| Gfra2 | 0.0393 | 2.053308 |
| Lrp6 | 0.01875 | 2.062265 |
| Zscan26 | 0.03115 | 2.062594 |
| Ostf1 | 0.03055 | 2.065427 |
| Dstn | 0.0331 | 2.069023 |
| Dab2 | 0.0089 | 2.076552 |
| Arfgef2 | 0.04685 | 2.077416 |
| Phf20l1 | 0.0352 | 2.08125 |
| Dcaf7 | 0.0372 | 2.086045 |
| Dclre1c | 0.0262 | 2.097644 |
| Adam19 | 0.0157 | 2.112234 |
| Trim56 | 0.02555 | 2.118334 |
| F13a1 | 0.00425 | 2.120508 |
| Rtf1 | 0.034 | 2.134088 |
| Zswim6 | 0.014 | 2.159983 |
| Ino80 | 0.0363 | 2.168444 |
| Iqgap2 | 0.0086 | 2.171512 |
| Rfx7 | 0.0377 | 2.177496 |
| Psd3 | 0.0193 | 2.179021 |
| Nfkbie | 0.038 | 2.185116 |
| Eps15 | 0.00885 | 2.18742 |
| Cd81 | 0.02715 | 2.18836 |
| Ccpg1 | 0.0382 | 2.188952 |
| Zkscan1 | 0.02495 | 2.189892 |
| Arhgef6 | 0.02685 | 2.195805 |
| Pdxdc1 | 0.0303 | 2.196414 |
| Ccar1 | 0.03185 | 2.197343 |
| Pja2 | 0.0428 | 2.204284 |
| Trafd1 | 0.0273 | 2.206898 |
| Ski | 0.0123 | 2.209776 |
| Smurf2 | 0.044 | 2.211017 |
| Wdr43 | 0.04685 | 2.218724 |
| Ier5 | 0.00585 | 2.220308 |
| Cnot6l | 0.04455 | 2.221109 |
| Egr1 | 0.0019 | 2.226457 |
| Bms1 | 0.0306 | 2.231169 |
| Tcof1 | 0.031 | 2.23284 |
| Hsbp1 | 0.04715 | 2.237473 |
| Dmxl2 | 0.02575 | 2.239785 |
| Cmah | 0.0251 | 2.241042 |
| Mgl2 | 0.01185 | 2.249602 |
| Myo9a | 0.04685 | 2.265705 |
| Akap8 | 0.02875 | 2.267496 |
| Prpf8 | 0.0101 | 2.271161 |
| Msl1 | 0.01685 | 2.273193 |

| | | |
|---|---|---|
| Gna12 | 0.0261 | 2.277498 |
| Igf1r | 0.0379 | 2.2811 |
| Washc2 | 0.01075 | 2.283869 |
| Atf7 | 0.0412 | 2.286182 |
| Plekhm3 | 0.0437 | 2.291512 |
| Smc1a | 0.00915 | 2.29797 |
| Jun | 0.00245 | 2.311501 |
| Usp7 | 0.01775 | 2.312334 |
| Fbxo38 | 0.03925 | 2.315253 |
| Zmynd11 | 0.0276 | 2.327127 |
| Fam208a | 0.0139 | 2.339921 |
| Hlcs | 0.0258 | 2.355886 |
| Trps1 | 0.0095 | 2.35976 |
| Cul1 | 0.0339 | 2.364705 |
| RP23-380L1 | 0.04645 | 2.366492 |
| CH36-200G | 0.0472 | 2.374971 |
| Ms4a6b | 0.00875 | 2.381285 |
| RP23-465M | 0.00825 | 2.387813 |
| Snrnp200 | 0.0364 | 2.388591 |
| Oxr1 | 0.03145 | 2.392037 |
| Dse | 0.00685 | 2.396701 |
| Anp32a | 0.04275 | 2.40144 |
| Cebpd | 0.0492 | 2.409577 |
| Thbs1 | 0.01485 | 2.438426 |
| Gabpb2 | 0.02585 | 2.438443 |
| Arhgap35 | 0.0396 | 2.440725 |
| Klhl28 | 0.0245 | 2.455369 |
| Sun2 | 0.01105 | 2.468133 |
| Arid4b | 0.0437 | 2.476787 |
| Klf7 | 0.0232 | 2.479501 |
| Prkab2 | 0.0279 | 2.48012 |
| Slfn9 | 0.04405 | 2.480877 |
| Hs2st1 | 0.0464 | 2.48344 |
| Ccl12 | 0.0452 | 2.485869 |
| Man2b2 | 0.0394 | 2.486679 |
| Ralgapa1 | 0.00955 | 2.489283 |
| Nsf | 0.01315 | 2.494914 |
| Atp11c | 0.0075 | 2.50585 |
| Maf | 0.00185 | 2.506944 |
| Csnk1g1 | 0.0316 | 2.510266 |
| Zfp871 | 0.00215 | 2.512738 |
| Rufy1 | 0.0354 | 2.515351 |
| RP23-243E1 | 0.00365 | 2.517916 |
| Ncoa2 | 0.01675 | 2.520465 |
| Nabp1 | 0.01385 | 2.520937 |
| Tnrc6c | 0.02235 | 2.527375 |
| Rnf144b | 0.0227 | 2.528216 |
| Dyrk2 | 0.0455 | 2.528777 |
| Ubn1 | 0.0323 | 2.544126 |
| Tsc22d3 | 0.0056 | 2.550358 |
| Brd8 | 0.03665 | 2.550783 |
| St3gal6 | 0.0363 | 2.5565 |
| Supt20 | 0.01515 | 2.557138 |

| | | |
|---|---|---|
| Chd9 | 0.0066 | 2.578514 |
| RP23-242O | 0.0113 | 2.587879 |
| Kdm3b | 0.0059 | 2.588237 |
| Golt1b | 0.0424 | 2.590589 |
| Pign | 0.0175 | 2.596197 |
| Rai14 | 0.04965 | 2.598664 |
| Slc15a2 | 0.04955 | 2.599168 |
| Slfn8 | 0.0084 | 2.631217 |
| Cnot10 | 0.03415 | 2.636365 |
| Vsir | 0.00725 | 2.638741 |
| Rif1 | 0.01765 | 2.645866 |
| RP23-410L1 | 0.0021 | 2.646159 |
| Fam222b | 0.04695 | 2.648141 |
| Rnf169 | 0.002 | 2.650528 |
| RP24-271M | 0.0132 | 2.65266 |
| Cd163 | 0.00045 | 2.654592 |
| Inpp5d | 0.0025 | 2.664546 |
| Nacc2 | 0.0089 | 2.669223 |
| Tanc2 | 0.0309 | 2.678379 |
| Chka | 0.0266 | 2.679679 |
| Nek9 | 0.01095 | 2.68295 |
| Larp7 | 0.04095 | 2.686783 |
| Mex3c | 0.01905 | 2.709056 |
| RP23-14F5. | 0.0114 | 2.720138 |
| Coro7 | 0.0495 | 2.728863 |
| Narf | 0.03935 | 2.729866 |
| Gbf1 | 0.04125 | 2.731872 |
| Atf7ip | 0.00375 | 2.738698 |
| Efr3a | 0.01635 | 2.743048 |
| Filip1l | 0.0025 | 2.745788 |
| Zcchc2 | 0.0029 | 2.758703 |
| Nbeal1 | 0.0106 | 2.763851 |
| Slc25a37 | 0.0025 | 2.791267 |
| Stab1 | 0.02675 | 2.791809 |
| Zmiz2 | 0.0224 | 2.793105 |
| Arid5a | 0.00835 | 2.805796 |
| Rnf141 | 0.00785 | 2.8085 |
| Engase | 0.03545 | 2.815303 |
| Klhl18 | 0.0438 | 2.818466 |
| Alkbh5 | 0.0127 | 2.829192 |
| Clcn4 | 0.01495 | 2.829368 |
| Nbas | 0.013 | 2.843918 |
| Etv1 | 0.0164 | 2.846443 |
| Senp1 | 0.01225 | 2.859712 |
| Zbtb4 | 0.0433 | 2.862846 |
| Cd33 | 0.0033 | 2.864613 |
| Lifr | 6.00E-04 | 2.869064 |
| Rgs5 | 0.04305 | 2.869263 |
| Plekhg1 | 0.0498 | 2.869402 |
| Ldah | 0.02 | 2.879803 |
| Zbtb34 | 0.00265 | 2.884477 |
| Tubgcp3 | 0.01885 | 2.890201 |
| Nop58 | 0.0193 | 2.895334 |

| | | |
|---|---|---|
| Opa1 | 0.0036 | 2.908207 |
| Pan2 | 0.01335 | 2.908328 |
| Tia1 | 0.034 | 2.909477 |
| Tmem8 | 0.03625 | 2.915473 |
| Ankfy1 | 0.0082 | 2.916989 |
| Jade2 | 0.01725 | 2.921663 |
| Xpnpep1 | 0.0332 | 2.92523 |
| Alms1 | 0.0364 | 2.926041 |
| Dnajc10 | 0.0269 | 2.9362 |
| Dgkd | 0.03815 | 2.944658 |
| ZNF654 | 0.02445 | 2.953468 |
| Klhl6 | 0.0354 | 2.976403 |
| Trim30c | 0.03555 | 2.978529 |
| Hook3 | 0.0389 | 2.979417 |
| Phf6 | 0.0233 | 3.018329 |
| Nfe2l3 | 0.03335 | 3.022202 |
| Anapc2 | 0.0486 | 3.040756 |
| Atxn1 | 0.006 | 3.043075 |
| Slc14a1 | 0.04865 | 3.047212 |
| Becn1 | 0.0055 | 3.057389 |
| Mknk1 | 0.01115 | 3.06643 |
| Rbm4b | 0.02335 | 3.070024 |
| Rxra | 0.0167 | 3.0797 |
| Spaca6 | 0.0267 | 3.102003 |
| Hspa1a | 0.0067 | 3.11288 |
| Slc25a36 | 0.0044 | 3.131969 |
| Creb5 | 0.0025 | 3.132013 |
| Strap | 0.013 | 3.145218 |
| Bbx | 0.00815 | 3.163584 |
| Brd9 | 0.0462 | 3.167578 |
| Txnip | 1.00E-04 | 3.168105 |
| RP23-314A | 0.0264 | 3.169774 |
| Hfe | 0.0147 | 3.174149 |
| Prr14 | 0.01665 | 3.175954 |
| Ptgs1 | 0.00725 | 3.177099 |
| Foxred2 | 0.01225 | 3.194345 |
| Trak2 | 0.00645 | 3.195895 |
| Idh2 | 0.0185 | 3.197136 |
| Tlr5 | 0.00615 | 3.204835 |
| Timm44 | 0.0493 | 3.205035 |
| Nfrkb | 0.04825 | 3.222544 |
| RP24-483G | 0.0403 | 3.235413 |
| Nhsl2 | 0.0014 | 3.250001 |
| Lclat1 | 0.0251 | 3.250384 |
| Gopc | 0.0073 | 3.253111 |
| Hexim1 | 0.0021 | 3.268254 |
| Zfp740 | 0.02205 | 3.268526 |
| Bcor | 0.03305 | 3.275239 |
| Sema6d | 1.00E-04 | 3.284469 |
| Relb | 0.04825 | 3.284514 |
| Crebrf | 0.04415 | 3.29457 |
| Med23 | 0.02205 | 3.309424 |
| Rxrb | 0.046 | 3.32838 |

| | | |
|---|---|---|
| Il4ra | 0.0456 | 3.335909 |
| Lpin1 | 0.00425 | 3.34589 |
| Primpol | 0.0257 | 3.352622 |
| Dennd1c | 0.02525 | 3.354831 |
| Slc35b3 | 0.04955 | 3.366245 |
| Ufd1l | 0.0418 | 3.369559 |
| Inip | 0.01065 | 3.372504 |
| Fnta | 0.0179 | 3.381187 |
| Dlg4 | 0.0333 | 3.381656 |
| Npl | 0.0362 | 3.391468 |
| Sipa1l1 | 0.0166 | 3.392291 |
| Fbxw2 | 0.01165 | 3.393726 |
| Ppcdc | 0.02855 | 3.409523 |
| Eef2k | 0.02675 | 3.426225 |
| Pik3r4 | 0.0229 | 3.426391 |
| Slc31a2 | 0.00165 | 3.429528 |
| Erp44 | 0.03585 | 3.436048 |
| Tsc22d4 | 0.02865 | 3.441744 |
| Nolc1 | 0.0245 | 3.453741 |
| Kif23 | 0.0174 | 3.458772 |
| Pigz | 0.02895 | 3.469289 |
| Homer1 | 0.0026 | 3.474872 |
| Dnajc18 | 0.04245 | 3.484085 |
| Sae1 | 0.0086 | 3.499988 |
| Slc20a1 | 0.014 | 3.501638 |
| Ddx24 | 0.00435 | 3.512358 |
| Tmem194b | 0.04185 | 3.521695 |
| Ccnb2 | 0.03945 | 3.524528 |
| Zmym5 | 0.01015 | 3.532624 |
| Arhgef5 | 0.039 | 3.56376 |
| Ifi213 | 0.0041 | 3.582732 |
| Ermard | 0.0144 | 3.585663 |
| Slfn1 | 0.03645 | 3.593077 |
| Ric1 | 0.0104 | 3.594522 |
| Adam33 | 0.0221 | 3.596915 |
| RP23-61D6 | 0.03545 | 3.599608 |
| Leng8 | 0.01555 | 3.600731 |
| Adar | 0.00855 | 3.614059 |
| Katna1 | 0.0117 | 3.629875 |
| Aoah | 2.00E-04 | 3.630051 |
| Pink1 | 0.029 | 3.631486 |
| Fam73a | 0.0461 | 3.633223 |
| Ddx60 | 1.00E-04 | 3.701885 |
| Spon1 | 0.0076 | 3.703733 |
| RP23-23B1 | 0.0043 | 3.721257 |
| Igfbp4 | 0.0108 | 3.721515 |
| Kmt2d | 0.04295 | 3.735496 |
| Slc26a11 | 0.02665 | 3.741742 |
| Gm26767 | 0.02975 | 3.756737 |
| Kat8 | 0.04535 | 3.7825 |
| Ttc5 | 0.0448 | 3.799343 |
| Qsox2 | 0.0124 | 3.818616 |
| Wdr44 | 0.0031 | 3.821185 |

| | | |
|---|---|---|
| Cd99l2 | 0.03945 | 3.827441 |
| Cdk19 | 6.00E-04 | 3.853902 |
| Cd2bp2 | 0.0321 | 3.877106 |
| Thrb | 0.04935 | 3.910302 |
| Fam65b | 0.0133 | 3.910383 |
| Casz1 | 0.03915 | 3.93379 |
| Ankrd50 | 0.0418 | 3.939684 |
| Fam160b2 | 0.00895 | 3.943673 |
| Cdkn2aip | 0.02685 | 3.94939 |
| Ebp | 0.0415 | 3.951663 |
| Cabin1 | 0.024 | 3.973196 |
| Ypel2 | 0.02795 | 3.988566 |
| Ccp110 | 0.0184 | 4.004577 |
| Tab3 | 0.01915 | 4.022744 |
| Calhm2 | 0.0057 | 4.03232 |
| Snrnp48 | 0.0481 | 4.039481 |
| Foxo4 | 0.0428 | 4.04402 |
| As3mt | 0.03875 | 4.047076 |
| Notch1 | 0.0277 | 4.049096 |
| Cxcl12 | 0.00895 | 4.061746 |
| Phf12 | 0.00555 | 4.080625 |
| Phf23 | 0.00175 | 4.101922 |
| Smad4 | 0.018 | 4.111942 |
| Hcfc2 | 0.03 | 4.121959 |
| RP23-331E1 | 0.0426 | 4.124988 |
| Sp2 | 0.03925 | 4.126504 |
| Itga9 | 0.00195 | 4.130024 |
| Zfp715 | 0.0203 | 4.132801 |
| RP23-185B1 | 0.01805 | 4.134922 |
| Lpar1 | 0.00515 | 4.157626 |
| Lcorl | 0.00345 | 4.167609 |
| Pacrgl | 0.04685 | 4.178601 |
| Cped1 | 0.04545 | 4.180397 |
| Dusp6 | 0.00735 | 4.190348 |
| Dbf4 | 0.03805 | 4.19651 |
| Smarcc1 | 0.00025 | 4.237167 |
| Txlna | 0.0173 | 4.240428 |
| Kansl2 | 0.0499 | 4.242104 |
| Mbtps2 | 0.03505 | 4.260167 |
| Fam118a | 0.02255 | 4.260433 |
| Hdac1 | 0.0073 | 4.270811 |
| Atp10d | 0.02005 | 4.271048 |
| Zfp950 | 0.02235 | 4.27235 |
| Snhg5 | 0.0434 | 4.272469 |
| Add1 | 0.00825 | 4.274957 |
| AC126938.1 | 0.01 | 4.285995 |
| Ptpn21 | 0.0246 | 4.303946 |
| Mmp13 | 0.0339 | 4.3247 |
| RP23-211P1 | 0.00285 | 4.328449 |
| B3gnt3 | 0.04315 | 4.329019 |
| Jrkl | 0.03445 | 4.33806 |
| Snx7 | 0.01645 | 4.346548 |
| Bank1 | 0.00045 | 4.371141 |

| | | |
|---|---|---|
| Znrf1 | 0.0136 | 4.376902 |
| Nadk2 | 0.03685 | 4.378267 |
| Rnase6 | 0.0231 | 4.38024 |
| Ccdc171 | 0.02245 | 4.403376 |
| RP24-354K1 | 0.0103 | 4.411808 |
| Prdm10 | 0.0367 | 4.41704 |
| C3 | 0.03975 | 4.458072 |
| Wdr13 | 0.0065 | 4.459463 |
| F830016B0 | 0.002 | 4.482986 |
| Zfp39 | 0.012 | 4.505696 |
| Mfsd6 | 0.00015 | 4.518613 |
| Zfyve9 | 0.01115 | 4.537948 |
| Mthfr | 0.0089 | 4.573663 |
| Fam210a | 0.00035 | 4.573885 |
| Gcc2 | 0.00945 | 4.589637 |
| Gsr | 0.0254 | 4.597183 |
| Zfp398 | 0.02365 | 4.642719 |
| Pou6f1 | 0.0436 | 4.643749 |
| Kank2 | 0.00335 | 4.65219 |
| RP23-212I2 | 0.0363 | 4.656158 |
| Adrb2 | 0.0068 | 4.705735 |
| Armc10 | 0.00485 | 4.816817 |
| Scaf1 | 0.0359 | 4.830056 |
| Snapc4 | 0.03415 | 4.849581 |
| Arc | 0.04715 | 4.851094 |
| Aasdh | 0.03375 | 4.867564 |
| Elac2 | 0.01655 | 4.87256 |
| Zfp180 | 0.011 | 4.898398 |
| Alkbh3 | 0.0119 | 4.910058 |
| Zfp942 | 0.0393 | 4.921882 |
| Tex10 | 0.0345 | 4.949353 |
| Zfp329 | 0.00335 | 4.951618 |
| RP23-354M | 0.01955 | 4.962957 |
| Pfas | 0.03 | 4.964368 |
| RP23-17C1 | 0.017 | 4.973116 |
| Zbtb20 | 3.00E-04 | 4.988652 |
| Ankrd13c | 0.00085 | 4.996992 |
| Kbtbd11 | 7.00E-04 | 5.034156 |
| Mcoln2 | 0.00765 | 5.057765 |
| Ptgr1 | 0.03405 | 5.060185 |
| Fam53c | 0.0108 | 5.135092 |
| Scly | 0.018 | 5.150777 |
| Zfp595 | 0.04995 | 5.152955 |
| Malsu1 | 0.0193 | 5.160247 |
| Mdc1 | 0.0062 | 5.229272 |
| Wdr77 | 0.0302 | 5.229489 |
| Lrrc40 | 0.00845 | 5.232825 |
| Prkaca | 0.0048 | 5.24161 |
| Dhx8 | 0.01445 | 5.243172 |
| Arfgap2 | 0.00705 | 5.30102 |
| Pole | 0.0182 | 5.325659 |
| Gcc1 | 0.001 | 5.335302 |
| Olfr101 | 0.0443 | 5.338262 |

| | | |
|---|---|---|
| Mob3b | 0.0032 | 5.352415 |
| Nubp2 | 0.045 | 5.364672 |
| Car3 | 0.0225 | 5.400826 |
| Ube2w | 0.0109 | 5.431309 |
| Snrk | 0.0065 | 5.449297 |
| Olfr920 | 0.01 | 5.456251 |
| Snx19 | 0.0248 | 5.459467 |
| Bpgm | 0.0314 | 5.47341 |
| Ica1l | 0.0289 | 5.502015 |
| Snupn | 0.03725 | 5.505067 |
| Il20rb | 0.0359 | 5.513047 |
| Abcc3 | 0.0012 | 5.541014 |
| Zfp939 | 0.04505 | 5.551317 |
| Arrdc4 | 0.00025 | 5.57708 |
| Icos | 0.0429 | 5.594736 |
| Sh3bp1 | 0.01635 | 5.639081 |
| Zbtb6 | 0.0092 | 5.6556 |
| Ddx55 | 0.03285 | 5.685117 |
| Wdr81 | 0.00285 | 5.709443 |
| Mrvi1 | 0.00765 | 5.722716 |
| Top3a | 0.0094 | 5.731012 |
| Arl6ip6 | 0.00215 | 5.779762 |
| Suds3 | 0.0167 | 5.781284 |
| Gatsl2 | 0.01235 | 5.861378 |
| Senp7 | 0.002 | 5.866866 |
| Hjurp | 0.00305 | 5.878712 |
| Zfp658 | 0.02795 | 5.884705 |
| Snrnp27 | 0.0292 | 5.902066 |
| Atcayos | 0.02625 | 5.925061 |
| Cep290 | 0.00045 | 5.979563 |
| Rnf135 | 0.0136 | 5.995043 |
| Accs | 0.00035 | 6.008438 |
| RP24-252B | 0.00855 | 6.044864 |
| Hace1 | 0.02985 | 6.128101 |
| Zfp212 | 0.0326 | 6.134178 |
| Banp | 0.0475 | 6.149376 |
| Nup54 | 0.00685 | 6.20792 |
| Sh3d19 | 0.0022 | 6.228393 |
| Edc3 | 0.0047 | 6.230034 |
| Cd209b | 0.00165 | 6.353053 |
| Kif24 | 0.01035 | 6.415492 |
| Serping1 | 0.0164 | 6.496893 |
| Cyb5d2 | 0.01835 | 6.512403 |
| RP24-64D2 | 0.0221 | 6.534514 |
| Helb | 0.01785 | 6.595536 |
| Gipc1 | 0.02425 | 6.633315 |
| Eci1 | 0.0252 | 6.701435 |
| Tmem144 | 0.03125 | 6.812621 |
| Carf | 0.00965 | 6.842199 |
| Kctd3 | 0.0244 | 6.845709 |
| Zfp169 | 0.04485 | 6.858485 |
| Hhat | 0.0311 | 7.046613 |
| Mx2 | 0.0419 | 7.16755 |

| | | |
|---|---|---|
| Ddx18 | 0.0083 | 7.188646 |
| Dcun1d3 | 0.0219 | 7.205957 |
| Slc35a1 | 0.00605 | 7.426642 |
| Atraid | 0.03095 | 7.427671 |
| Zbtb7b | 0.00485 | 7.476277 |
| Rit1 | 0.0061 | 7.557767 |
| Tor4a | 0.00535 | 7.711332 |
| RP23-313J1 | 0.04655 | 7.803439 |
| AC103620.4 | 0.0018 | 7.819845 |
| AC124461.3 | 6.00E-04 | 7.866653 |
| Zfp637 | 0.02985 | 7.978845 |
| Zfp747 | 0.0034 | 8.142264 |
| Syngap1 | 0.02795 | 8.326903 |
| Ralgps2 | 0.01195 | 8.3861 |
| Zdhhc23 | 0.00155 | 8.400936 |
| Aldh7a1 | 0.0134 | 8.401518 |
| Dhdds | 0.00525 | 8.579479 |
| Setd1a | 0.00595 | 8.77458 |
| Med12 | 5.00E-05 | 8.845723 |
| Nipa1 | 0.03075 | 8.936439 |
| Urb1 | 0.0077 | 8.937616 |
| Nmb | 0.04335 | 9.001341 |
| Zfp961 | 0.0019 | 9.086281 |
| Fbxo7 | 0.00345 | 9.105637 |
| RP23-95L9. | 0.03225 | 9.121746 |
| Ada | 0.0071 | 9.157857 |
| RP23-446P1 | 0.01675 | 9.173994 |
| Rasgrp2 | 0.0048 | 9.26493 |
| Dnaaf5 | 0.0146 | 9.433937 |
| Zfp952 | 0.0017 | 9.557147 |
| Bcl2 | 5.00E-05 | 9.686933 |
| Rffl | 0.0033 | 9.776581 |
| Cyp51 | 0.02675 | 9.857419 |
| RP23-293N | 0.00545 | 9.886638 |
| Ttll1 | 0.0312 | 10.26997 |
| Tmem91 | 0.0046 | 10.46152 |
| Trim41 | 0.04645 | 10.50635 |
| Zfp764 | 0.0184 | 10.82361 |
| Aspa | 0.00015 | 11.26668 |
| Mfsd7a | 0.00675 | 11.27332 |
| Ankrd26 | 0.00095 | 11.31575 |
| Zfyve16 | 5.00E-05 | 11.43821 |
| St5 | 0.00315 | 11.67637 |
| Mdm1 | 0.0044 | 11.68892 |
| Map3k9 | 0.00365 | 11.72609 |
| Polh | 0.00755 | 11.72885 |
| F5 | 5.00E-05 | 11.78418 |
| Ralgps1 | 0.00485 | 11.82829 |
| Acads | 0.0206 | 11.87685 |
| Nedd4 | 0.00115 | 12.16614 |
| Chek2 | 6.00E-04 | 12.22913 |
| Exosc3 | 0.008 | 12.43513 |
| Pdk4 | 0.0015 | 12.46966 |

| | | |
|---|---|---|
| Mccc2 | 0.02185 | 13.34225 |
| Surf6 | 0.00225 | 13.34872 |
| Tmem140 | 0.0156 | 14.89659 |
| Plag1 | 0.0041 | 15.51154 |
| Rabepk | 5.00E-05 | 16.05099 |
| Trip10 | 0.0393 | 16.58722 |
| RP23-357J7 | 0.01415 | 16.89343 |
| AC115748.[illegible] | 0.03715 | 16.94115 |
| Sun1 | 5.00E-05 | 17.1245 |
| Arl15 | 0.01425 | 18.07658 |
| Rpgrip1 | 0.0026 | 19.78685 |
| RP23-281H | 0.04665 | 23.2734 |
| Dars2 | 0.003 | 32.96698 |
| Rnase2a | 0.01825 | 34.18567 |
| RP24-180B[illegible] | 0.0417 | 40.11111 |
| Zfp882 | 0.0041 | 40.38339 |
| Prg4 | 3.00E-04 | 45.33364 |

| n WT |
|---|
| FC |
| 0.087658 |
| 0.097883 |
| 0.115284 |
| 0.120178 |
| 0.122567 |
| 0.125804 |
| 0.140501 |
| 0.142406 |
| 0.149618 |
| 0.164021 |
| 0.188205 |
| 0.191757 |
| 0.192576 |
| 0.192648 |
| 0.201108 |
| 0.205881 |
| 0.207408 |
| 0.211421 |
| 0.241339 |
| 0.261101 |
| 0.262498 |
| 0.262711 |
| 0.273482 |
| 0.276787 |
| 0.278191 |
| 0.282874 |
| 0.290321 |
| 0.293197 |
| 0.29703 |
| 0.297621 |
| 0.307262 |
| 0.307596 |
| 0.316373 |
| 0.317443 |
| 0.330976 |
| 0.33105 |
| 0.336475 |
| 0.336857 |
| 0.3382 |
| 0.338472 |
| 0.34031 |
| 0.348022 |
| 0.367068 |
| 0.373736 |
| 0.398687 |
| 0.399348 |
| 0.405215 |
| 0.408738 |
| 0.415488 |
| 0.415794 |

| Genes unique in S196A | | |
|---|---|---|
| gene | pvalue | FC |
| Tnip1 | 5.00E-05 | 0.006364 |
| Mrpl54 | 0.01665 | 0.019073 |
| Yipf1 | 5.00E-05 | 0.03966 |
| F7 | 0.03565 | 0.040351 |
| Gnpda2 | 0.01485 | 0.05584 |
| Lyrm2 | 0.023 | 0.056942 |
| Ykt6 | 0.00935 | 0.064599 |
| Stau2 | 0.00095 | 0.068547 |
| Unc119 | 0.04285 | 0.070228 |
| Gen1 | 0.00425 | 0.071123 |
| Xndc1 | 0.03385 | 0.073328 |
| Pcnxl3 | 0.0013 | 0.076628 |
| Siva1 | 0.01475 | 0.076911 |
| Tmem184b | 0.00385 | 0.081547 |
| Psmd10 | 0.0052 | 0.082331 |
| Med10 | 0.02035 | 0.087007 |
| Spic | 0.01585 | 0.08922 |
| Fam220a | 0.0465 | 0.091289 |
| Acp5 | 0.04345 | 0.091356 |
| Ptgir | 0.0154 | 0.097516 |
| RP23-116E[illegible] | 0.0383 | 0.105679 |
| Eif2s3x | 0.0035 | 0.108618 |
| Slc2a9 | 0.00135 | 0.109902 |
| Reep4 | 0.0326 | 0.111216 |
| Cbll1 | 0.0249 | 0.112818 |
| Cuta | 0.0139 | 0.113176 |
| Zfp597 | 0.0423 | 0.11332 |
| Timm17a | 0.0281 | 0.114094 |
| Car5b | 0.0209 | 0.115683 |
| Airn | 0.038 | 0.119035 |
| Kctd2 | 0.04405 | 0.120141 |
| Numb | 3.00E-04 | 0.120521 |
| Ggta1 | 0.00065 | 0.123461 |
| Parp6 | 0.0022 | 0.125185 |
| Psma5 | 0.01025 | 0.1257 |
| Brcc3 | 0.01535 | 0.125903 |
| Hras | 0.0227 | 0.131773 |
| Pdgfa | 0.03305 | 0.132251 |
| Nbeal2 | 0.01765 | 0.133854 |
| Rpf2 | 0.0394 | 0.134368 |
| Gart | 0.01855 | 0.134944 |
| Car13 | 0.02475 | 0.136328 |
| Exosc8 | 0.01225 | 0.138156 |
| Vkorc1l1 | 0.00955 | 0.141436 |
| Fam78b | 0.01205 | 0.144517 |
| Lrrc61 | 0.0027 | 0.146319 |
| Coro2a | 0.0052 | 0.146631 |
| Cdh5 | 0.00695 | 0.146821 |
| Mrpl4 | 0.00695 | 0.146842 |
| Lrp12 | 0.0077 | 0.146975 |

| Commun in WT and SA | | | | |
|---|---|---|---|---|
| | FB WT vs FBC WT | | FB WT vs FBC SA | |
| gene | pvalue | FC | pvalue | FC |
| Fabp5 | 0.01435 | 0.082625 | 0.04035 | 0.227432 |
| Mmp12 | 5.00E-05 | 0.083651 | 5.00E-05 | 0.05524 |
| Zranb3 | 0.0032 | 0.09101 | 0.00415 | 0.056565 |
| Itgax | 5.00E-05 | 0.094355 | 5.00E-05 | 0.152995 |
| Gpnmb | 5.00E-05 | 0.096862 | 5.00E-05 | 0.084941 |
| Il7r | 5.00E-05 | 0.098074 | 5.00E-05 | 0.065137 |
| Cadm1 | 0.0032 | 0.121517 | 0.0476 | 0.188876 |
| Adk | 0.0152 | 0.136709 | 0.03375 | 0.198189 |
| Kcnn4 | 0.01095 | 0.145048 | 0.0241 | 0.160796 |
| Igf2bp2 | 0.0187 | 0.151089 | 0.0031 | 0.068139 |
| Ear2 | 4.00E-04 | 0.180829 | 0.00045 | 0.245083 |
| Bcl2 | 0.016 | 0.18198 | 5.00E-05 | 9.686933 |
| Atp6v0d2 | 5.00E-05 | 0.19039 | 5.00E-05 | 0.183168 |
| Cd9 | 0.001 | 0.199667 | 0.01795 | 0.23959 |
| Atp1a3 | 0.0041 | 0.220343 | 0.0029 | 0.208367 |
| Pcdh7 | 0.00745 | 0.220732 | 0.0038 | 0.126905 |
| Dusp5 | 0.0449 | 0.243334 | 0.03575 | 0.262704 |
| Slc6a8 | 0.00095 | 0.248125 | 0.0023 | 0.228724 |
| Adam8 | 3.00E-04 | 0.25816 | 0.0019 | 0.245579 |
| Lpl | 5.00E-05 | 0.263141 | 0.00015 | 0.353531 |
| Ahnak2 | 0.00505 | 0.267683 | 0.00265 | 0.168332 |
| Igf2r | 0.0011 | 0.274582 | 2.00E-04 | 0.202768 |
| Aldh1a2 | 0.01835 | 0.32023 | 0.00425 | 0.239869 |
| Bhlhe40 | 0.00595 | 0.344955 | 0.008 | 0.406924 |
| Lipa | 2.00E-04 | 0.360297 | 9.00E-04 | 0.409365 |
| Lgals3 | 0.0019 | 0.367916 | 5.00E-05 | 0.232052 |
| Anxa2 | 0.00425 | 0.376353 | 0.0019 | 0.325762 |
| Arhgap25 | 0.01355 | 0.376458 | 0.0425 | 0.530589 |
| Lyz1 | 0.0127 | 0.393319 | 0.0325 | 0.523874 |
| Atp6v1b2 | 0.00315 | 0.398494 | 0.0362 | 0.452469 |
| Cd180 | 0.00595 | 0.404595 | 0.0149 | 0.4943 |
| Slc9a4 | 0.04555 | 0.407957 | 0.03865 | 0.4304 |
| Capg | 0.0219 | 0.415189 | 0.00525 | 0.3295 |
| Dpep2 | 0.02555 | 0.428465 | 0.0043 | 0.335055 |
| Efr3a | 0.036 | 0.444686 | 0.01635 | 2.743048 |
| Rgs1 | 0.0282 | 0.456818 | 0.04075 | 0.455875 |
| Cd36 | 0.0151 | 0.458407 | 0.0176 | 0.47916 |
| Ctss | 0.00275 | 0.462803 | 5.00E-05 | 0.296841 |
| Fabp4 | 0.04615 | 0.479938 | 0.00925 | 0.323209 |
| Pld3 | 0.0369 | 0.49545 | 0.0399 | 0.503397 |
| Ctsd | 0.01095 | 0.498965 | 0.00015 | 0.321259 |
| Anpep | 0.03885 | 0.543552 | 2.00E-04 | 0.332635 |
| Psap | 0.03 | 0.572203 | 0.0058 | 0.490281 |
| Mrc1 | 0.0326 | 1.724018 | 0.0054 | 2.044857 |
| Rnase4 | 0.04085 | 1.950267 | 0.0463 | 2.007584 |
| Klf4 | 0.0164 | 1.991165 | 0.0205 | 1.941305 |
| Abca9 | 0.0105 | 2.041203 | 0.02835 | 1.831151 |
| Gas6 | 0.0115 | 2.091227 | 0.0361 | 1.82918 |
| Tgfbr2 | 0.01815 | 2.098982 | 0.03595 | 1.909542 |
| Vcam1 | 0.0114 | 2.108387 | 0.04325 | 1.811656 |

| |
|---|
| 0.4179 |
| 0.421239 |
| 0.422045 |
| 0.423605 |
| 0.43632 |
| 0.436714 |
| 0.43844 |
| 0.442365 |
| 0.443523 |
| 0.460068 |
| 0.469293 |
| 0.469937 |
| 0.474257 |
| 0.486351 |
| 0.488588 |
| 0.51656 |
| 0.527928 |
| 1.687848 |
| 1.772002 |
| 1.773534 |
| 1.843572 |
| 1.876398 |
| 1.891616 |
| 1.893971 |
| 1.897868 |
| 1.912071 |
| 1.919737 |
| 1.954299 |
| 1.980388 |
| 1.985241 |
| 1.993011 |
| 1.996266 |
| 1.999579 |
| 2.026289 |
| 2.042576 |
| 2.053778 |
| 2.056527 |
| 2.096539 |
| 2.117101 |
| 2.147428 |
| 2.148054 |
| 2.150825 |
| 2.150944 |
| 2.170338 |
| 2.174344 |
| 2.186055 |
| 2.187329 |
| 2.200483 |
| 2.208398 |
| 2.209684 |
| 2.224868 |
| 2.229005 |
| 2.262927 |

| | | |
|---|---|---|
| Cstb | 1.00E-04 | 0.147244 |
| Psma6 | 0.0113 | 0.14755 |
| Mrps18a | 0.0182 | 0.14875 |
| Ccl22 | 0.02135 | 0.14937 |
| AC157774.( | 0.02235 | 0.152498 |
| Mtmr9 | 0.01465 | 0.152974 |
| Stx3 | 0.0052 | 0.153791 |
| Cope | 0.0055 | 0.154166 |
| Ext2 | 0.005 | 0.155466 |
| Pet100 | 0.0111 | 0.155552 |
| Ndufab1 | 0.02735 | 0.155593 |
| Tex30 | 0.03455 | 0.156325 |
| Ptchd1 | 0.029 | 0.15822 |
| Aamdc | 0.04955 | 0.160915 |
| L1cam | 0.0353 | 0.163192 |
| AC157561.9 | 3.00E-04 | 0.163436 |
| Cacfd1 | 0.0018 | 0.163626 |
| Fam213a | 0.01525 | 0.165829 |
| Gpr68 | 0.0231 | 0.168783 |
| Higd2a | 0.03635 | 0.170318 |
| Abcb8 | 0.02535 | 0.171291 |
| Hilpda | 0.0152 | 0.173012 |
| Eya4 | 0.01245 | 0.17351 |
| Apbb2 | 0.00695 | 0.173556 |
| Zfp946 | 0.0473 | 0.173878 |
| Uprt | 0.00625 | 0.175634 |
| Chn2 | 0.01755 | 0.176496 |
| Wdr91 | 0.00075 | 0.178844 |
| Hspb7 | 0.01875 | 0.18005 |
| Slfn10-ps | 0.0086 | 0.18021 |
| Glipr1 | 0.0293 | 0.180829 |
| Rnf146 | 0.03445 | 0.181422 |
| Sympk | 0.01255 | 0.184785 |
| Utp14a | 0.0057 | 0.185621 |
| Gdf15 | 0.0467 | 0.185824 |
| Sh2d1b1 | 0.03925 | 0.187497 |
| Phf11a | 0.02085 | 0.191093 |
| Drosha | 0.00245 | 0.1913 |
| Plp2 | 0.04395 | 0.192007 |
| Phtf2 | 0.00235 | 0.192512 |
| Zfp748 | 0.0266 | 0.193765 |
| Zfp64 | 0.0313 | 0.194174 |
| Gyg | 0.0111 | 0.194502 |
| Prmt1 | 0.03245 | 0.195961 |
| Zbtb8os | 0.0485 | 0.197342 |
| Ikbkap | 0.02065 | 0.197453 |
| Ndor1 | 0.04945 | 0.19869 |
| Nudc | 0.03735 | 0.198706 |
| Timm9 | 0.0466 | 0.199749 |
| Mpv17 | 0.0277 | 0.200666 |
| Zfp236 | 0.0281 | 0.203153 |
| Swi5 | 0.04375 | 0.20447 |
| Slc39a14 | 0.0305 | 0.20546 |

| | | | | |
|---|---|---|---|---|
| Psd3 | 0.02505 | 2.145167 | 0.0193 | 2.179021 |
| Lrp6 | 0.00465 | 2.157798 | 0.01875 | 2.062265 |
| Rnf169 | 0.0066 | 2.170068 | 0.002 | 2.650528 |
| Filip1l | 0.01515 | 2.178719 | 0.0025 | 2.745788 |
| Zfhx3 | 0.00985 | 2.206959 | 0.03215 | 1.877779 |
| Jun | 0.00205 | 2.289797 | 0.00245 | 2.311501 |
| Egr1 | 0.0011 | 2.29423 | 0.0019 | 2.226457 |
| Igfbp4 | 0.0243 | 2.316843 | 0.0108 | 3.721515 |
| F13a1 | 0.00205 | 2.352199 | 0.00425 | 2.120508 |
| Man2b2 | 0.03995 | 2.385365 | 0.0394 | 2.486679 |
| Thbs1 | 0.0347 | 2.406056 | 0.01485 | 2.438426 |
| Tcof1 | 0.0186 | 2.516764 | 0.031 | 2.23284 |
| Sema6d | 0.003 | 2.530461 | 1.00E-04 | 3.284469 |
| RP23-410L1 | 0.0038 | 2.57671 | 0.0021 | 2.646159 |
| Maf | 5.00E-04 | 2.629922 | 0.00185 | 2.506944 |
| Kbtbd11 | 0.0128 | 2.683043 | 7.00E-04 | 5.034156 |
| Ifi213 | 0.01065 | 2.738071 | 0.0041 | 3.582732 |
| Lifr | 0.0013 | 2.743714 | 6.00E-04 | 2.869064 |
| Bank1 | 0.00895 | 2.78431 | 0.00045 | 4.371141 |
| Nhsl2 | 0.00205 | 2.799327 | 0.0014 | 3.250001 |
| Zfp329 | 0.0471 | 2.809727 | 0.00335 | 4.951618 |
| Tlr5 | 0.00625 | 2.906615 | 0.00615 | 3.204835 |
| Arhgef5 | 0.03055 | 2.978715 | 0.039 | 3.56376 |
| F830016B0 | 0.0197 | 3.054254 | 0.002 | 4.482986 |
| Adrb2 | 0.0143 | 3.097255 | 0.0068 | 4.705735 |
| Foxred2 | 0.0108 | 3.137662 | 0.01225 | 3.194345 |
| Cd163 | 0.00015 | 3.172478 | 0.00045 | 2.654592 |
| Mgl2 | 0.00185 | 3.193083 | 0.01185 | 2.249602 |
| Cd33 | 0.0015 | 3.305687 | 0.0033 | 2.864613 |
| Wipf2 | 0.0415 | 3.406476 | 0.043 | 0.270539 |
| Cmah | 0.0021 | 3.414442 | 0.0251 | 2.241042 |
| Dph6 | 0.03515 | 3.623716 | 0.00045 | 0.040824 |
| Slc14a1 | 0.0111 | 3.641367 | 0.04865 | 3.047212 |
| Sh3d19 | 0.01395 | 3.98793 | 0.0022 | 6.228393 |
| Mcoln2 | 0.0083 | 4.352306 | 0.00765 | 5.057765 |
| Slfn1 | 0.0221 | 4.402399 | 0.03645 | 3.593077 |
| Il20rb | 0.03415 | 7.287985 | 0.0359 | 5.513047 |
| F5 | 0.00195 | 9.296966 | 5.00E-05 | 11.78418 |
| Rnase2a | 0.03425 | 14.871 | 0.01825 | 34.18567 |

| |
|---|
| 2.289575 |
| 2.292068 |
| 2.310075 |
| 2.333815 |
| 2.341203 |
| 2.361445 |
| 2.389171 |
| 2.390513 |
| 2.442977 |
| 2.447587 |
| 2.453481 |
| 2.453566 |
| 2.479433 |
| 2.515195 |
| 2.515543 |
| 2.552003 |
| 2.552109 |
| 2.564629 |
| 2.655659 |
| 2.700544 |
| 2.724554 |
| 2.823785 |
| 2.872944 |
| 2.882159 |
| 2.893569 |
| 2.95525 |
| 2.968245 |
| 3.043856 |
| 3.067365 |
| 3.084571 |
| 3.086453 |
| 3.110917 |
| 3.111068 |
| 3.115362 |
| 3.197579 |
| 3.213198 |
| 3.23017 |
| 3.233777 |
| 3.29393 |
| 3.342089 |
| 3.417781 |
| 3.470708 |
| 3.484472 |
| 3.493444 |
| 3.569668 |
| 3.62545 |
| 3.637381 |
| 3.688234 |
| 3.703425 |
| 3.824073 |
| 3.841421 |
| 3.856174 |
| 3.867148 |

| | | |
|---|---|---|
| Sf3b6 | 0.0152 | 0.207662 |
| Il10 | 0.0029 | 0.208366 |
| Fmn1 | 0.01095 | 0.21076 |
| Ctnnbl1 | 0.04065 | 0.212106 |
| AC158971.1 | 0.04915 | 0.212246 |
| Trem2 | 0.0033 | 0.213054 |
| Dscr3 | 0.0466 | 0.2134 |
| Mrpl32 | 0.0325 | 0.214323 |
| Atic | 0.027 | 0.214371 |
| Fndc7 | 0.04565 | 0.216373 |
| D3Ertd751e | 0.04295 | 0.218564 |
| Ankrd33b | 0.0276 | 0.219185 |
| Tbcb | 0.0117 | 0.2193 |
| Aldh3b1 | 0.0148 | 0.219565 |
| Zdhhc13 | 0.04985 | 0.220013 |
| Blm | 0.04105 | 0.220112 |
| Nufip1 | 0.0287 | 0.221807 |
| Slc25a40 | 0.02375 | 0.221994 |
| Dpagt1 | 0.0402 | 0.222089 |
| Atox1 | 0.0241 | 0.222221 |
| Lrif1 | 0.02645 | 0.225717 |
| Fbxl14 | 0.0173 | 0.225771 |
| Brca1 | 0.0452 | 0.226609 |
| Zscan21 | 0.03035 | 0.228808 |
| Dusp10 | 0.028 | 0.229102 |
| Tnfsf9 | 0.01925 | 0.229318 |
| Brca2 | 0.04655 | 0.229792 |
| Arhgap6 | 0.026 | 0.231525 |
| Timm29 | 0.04295 | 0.232031 |
| Bhlhe41 | 0.02845 | 0.232972 |
| Rab35 | 0.0155 | 0.234934 |
| Pi4kb | 0.01975 | 0.237288 |
| Dcun1d5 | 0.03655 | 0.237638 |
| Pold3 | 0.01075 | 0.238118 |
| Cd200r4 | 0.04355 | 0.238817 |
| Csnk2a2 | 0.03285 | 0.240354 |
| Med6 | 0.0493 | 0.241163 |
| Jag1 | 0.0435 | 0.241377 |
| Sepw1 | 0.02325 | 0.241539 |
| Slc29a1 | 0.04055 | 0.242233 |
| Dlat | 0.03835 | 0.242395 |
| Man1a2 | 0.04135 | 0.246551 |
| Gdf3 | 0.01275 | 0.247399 |
| Evl | 0.00785 | 0.248711 |
| Mad2l1 | 0.03955 | 0.250678 |
| Trappc1 | 0.04825 | 0.251307 |
| Txn1 | 0.00195 | 0.25151 |
| Dpp9 | 0.04665 | 0.25462 |
| Ufc1 | 0.03085 | 0.254765 |
| Aip | 0.04915 | 0.255953 |
| Etaa1 | 0.04585 | 0.258051 |
| Leo1 | 0.0462 | 0.258293 |
| Acat1 | 0.0297 | 0.258397 |

| |
|---|
| 3.908188 |
| 3.985747 |
| 4.024027 |
| 4.153881 |
| 4.173044 |
| 4.176487 |
| 4.195172 |
| 4.20006 |
| 4.261289 |
| 4.271403 |
| 4.304364 |
| 4.316404 |
| 4.322302 |
| 4.346277 |
| 4.578548 |
| 4.609116 |
| 4.721646 |
| 4.971909 |
| 5.028751 |
| 5.203347 |
| 5.450959 |
| 5.461738 |
| 5.506059 |
| 5.588457 |
| 5.661365 |
| 5.892623 |
| 6.15219 |
| 6.221144 |
| 6.292616 |
| 6.460071 |
| 6.600796 |
| 6.606563 |
| 6.613849 |
| 6.863764 |
| 7.190988 |
| 7.666191 |
| 7.826461 |
| 8.160797 |
| 8.511067 |
| 8.532509 |
| 9.320582 |
| 9.549863 |
| 10.94052 |
| 12.01754 |
| 12.04765 |
| 13.0607 |
| 16.52067 |

| | | |
|---|---|---|
| Tfdp2 | 0.02175 | 0.258402 |
| Chchd3 | 0.014 | 0.259084 |
| Pparg | 0.01365 | 0.260588 |
| Fuca2 | 0.0184 | 0.261065 |
| Trappc3 | 0.0163 | 0.261485 |
| Dot1l | 0.01515 | 0.261944 |
| Sh2b2 | 0.0234 | 0.26226 |
| Pus7 | 0.04415 | 0.264117 |
| Ugp2 | 0.004 | 0.264896 |
| Fam20b | 0.03895 | 0.265941 |
| Smim15 | 0.01205 | 0.267988 |
| Mcm3 | 0.03565 | 0.26889 |
| Stard7 | 0.03165 | 0.270069 |
| Mcm9 | 0.04445 | 0.271112 |
| Synj2 | 0.0352 | 0.271987 |
| Traf1 | 0.0209 | 0.273083 |
| Csnk1g2 | 0.03995 | 0.276803 |
| Paip2b | 0.0499 | 0.278823 |
| Itpr1 | 0.0207 | 0.280799 |
| Ociad1 | 0.01225 | 0.282984 |
| Atp6v0e | 0.01365 | 0.283675 |
| Vipas39 | 0.03495 | 0.283732 |
| Mamld1 | 0.04475 | 0.283942 |
| Nostrin | 0.04275 | 0.284433 |
| Ddx47 | 0.03165 | 0.285357 |
| Mtch2 | 0.03275 | 0.286338 |
| Cul5 | 0.00785 | 0.286509 |
| Vps11 | 0.02385 | 0.28884 |
| Heg1 | 0.0148 | 0.289058 |
| Phf11b | 0.0416 | 0.289114 |
| Arpp19 | 0.0393 | 0.289581 |
| Prpf31 | 0.03615 | 0.289653 |
| Sntb2 | 0.0073 | 0.290631 |
| Chst11 | 0.0469 | 0.294476 |
| Tcirg1 | 0.00405 | 0.294612 |
| Ak2 | 0.01155 | 0.295154 |
| Pdk1 | 0.0335 | 0.297186 |
| Mfsd12 | 0.01615 | 0.300644 |
| Ago4 | 0.00855 | 0.300724 |
| Leprot | 0.0128 | 0.301435 |
| Cul2 | 0.0326 | 0.30204 |
| Wsb2 | 0.01895 | 0.302631 |
| Fem1b | 0.00235 | 0.302805 |
| Dnajb11 | 0.0129 | 0.304309 |
| Rac2 | 0.01785 | 0.304529 |
| Reps1 | 0.04725 | 0.305114 |
| P4ha1 | 0.04265 | 0.307068 |
| Pik3cb | 0.0122 | 0.307381 |
| Stap1 | 0.0116 | 0.307677 |
| Ticam1 | 0.037 | 0.310305 |
| Actr1b | 0.0213 | 0.312319 |
| Zc3h12c | 0.01535 | 0.313411 |
| Fbxo33 | 0.0294 | 0.313655 |

| | | |
|---|---|---|
| Tspan5 | 0.03465 | 0.313694 |
| Sod2 | 0.0291 | 0.315325 |
| Ccdc90b | 0.04695 | 0.315344 |
| Slc16a10 | 0.02055 | 0.316569 |
| Atg12 | 0.0265 | 0.317494 |
| RP23-167C! | 0.03915 | 0.320456 |
| Ndufa4 | 0.0443 | 0.32164 |
| Slc7a1 | 0.0159 | 0.321908 |
| Coa5 | 0.00195 | 0.322216 |
| Slc25a4 | 0.0319 | 0.324044 |
| Sort1 | 0.0405 | 0.326142 |
| Tars | 0.0252 | 0.327487 |
| Abcd3 | 0.04425 | 0.328722 |
| Trap1 | 0.0278 | 0.330474 |
| Sparc | 0.0301 | 0.330685 |
| Eif3h | 0.03405 | 0.333514 |
| S100a11 | 0.0248 | 0.335129 |
| Lrpap1 | 0.0365 | 0.336832 |
| Plaur | 0.0399 | 0.33707 |
| Txndc9 | 0.03725 | 0.337488 |
| Txndc17 | 0.00865 | 0.337502 |
| Flrt2 | 0.03165 | 0.337748 |
| Atxn3 | 0.0399 | 0.339071 |
| Arl8b | 0.01585 | 0.33949 |
| Rnf4 | 0.0266 | 0.339789 |
| Rcan1 | 0.0291 | 0.34304 |
| Sike1 | 0.0302 | 0.343962 |
| RP24-482A | 0.0496 | 0.346746 |
| Ddx52 | 0.024 | 0.348111 |
| Cyp4v3 | 0.0167 | 0.348374 |
| Lrrc47 | 0.02985 | 0.351986 |
| Elavl1 | 0.0347 | 0.355239 |
| Nlrp1c-ps | 0.0209 | 0.355362 |
| Tmem126a | 0.04955 | 0.356287 |
| Atp5c1 | 0.01845 | 0.35789 |
| Cd52 | 0.00925 | 0.359004 |
| Ctbs | 0.01945 | 0.35993 |
| Rilpl2 | 0.03965 | 0.36592 |
| Sdhd | 0.02175 | 0.36733 |
| Rap2a | 0.02875 | 0.367746 |
| Prelid3b | 0.02305 | 0.368179 |
| Dync1i2 | 0.0126 | 0.368286 |
| Fermt3 | 0.02105 | 0.368552 |
| Setd3 | 0.0375 | 0.368805 |
| Ube2z | 0.0372 | 0.372905 |
| Ipo8 | 0.03725 | 0.373927 |
| Bmi1 | 0.04285 | 0.375223 |
| Nek6 | 0.0224 | 0.380477 |
| Nt5c2 | 0.0292 | 0.382937 |
| Kpna6 | 0.0316 | 0.383958 |
| Mfhas1 | 0.03405 | 0.385855 |
| Smpdl3a | 0.00605 | 0.385951 |
| Ost4 | 0.03025 | 0.389493 |

| | | |
|---|---|---|
| Vdac2 | 0.0191 | 0.396288 |
| Tbl1xr1 | 0.0113 | 0.399443 |
| Gas2l3 | 0.00405 | 0.401096 |
| Cd84 | 0.00505 | 0.401241 |
| Rps17 | 0.021 | 0.404859 |
| Cct4 | 0.0299 | 0.404988 |
| Ide | 0.04285 | 0.405451 |
| Nceh1 | 0.0097 | 0.408014 |
| Pla2g7 | 0.0116 | 0.408571 |
| Rab6a | 0.0341 | 0.408968 |
| Akr1a1 | 0.0068 | 0.40903 |
| Psmd8 | 0.03585 | 0.411513 |
| Wdfy1 | 0.0376 | 0.412307 |
| Eif5a | 0.0334 | 0.413704 |
| Ppp2r5c | 0.02295 | 0.41586 |
| Amfr | 0.04615 | 0.425429 |
| Rpsa | 0.0168 | 0.429699 |
| Gnas | 0.01855 | 0.430531 |
| Fxyd5 | 0.0118 | 0.431332 |
| Rps27a | 0.04455 | 0.43649 |
| Gusb | 0.018 | 0.4382 |
| Ccrl2 | 0.0158 | 0.440917 |
| Cdc27 | 0.03265 | 0.445813 |
| Cct5 | 0.0473 | 0.446893 |
| Tcp1 | 0.0287 | 0.449434 |
| Ifrd1 | 0.034 | 0.452168 |
| Clta | 0.03765 | 0.461199 |
| Rpl32 | 0.0248 | 0.463159 |
| Il1b | 0.0151 | 0.463654 |
| Pttg1ip | 0.04405 | 0.471883 |
| Gfpt1 | 0.04625 | 0.473478 |
| Efhd2 | 0.0399 | 0.480664 |
| Vat1 | 0.04665 | 0.488459 |
| M6pr | 0.03035 | 0.495123 |
| Ptbp1 | 0.02835 | 0.496409 |
| Idh1 | 0.0479 | 0.497501 |
| Pkm | 0.049 | 0.501114 |
| Cxcl1 | 0.01935 | 0.504255 |
| Arhgap5 | 0.04635 | 0.506802 |
| Cat | 0.0474 | 0.521437 |
| Pmp22 | 0.0389 | 0.527088 |
| Cd68 | 0.03135 | 0.533575 |
| Itgb2 | 0.03755 | 0.539211 |
| Cd83 | 0.0497 | 0.567596 |
| Ctsb | 0.03255 | 0.579054 |
| Zcchc6 | 0.0487 | 1.769383 |
| Slc43a2 | 0.04465 | 1.822309 |
| Fosb | 0.02945 | 1.843142 |
| Parp14 | 0.04235 | 1.849355 |
| Cd300ld | 0.034 | 1.851085 |
| Sat1 | 0.046 | 1.90176 |
| Sp1 | 0.03655 | 1.941379 |
| Ifi209 | 0.045 | 1.952689 |

| | | |
|---|---|---|
| Arf3 | 0.0431 | 1.9572 |
| Frmd4b | 0.0323 | 1.957525 |
| Rbpj | 0.03105 | 1.959306 |
| Il6ra | 0.04515 | 1.981431 |
| Slc9a9 | 0.0486 | 1.984589 |
| Slfn5 | 0.03445 | 2.013227 |
| Gpatch8 | 0.03595 | 2.01352 |
| Samhd1 | 0.01925 | 2.027511 |
| Rb1cc1 | 0.038 | 2.037161 |
| Man2a2 | 0.021 | 2.046459 |
| Gfra2 | 0.0393 | 2.053308 |
| Zscan26 | 0.03115 | 2.062594 |
| Ostf1 | 0.03055 | 2.065427 |
| Dstn | 0.0331 | 2.069023 |
| Dab2 | 0.0089 | 2.076552 |
| Arfgef2 | 0.04685 | 2.077416 |
| Phf20l1 | 0.0352 | 2.08125 |
| Dcaf7 | 0.0372 | 2.086045 |
| Dclre1c | 0.0262 | 2.097644 |
| Adam19 | 0.0157 | 2.112234 |
| Trim56 | 0.02555 | 2.118334 |
| Rtf1 | 0.034 | 2.134088 |
| Zswim6 | 0.014 | 2.159983 |
| Ino80 | 0.0363 | 2.168444 |
| Iqgap2 | 0.0086 | 2.171512 |
| Rfx7 | 0.0377 | 2.177496 |
| Nfkbie | 0.038 | 2.185116 |
| Eps15 | 0.00885 | 2.18742 |
| Cd81 | 0.02715 | 2.18836 |
| Ccpg1 | 0.0382 | 2.188952 |
| Zkscan1 | 0.02495 | 2.189892 |
| Arhgef6 | 0.02685 | 2.195805 |
| Pdxdc1 | 0.0303 | 2.196414 |
| Ccar1 | 0.03185 | 2.197343 |
| Pja2 | 0.0428 | 2.204284 |
| Trafd1 | 0.0273 | 2.206898 |
| Ski | 0.0123 | 2.209776 |
| Smurf2 | 0.044 | 2.211017 |
| Wdr43 | 0.04685 | 2.218724 |
| Ier5 | 0.00585 | 2.220308 |
| Cnot6l | 0.04455 | 2.221109 |
| Bms1 | 0.0306 | 2.231169 |
| Hsbp1 | 0.04715 | 2.237473 |
| Dmxl2 | 0.02575 | 2.239785 |
| Myo9a | 0.04685 | 2.265705 |
| Akap8 | 0.02875 | 2.267496 |
| Prpf8 | 0.0101 | 2.271161 |
| Msl1 | 0.01685 | 2.273193 |
| Gna12 | 0.0261 | 2.277498 |
| Igf1r | 0.0379 | 2.2811 |
| Washc2 | 0.01075 | 2.283869 |
| Atf7 | 0.0412 | 2.286182 |
| Plekhm3 | 0.0437 | 2.291512 |

| | | |
|---|---|---|
| Smc1a | 0.00915 | 2.29797 |
| Usp7 | 0.01775 | 2.312334 |
| Fbxo38 | 0.03925 | 2.315253 |
| Zmynd11 | 0.0276 | 2.327127 |
| Fam208a | 0.0139 | 2.339921 |
| Hlcs | 0.0258 | 2.355886 |
| Trps1 | 0.0095 | 2.35976 |
| Cul1 | 0.0339 | 2.364705 |
| RP23-380L1 | 0.04645 | 2.366492 |
| CH36-200G | 0.0472 | 2.374971 |
| Ms4a6b | 0.00875 | 2.381285 |
| RP23-465M | 0.00825 | 2.387813 |
| Snrnp200 | 0.0364 | 2.388591 |
| Oxr1 | 0.03145 | 2.392037 |
| Dse | 0.00685 | 2.396701 |
| Anp32a | 0.04275 | 2.40144 |
| Cebpd | 0.0492 | 2.409577 |
| Gabpb2 | 0.02585 | 2.438443 |
| Arhgap35 | 0.0396 | 2.440725 |
| Klhl28 | 0.0245 | 2.455369 |
| Sun2 | 0.01105 | 2.468133 |
| Arid4b | 0.0437 | 2.476787 |
| Klf7 | 0.0232 | 2.479501 |
| Prkab2 | 0.0279 | 2.48012 |
| Slfn9 | 0.04405 | 2.480877 |
| Hs2st1 | 0.0464 | 2.48344 |
| Ccl12 | 0.0452 | 2.485869 |
| Ralgapa1 | 0.00955 | 2.489283 |
| Nsf | 0.01315 | 2.494914 |
| Atp11c | 0.0075 | 2.50585 |
| Csnk1g1 | 0.0316 | 2.510266 |
| Zfp871 | 0.00215 | 2.512738 |
| Rufy1 | 0.0354 | 2.515351 |
| RP23-243E1 | 0.00365 | 2.517916 |
| Ncoa2 | 0.01675 | 2.520465 |
| Nabp1 | 0.01385 | 2.520937 |
| Tnrc6c | 0.02235 | 2.527375 |
| Rnf144b | 0.0227 | 2.528216 |
| Dyrk2 | 0.0455 | 2.528777 |
| Ubn1 | 0.0323 | 2.544126 |
| Tsc22d3 | 0.0056 | 2.550358 |
| Brd8 | 0.03665 | 2.550783 |
| St3gal6 | 0.0363 | 2.5565 |
| Supt20 | 0.01515 | 2.557138 |
| Chd9 | 0.0066 | 2.578514 |
| RP23-242O | 0.0113 | 2.587879 |
| Kdm3b | 0.0059 | 2.588237 |
| Golt1b | 0.0424 | 2.590589 |
| Pign | 0.0175 | 2.596197 |
| Rai14 | 0.04965 | 2.598664 |
| Slc15a2 | 0.04955 | 2.599168 |
| Slfn8 | 0.0084 | 2.631217 |
| Cnot10 | 0.03415 | 2.636365 |

| | | |
|---|---|---|
| Vsir | 0.00725 | 2.638741 |
| Rif1 | 0.01765 | 2.645866 |
| Fam222b | 0.04695 | 2.648141 |
| RP24-271M | 0.0132 | 2.65266 |
| Inpp5d | 0.0025 | 2.664546 |
| Nacc2 | 0.0089 | 2.669223 |
| Tanc2 | 0.0309 | 2.678379 |
| Chka | 0.0266 | 2.679679 |
| Nek9 | 0.01095 | 2.68295 |
| Larp7 | 0.04095 | 2.686783 |
| Mex3c | 0.01905 | 2.709056 |
| RP23-14F5. | 0.0114 | 2.720138 |
| Coro7 | 0.0495 | 2.728863 |
| Narf | 0.03935 | 2.729866 |
| Gbf1 | 0.04125 | 2.731872 |
| Atf7ip | 0.00375 | 2.738698 |
| Zcchc2 | 0.0029 | 2.758703 |
| Nbeal1 | 0.0106 | 2.763851 |
| Slc25a37 | 0.0025 | 2.791267 |
| Stab1 | 0.02675 | 2.791809 |
| Zmiz2 | 0.0224 | 2.793105 |
| Arid5a | 0.00835 | 2.805796 |
| Rnf141 | 0.00785 | 2.8085 |
| Engase | 0.03545 | 2.815303 |
| Klhl18 | 0.0438 | 2.818466 |
| Alkbh5 | 0.0127 | 2.829192 |
| Clcn4 | 0.01495 | 2.829368 |
| Nbas | 0.013 | 2.843918 |
| Etv1 | 0.0164 | 2.846443 |
| Senp1 | 0.01225 | 2.859712 |
| Zbtb4 | 0.0433 | 2.862846 |
| Rgs5 | 0.04305 | 2.869263 |
| Plekhg1 | 0.0498 | 2.869402 |
| Ldah | 0.02 | 2.879803 |
| Zbtb34 | 0.00265 | 2.884477 |
| Tubgcp3 | 0.01885 | 2.890201 |
| Nop58 | 0.0193 | 2.895334 |
| Opa1 | 0.0036 | 2.908207 |
| Pan2 | 0.01335 | 2.908328 |
| Tia1 | 0.034 | 2.909477 |
| Tmem8 | 0.03625 | 2.915473 |
| Ankfy1 | 0.0082 | 2.916989 |
| Jade2 | 0.01725 | 2.921663 |
| Xpnpep1 | 0.0332 | 2.92523 |
| Alms1 | 0.0364 | 2.926041 |
| Dnajc10 | 0.0269 | 2.9362 |
| Dgkd | 0.03815 | 2.944658 |
| ZNF654 | 0.02445 | 2.953468 |
| Klhl6 | 0.0354 | 2.976403 |
| Trim30c | 0.03555 | 2.978529 |
| Hook3 | 0.0389 | 2.979417 |
| Phf6 | 0.0233 | 3.018329 |
| Nfe2l3 | 0.03335 | 3.022202 |

| | | |
|---|---|---|
| Anapc2 | 0.0486 | 3.040756 |
| Atxn1 | 0.006 | 3.043075 |
| Becn1 | 0.0055 | 3.057389 |
| Mknk1 | 0.01115 | 3.06643 |
| Rbm4b | 0.02335 | 3.070024 |
| Rxra | 0.0167 | 3.0797 |
| Spaca6 | 0.0267 | 3.102003 |
| Hspa1a | 0.0067 | 3.11288 |
| Slc25a36 | 0.0044 | 3.131969 |
| Creb5 | 0.0025 | 3.132013 |
| Strap | 0.013 | 3.145218 |
| Bbx | 0.00815 | 3.163584 |
| Brd9 | 0.0462 | 3.167578 |
| Txnip | 1.00E-04 | 3.168105 |
| RP23-314A | 0.0264 | 3.169774 |
| Hfe | 0.0147 | 3.174149 |
| Prr14 | 0.01665 | 3.175954 |
| Ptgs1 | 0.00725 | 3.177099 |
| Trak2 | 0.00645 | 3.195895 |
| Idh2 | 0.0185 | 3.197136 |
| Timm44 | 0.0493 | 3.205035 |
| Nfrkb | 0.04825 | 3.222544 |
| RP24-483G | 0.0403 | 3.235413 |
| Lclat1 | 0.0251 | 3.250384 |
| Gopc | 0.0073 | 3.253111 |
| Hexim1 | 0.0021 | 3.268254 |
| Zfp740 | 0.02205 | 3.268526 |
| Bcor | 0.03305 | 3.275239 |
| Relb | 0.04825 | 3.284514 |
| Crebrf | 0.04415 | 3.29457 |
| Med23 | 0.02205 | 3.309424 |
| Rxrb | 0.046 | 3.32838 |
| Il4ra | 0.0456 | 3.335909 |
| Lpin1 | 0.00425 | 3.34589 |
| Primpol | 0.0257 | 3.352622 |
| Dennd1c | 0.02525 | 3.354831 |
| Slc35b3 | 0.04955 | 3.366245 |
| Ufd1l | 0.0418 | 3.369559 |
| Inip | 0.01065 | 3.372504 |
| Fnta | 0.0179 | 3.381187 |
| Dlg4 | 0.0333 | 3.381656 |
| Npl | 0.0362 | 3.391468 |
| Sipa1l1 | 0.0166 | 3.392291 |
| Fbxw2 | 0.01165 | 3.393726 |
| Ppcdc | 0.02855 | 3.409523 |
| Eef2k | 0.02675 | 3.426225 |
| Pik3r4 | 0.0229 | 3.426391 |
| Slc31a2 | 0.00165 | 3.429528 |
| Erp44 | 0.03585 | 3.436048 |
| Tsc22d4 | 0.02865 | 3.441744 |
| Nolc1 | 0.0245 | 3.453741 |
| Kif23 | 0.0174 | 3.458772 |
| Pigz | 0.02895 | 3.469289 |

| | | |
|---|---|---|
| Homer1 | 0.0026 | 3.474872 |
| Dnajc18 | 0.04245 | 3.484085 |
| Sae1 | 0.0086 | 3.499988 |
| Slc20a1 | 0.014 | 3.501638 |
| Ddx24 | 0.00435 | 3.512358 |
| Tmem194b | 0.04185 | 3.521695 |
| Ccnb2 | 0.03945 | 3.524528 |
| Zmym5 | 0.01015 | 3.532624 |
| Ermard | 0.0144 | 3.585663 |
| Ric1 | 0.0104 | 3.594522 |
| Adam33 | 0.0221 | 3.596915 |
| RP23-61D6 | 0.03545 | 3.599608 |
| Leng8 | 0.01555 | 3.600731 |
| Adar | 0.00855 | 3.614059 |
| Katna1 | 0.0117 | 3.629875 |
| Aoah | 2.00E-04 | 3.630051 |
| Pink1 | 0.029 | 3.631486 |
| Fam73a | 0.0461 | 3.633223 |
| Ddx60 | 1.00E-04 | 3.701885 |
| Spon1 | 0.0076 | 3.703733 |
| RP23-23B1 | 0.0043 | 3.721257 |
| Kmt2d | 0.04295 | 3.735496 |
| Slc26a11 | 0.02665 | 3.741742 |
| Gm26767 | 0.02975 | 3.756737 |
| Kat8 | 0.04535 | 3.7825 |
| Ttc5 | 0.0448 | 3.799343 |
| Qsox2 | 0.0124 | 3.818616 |
| Wdr44 | 0.0031 | 3.821185 |
| Cd99l2 | 0.03945 | 3.827441 |
| Cdk19 | 6.00E-04 | 3.853902 |
| Cd2bp2 | 0.0321 | 3.877106 |
| Thrb | 0.04935 | 3.910302 |
| Fam65b | 0.0133 | 3.910383 |
| Casz1 | 0.03915 | 3.93379 |
| Ankrd50 | 0.0418 | 3.939684 |
| Fam160b2 | 0.00895 | 3.943673 |
| Cdkn2aip | 0.02685 | 3.94939 |
| Ebp | 0.0415 | 3.951663 |
| Cabin1 | 0.024 | 3.973196 |
| Ypel2 | 0.02795 | 3.988566 |
| Ccp110 | 0.0184 | 4.004577 |
| Tab3 | 0.01915 | 4.022744 |
| Calhm2 | 0.0057 | 4.03232 |
| Snrnp48 | 0.0481 | 4.039481 |
| Foxo4 | 0.0428 | 4.04402 |
| As3mt | 0.03875 | 4.047076 |
| Notch1 | 0.0277 | 4.049096 |
| Cxcl12 | 0.00895 | 4.061746 |
| Phf12 | 0.00555 | 4.080625 |
| Phf23 | 0.00175 | 4.101922 |
| Smad4 | 0.018 | 4.111942 |
| Hcfc2 | 0.03 | 4.121959 |
| RP23-331E | 0.0426 | 4.124988 |

| | | |
|---|---|---|
| Sp2 | 0.03925 | 4.126504 |
| Itga9 | 0.00195 | 4.130024 |
| Zfp715 | 0.0203 | 4.132801 |
| RP23-185B | 0.01805 | 4.134922 |
| Lpar1 | 0.00515 | 4.157626 |
| Lcorl | 0.00345 | 4.167609 |
| Pacrgl | 0.04685 | 4.178601 |
| Cped1 | 0.04545 | 4.180397 |
| Dusp6 | 0.00735 | 4.190348 |
| Dbf4 | 0.03805 | 4.19651 |
| Smarcc1 | 0.00025 | 4.237167 |
| Txlna | 0.0173 | 4.240428 |
| Kansl2 | 0.0499 | 4.242104 |
| Mbtps2 | 0.03505 | 4.260167 |
| Fam118a | 0.02255 | 4.260433 |
| Hdac1 | 0.0073 | 4.270811 |
| Atp10d | 0.02005 | 4.271048 |
| Zfp950 | 0.02235 | 4.27235 |
| Snhg5 | 0.0434 | 4.272469 |
| Add1 | 0.00825 | 4.274957 |
| AC126938. | 0.01 | 4.285995 |
| Ptpn21 | 0.0246 | 4.303946 |
| Mmp13 | 0.0339 | 4.3247 |
| RP23-211P | 0.00285 | 4.328449 |
| B3gnt3 | 0.04315 | 4.329019 |
| Jrkl | 0.03445 | 4.33806 |
| Snx7 | 0.01645 | 4.346548 |
| Znrf1 | 0.0136 | 4.376902 |
| Nadk2 | 0.03685 | 4.378267 |
| Rnase6 | 0.0231 | 4.38024 |
| Ccdc171 | 0.02245 | 4.403376 |
| RP24-354K | 0.0103 | 4.411808 |
| Prdm10 | 0.0367 | 4.41704 |
| C3 | 0.03975 | 4.458072 |
| Wdr13 | 0.0065 | 4.459463 |
| Zfp39 | 0.012 | 4.505696 |
| Mfsd6 | 0.00015 | 4.518613 |
| Zfyve9 | 0.01115 | 4.537948 |
| Mthfr | 0.0089 | 4.573663 |
| Fam210a | 0.00035 | 4.573885 |
| Gcc2 | 0.00945 | 4.589637 |
| Gsr | 0.0254 | 4.597183 |
| Zfp398 | 0.02365 | 4.642719 |
| Pou6f1 | 0.0436 | 4.643749 |
| Kank2 | 0.00335 | 4.65219 |
| RP23-212I2 | 0.0363 | 4.656158 |
| Armc10 | 0.00485 | 4.816817 |
| Scaf1 | 0.0359 | 4.830056 |
| Snapc4 | 0.03415 | 4.849581 |
| Arc | 0.04715 | 4.851094 |
| Aasdh | 0.03375 | 4.867564 |
| Elac2 | 0.01655 | 4.87256 |
| Zfp180 | 0.011 | 4.898398 |

| | | |
|---|---|---|
| Alkbh3 | 0.0119 | 4.910058 |
| Zfp942 | 0.0393 | 4.921882 |
| Tex10 | 0.0345 | 4.949353 |
| RP23-354M | 0.01955 | 4.962957 |
| Pfas | 0.03 | 4.964368 |
| RP23-17C1 | 0.017 | 4.973116 |
| Zbtb20 | 3.00E-04 | 4.988652 |
| Ankrd13c | 0.00085 | 4.996992 |
| Ptgr1 | 0.03405 | 5.060185 |
| Fam53c | 0.0108 | 5.135092 |
| Scly | 0.018 | 5.150777 |
| Zfp595 | 0.04995 | 5.152955 |
| Malsu1 | 0.0193 | 5.160247 |
| Mdc1 | 0.0062 | 5.229272 |
| Wdr77 | 0.0302 | 5.229489 |
| Lrrc40 | 0.00845 | 5.232825 |
| Prkaca | 0.0048 | 5.24161 |
| Dhx8 | 0.01445 | 5.243172 |
| Arfgap2 | 0.00705 | 5.30102 |
| Pole | 0.0182 | 5.325659 |
| Gcc1 | 0.001 | 5.335302 |
| Olfr101 | 0.0443 | 5.338262 |
| Mob3b | 0.0032 | 5.352415 |
| Nubp2 | 0.045 | 5.364672 |
| Car3 | 0.0225 | 5.400826 |
| Ube2w | 0.0109 | 5.431309 |
| Snrk | 0.0065 | 5.449297 |
| Olfr920 | 0.01 | 5.456251 |
| Snx19 | 0.0248 | 5.459467 |
| Bpgm | 0.0314 | 5.47341 |
| Ica1l | 0.0289 | 5.502015 |
| Snupn | 0.03725 | 5.505067 |
| Abcc3 | 0.0012 | 5.541014 |
| Zfp939 | 0.04505 | 5.551317 |
| Arrdc4 | 0.00025 | 5.57708 |
| Icos | 0.0429 | 5.594736 |
| Sh3bp1 | 0.01635 | 5.639081 |
| Zbtb6 | 0.0092 | 5.6556 |
| Ddx55 | 0.03285 | 5.685117 |
| Wdr81 | 0.00285 | 5.709443 |
| Mrvi1 | 0.00765 | 5.722716 |
| Top3a | 0.0094 | 5.731012 |
| Arl6ip6 | 0.00215 | 5.779762 |
| Suds3 | 0.0167 | 5.781284 |
| Gatsl2 | 0.01235 | 5.861378 |
| Senp7 | 0.002 | 5.866866 |
| Hjurp | 0.00305 | 5.878712 |
| Zfp658 | 0.02795 | 5.884705 |
| Snrnp27 | 0.0292 | 5.902066 |
| Atcayos | 0.02625 | 5.925061 |
| Cep290 | 0.00045 | 5.979563 |
| Rnf135 | 0.0136 | 5.995043 |
| Accs | 0.00035 | 6.008438 |

| | | |
|---|---|---|
| RP24-252B[illegible] | 0.00855 | 6.044864 |
| Hace1 | 0.02985 | 6.128101 |
| Zfp212 | 0.0326 | 6.134178 |
| Banp | 0.0475 | 6.149376 |
| Nup54 | 0.00685 | 6.20792 |
| Edc3 | 0.0047 | 6.230034 |
| Cd209b | 0.00165 | 6.353053 |
| Kif24 | 0.01035 | 6.415492 |
| Serping1 | 0.0164 | 6.496893 |
| Cyb5d2 | 0.01835 | 6.512403 |
| RP24-64D2 | 0.0221 | 6.534514 |
| Helb | 0.01785 | 6.595536 |
| Gipc1 | 0.02425 | 6.633315 |
| Eci1 | 0.0252 | 6.701435 |
| Tmem144 | 0.03125 | 6.812621 |
| Carf | 0.00965 | 6.842199 |
| Kctd3 | 0.0244 | 6.845709 |
| Zfp169 | 0.04485 | 6.858485 |
| Hhat | 0.0311 | 7.046613 |
| Mx2 | 0.0419 | 7.16755 |
| Ddx18 | 0.0083 | 7.188646 |
| Dcun1d3 | 0.0219 | 7.205957 |
| Slc35a1 | 0.00605 | 7.426642 |
| Atraid | 0.03095 | 7.427671 |
| Zbtb7b | 0.00485 | 7.476277 |
| Rit1 | 0.0061 | 7.557767 |
| Tor4a | 0.00535 | 7.711332 |
| RP23-313J1 | 0.04655 | 7.803439 |
| AC103620.[illegible] | 0.0018 | 7.819845 |
| AC124461.[illegible] | 6.00E-04 | 7.866653 |
| Zfp637 | 0.02985 | 7.978845 |
| Zfp747 | 0.0034 | 8.142264 |
| Syngap1 | 0.02795 | 8.326903 |
| Ralgps2 | 0.01195 | 8.3861 |
| Zdhhc23 | 0.00155 | 8.400936 |
| Aldh7a1 | 0.0134 | 8.401518 |
| Dhdds | 0.00525 | 8.579479 |
| Setd1a | 0.00595 | 8.77458 |
| Med12 | 5.00E-05 | 8.845723 |
| Nipa1 | 0.03075 | 8.936439 |
| Urb1 | 0.0077 | 8.937616 |
| Nmb | 0.04335 | 9.001341 |
| Zfp961 | 0.0019 | 9.086281 |
| Fbxo7 | 0.00345 | 9.105637 |
| RP23-95L9. | 0.03225 | 9.121746 |
| Ada | 0.0071 | 9.157857 |
| RP23-446P[illegible] | 0.01675 | 9.173994 |
| Rasgrp2 | 0.0048 | 9.26493 |
| Dnaaf5 | 0.0146 | 9.433937 |
| Zfp952 | 0.0017 | 9.557147 |
| Rffl | 0.0033 | 9.776581 |
| Cyp51 | 0.02675 | 9.857419 |
| RP23-293N | 0.00545 | 9.886638 |

| | | |
|---|---|---|
| Ttll1 | 0.0312 | 10.26997 |
| Tmem91 | 0.0046 | 10.46152 |
| Trim41 | 0.04645 | 10.50635 |
| Zfp764 | 0.0184 | 10.82361 |
| Aspa | 0.00015 | 11.26668 |
| Mfsd7a | 0.00675 | 11.27332 |
| Ankrd26 | 0.00095 | 11.31575 |
| Zfyve16 | 5.00E-05 | 11.43821 |
| St5 | 0.00315 | 11.67637 |
| Mdm1 | 0.0044 | 11.68892 |
| Map3k9 | 0.00365 | 11.72609 |
| Polh | 0.00755 | 11.72885 |
| Ralgps1 | 0.00485 | 11.82829 |
| Acads | 0.0206 | 11.87685 |
| Nedd4 | 0.00115 | 12.16614 |
| Chek2 | 6.00E-04 | 12.22913 |
| Exosc3 | 0.008 | 12.43513 |
| Pdk4 | 0.0015 | 12.46966 |
| Mccc2 | 0.02185 | 13.34225 |
| Surf6 | 0.00225 | 13.34872 |
| Tmem140 | 0.0156 | 14.89659 |
| Plag1 | 0.0041 | 15.51154 |
| Rabepk | 5.00E-05 | 16.05099 |
| Trip10 | 0.0393 | 16.58722 |
| RP23-357J7 | 0.01415 | 16.89343 |
| AC115748.1 | 0.03715 | 16.94115 |
| Sun1 | 5.00E-05 | 17.1245 |
| Arl15 | 0.01425 | 18.07658 |
| Rpgrip1 | 0.0026 | 19.78685 |
| RP23-281H | 0.04665 | 23.2734 |
| Dars2 | 0.003 | 32.96698 |
| RP24-180B | 0.0417 | 40.11111 |
| Zfp882 | 0.0041 | 40.38339 |
| Prg4 | 3.00E-04 | 45.33364 |

Supplementary Table 4: Genes within IPA pathways unique to LXRαWT in ATMs

| IL17A signaling | |
|---|---|
| **Gene** | **Fold Change** |
| Cxcl10 | 3.64 |
| Rela | 2.31 |
| Egfr | 2.21 |

| Interferon signaling | |
|---|---|
| **Gene** | **Fold Change** |
| Ifit3b | 5.2 |
| Rela | 2.31 |
| Ifitm2 | 1.98 |

| TREM1 pathway | |
|---|---|
| **Gene** | **Fold Change** |
| Tlr9 | 2.48 |
| Rela | 2.31 |
| Fcgr2b | 2.19 |

| TLR signaling | |
|---|---|
| **Gene** | **Fold Change** |
| Tlr9 | 2.48 |
| Rela | 2.31 |
| Ubc | 2.15 |

| Autophagy | |
|---|---|
| **Gene** | **Fold Change** |
| Ctsl | 0.46 |
| Ctsz | 0.44 |
| Ctsk | 0.19 |

Supplementary Table 4: Genes within IPA pathways unique to LXRα S196A in ATMs

| Lysine degradation | |
|---|---|
| **Gene** | **Fold Change** |
| Aldh7a1 | 8.40 |
| Aasdh | 4.87 |

| Mitosis inhibition | |
|---|---|
| **Gene** | **Fold Change** |
| Senp7 | 5.87 |
| Hdac1 | 4.27 |
| Smad4 | 4.11 |
| Sae1 | 3.50 |
| Senp1 | 2.86 |
| Sp1 | 1.94 |

| Sumoylation pathway | |
|---|---|
| **Gene** | **Fold Change** |
| Chek2 | 12.23 |
| Pole | 5.33 |
| Dbf4 | 4.2 |
| Csk19 | 3.85 |

| PPAR signaling | |
|---|---|
| **Gene** | **Fold Change** |
| Il1b | 0.46 |
| Rap2a | 0.37 |
| Pparg | 0.26 |
| Aip | 0.26 |
| Pdgfa | 0.13 |
| Hras | 0.13 |

| Glycogen biosynthesis | |
|---|---|
| **Gene** | **Fold Change** |
| Ugp2 | 0.26 |
| Gyg | 0.2 |

| PPAR signaling | |
|---|---|
| **Gene** | **Fold Change** |
| Il1b | 0.46 |
| Rap2a | 0.37 |
| Pik3cb | 0.31 |
| Csnk2a2 | 0.24 |
| Hspb7 | 0.18 |
| Hras | 0.13 |

| NFAT in immune resp. | |
|---|---|
| **Gene** | **Fold Change** |
| Gnas | 0.43 |
| Rap2a | 0.37 |
| Rcan1 | 0.34 |
| Pik3cb | 0.31 |
| Itpr1 | 0.28 |
| Csnk1g2 | 0.28 |
| Hras | 0.13 |

Supplementary Table 5: Genes common between plaque CD68+ cells and ATMs

| Common in LCM and unique in WT ATM RNAseq | | | | |
|---|---|---|---|---|
| | LCM RNAseq | | ATM: Unique in WT | |
| Gene | pvalue | FC | pvalue | FC |
| St18 | 2.9E-05 | 0.406 | 0.0386 | 0.115284 |
| Fat3 | 0.002633 | 0.500 | 0.00085 | 0.188205 |
| Myo1b | 0.00219 | 0.518 | 0.0378 | 6.613849 |
| Ctsc | 4.24E-06 | 0.525 | 0.0423 | 1.897868 |
| Cxcl10 | 0.00959 | 0.532 | 0.01115 | 3.637381 |
| Wdfy3 | 0.000597 | 0.559 | 0.02355 | 2.186055 |
| Slc28a2 | 0.031783 | 0.582 | 0.03445 | 2.442977 |
| Tlr9 | 0.034262 | 0.595 | 0.04505 | 2.479433 |
| Ndrg1 | 0.00992 | 0.629 | 0.02165 | 0.262498 |
| C1qc | 0.00134 | 0.655 | 0.04925 | 1.687848 |
| Nfkbiz | 0.046718 | 0.660 | 0.03225 | 1.773534 |
| Cd38 | 0.0131 | 0.674 | 0.0053 | 3.115362 |
| Ifi207 | 0.00502 | 0.678 | 0.0274 | 1.772002 |
| Card11 | 0.049001 | 0.693 | 0.0227 | 0.276787 |
| Mylip | 0.024274 | 0.747 | 0.03265 | 2.515195 |
| Abcg1 | 0.019788 | 0.763 | 0.0153 | 0.486351 |
| Ogdh | 0.004398 | 1.444 | 0.03955 | 2.552003 |
| Aldh6a1 | 0.03824 | 1.548 | 0.00695 | 4.971909 |
| Prelp | 0.000159 | 1.647 | 0.0373 | 4.609116 |
| Slit3 | 0.01186 | 1.670 | 0.0483 | 12.01754 |
| Tppp | 0.00469 | 2.085 | 0.03485 | 2.872944 |
| Fxyd1 | 0.0017 | 2.173 | 0.048 | 6.221144 |
| Lyve1 | 0.000637 | 2.445 | 0.00115 | 2.453481 |

| Common in LCM and unique in | | |
|---|---|---|
| | LCM RNAseq | |
| Gene | pvalue | FC |
| Ddx60 | 0.000379 | 0.486 |
| Slc9a9 | 0.002567 | 0.521 |
| Stap1 | 0.006121 | 0.525 |
| Acp5 | 0.001101 | 0.551 |
| Cd83 | 0.003568 | 0.559 |
| Hfe | 0.003073 | 0.563 |
| Sh2d1b1 | 0.031713 | 0.570 |
| Cd300ld | 0.000337 | 0.582 |
| Mdm1 | 0.049138 | 0.599 |
| Spic | 0.050479 | 0.599 |
| Nlrp1c-ps | 0.040659 | 0.603 |
| Iqgap2 | 0.000682 | 0.607 |
| Dclre1c | 0.022673 | 0.629 |
| Sh3bp1 | 0.050932 | 0.637 |
| Slfn8 | 0.019654 | 0.642 |
| Cd52 | 0.047017 | 0.646 |
| Dusp6 | 0.019471 | 0.664 |
| Evl | 0.004989 | 0.669 |
| Myo9a | 0.002191 | 0.678 |
| Inpp5d | 0.012735 | 0.688 |
| Abcc3 | 0.013843 | 0.693 |
| Stab1 | 0.007895 | 0.697 |
| Slc43a2 | 0.021923 | 0.697 |
| Dab2 | 0.026957 | 0.712 |
| Samhd1 | 0.021986 | 0.717 |
| Bbx | 0.031904 | 0.717 |
| Efhd2 | 0.023783 | 0.732 |
| Tanc2 | 0.031345 | 0.732 |
| Fmn1 | 0.027033 | 0.742 |
| Nedd4 | 0.022435 | 1.376 |
| Trak2 | 0.012202 | 1.424 |
| Serping1 | 0.032939 | 1.434 |
| Sort1 | 0.010882 | 1.454 |
| C3 | 0.002109 | 1.495 |
| Zbtb20 | 0.000665 | 1.516 |
| Narf | 0.044532 | 1.516 |
| Timm17a | 0.048512 | 1.537 |
| Bpgm | 0.035917 | 1.558 |
| Tfdp2 | 0.020786 | 1.613 |
| Fam220a | 0.044426 | 1.613 |
| Flrt2 | 0.027468 | 1.625 |
| Dlat | 0.013081 | 1.636 |

| | | |
|---|---|---|
| Eci1 | 0.04786 | 1.670 |
| Pink1 | 0.005881 | 1.682 |
| Homer1 | 0.006185 | 1.682 |
| Adam33 | 0.024157 | 1.682 |
| Etv1 | 0.033466 | 1.693 |
| Chchd3 | 0.006646 | 1.729 |
| Rgs5 | 0.005746 | 1.840 |
| Cped1 | 0.000527 | 1.879 |
| Atcayos | 0.006414 | 1.879 |
| Fosb | 0.014026 | 1.892 |
| Atp5c1 | 0.000202 | 1.959 |
| Acat1 | 0.000746 | 1.986 |
| Hspb7 | 0.008291 | 1.986 |
| Ndufa4 | 0.001554 | 2.042 |
| Pdk1 | 0.000336 | 2.235 |
| Idh2 | 3E-05 | 2.282 |
| Casz1 | 0.000887 | 2.378 |
| Slc25a4 | 0.00056 | 2.445 |

| S196A ATM RNAseq | |
|---|---|
| ATM: Unique in S196A | |
| pvalue | FC |
| 1.00E-04 | 3.701885 |
| 0.0486 | 1.984589 |
| 0.0116 | 0.307677 |
| 0.04345 | 0.091356 |
| 0.0497 | 0.567596 |
| 0.0147 | 3.174149 |
| 0.03925 | 0.187497 |
| 0.034 | 1.851085 |
| 0.0044 | 11.68892 |
| 0.01585 | 0.08922 |
| 0.0209 | 0.355362 |
| 0.0086 | 2.171512 |
| 0.0262 | 2.097644 |
| 0.01635 | 5.639081 |
| 0.0084 | 2.631217 |
| 0.00925 | 0.359004 |
| 0.00735 | 4.190348 |
| 0.00785 | 0.248711 |
| 0.04685 | 2.265705 |
| 0.0025 | 2.664546 |
| 0.0012 | 5.541014 |
| 0.02675 | 2.791809 |
| 0.04465 | 1.822309 |
| 0.0089 | 2.076552 |
| 0.01925 | 2.027511 |
| 0.00815 | 3.163584 |
| 0.0399 | 0.480664 |
| 0.0309 | 2.678379 |
| 0.01095 | 0.21076 |
| 0.00115 | 12.16614 |
| 0.00645 | 3.195895 |
| 0.0164 | 6.496893 |
| 0.0405 | 0.326142 |
| 0.03975 | 4.458072 |
| 3.00E-04 | 4.988652 |
| 0.03935 | 2.729866 |
| 0.0281 | 0.114094 |
| 0.0314 | 5.47341 |
| 0.02175 | 0.258402 |
| 0.0465 | 0.091289 |
| 0.03165 | 0.337748 |
| 0.03835 | 0.242395 |

| Common in LCM and WT/S196A ATM RNAseq | | | | | |
|---|---|---|---|---|---|
| | LCM RNAseq | | FB WT vs FBC WT | | FB WT vs |
| Gene | pvalue | FC | pvalue | FC | pvalue |
| Bank1 | 0.000537 | 0.406 | 0.00895 | 2.78431 | 0.00045 |
| Abca9 | 0.001987 | 0.543 | 0.0105 | 2.041203 | 0.02835 |
| Maf | 0.000944 | 0.547 | 5.00E-04 | 2.629922 | 0.00185 |
| Ear2 | 0.001904 | 0.603 | 4.00E-04 | 0.180829 | 0.00045 |
| Mmp12 | 0.000573 | 0.616 | 5.00E-05 | 0.083651 | 5.00E-05 |
| Cd180 | 0.004697 | 0.629 | 0.00595 | 0.404595 | 0.0149 |
| Bhlhe40 | 0.036479 | 1.338 | 0.00595 | 0.344955 | 0.008 |
| Egr1 | 0.025788 | 1.505 | 0.0011 | 2.29423 | 0.0019 |
| Fabp4 | 0.006305 | 1.613 | 0.04615 | 0.479938 | 0.00925 |

| | |
|---|---|
| 0.0252 | 6.701435 |
| 0.029 | 3.631486 |
| 0.0026 | 3.474872 |
| 0.0221 | 3.596915 |
| 0.0164 | 2.846443 |
| 0.014 | 0.259084 |
| 0.04305 | 2.869263 |
| 0.04545 | 4.180397 |
| 0.02625 | 5.925061 |
| 0.02945 | 1.843142 |
| 0.01845 | 0.35789 |
| 0.0297 | 0.258397 |
| 0.01875 | 0.18005 |
| 0.0443 | 0.32164 |
| 0.0335 | 0.297186 |
| 0.0185 | 3.197136 |
| 0.03915 | 3.93379 |
| 0.0319 | 0.324044 |

| |
|---|
| s FBC SA |
| FC |
| 4.371141 |
| 1.831151 |
| 2.506944 |
| 0.245083 |
| 0.05524 |
| 0.4943 |
| 0.406924 |
| 2.226457 |
| 0.323209 |

Gene in mitochondrial function pathway
